# Human ZKSCAN3 and *Drosophila* M1BP are functionally homologous transcription factors in autophagy regulation

**DOI:** 10.1101/747824

**Authors:** Marine Barthez, Mathilde Poplineau, Marwa Elrefaey, Nathalie Caruso, Yacine Graba, Andrew J. Saurin

## Abstract

Autophagy is an essential cellular process that maintains homeostasis by recycling damaged organelles and nutrients during development and cellular stress. ZKSCAN3 is the sole identified master transcriptional repressor of autophagy in humans. How ZKSCAN3 achieves autophagy repression at the mechanistic or organismal level however still remains to be elucidated. Here, we demonstrate that vertebrate ZKSCAN3 and Drosophila M1BP are functionally homologous transcription factors in autophagy repression. Expression of ZKSCAN3 in Drosophila prevents premature autophagy onset due to loss of M1BP function and conversely, M1BP expression in human cells can prevent starvation-induced autophagy due to loss of nuclear ZKSCAN3 function. In Drosophila ZKSCAN3 binds genome-wide to sequences targeted by M1BP and transcriptionally regulates the majority of M1BP-controlled genes. These data allow the potential for transitioning the mechanisms, gene targets and plethora metabolic processes controlled by M1BP onto ZKSCAN3 and opens up *Drosophila* as a tool in studying the function of ZKSCAN3 in autophagy and tumourigenesis.

## Introduction

Macroautophagy (hereafter simply termed autophagy) is an evolutionary conserved process in eukaryotes by which protein aggregates and damaged organelles are degraded by the lysosome and is implicated in several central biological processes including tissue remodelling during development, starvation adaptation, anti-aging, tumour suppression, and cell death (reviewed in (Mizushima, 2007)). Given its importance in cellular homeostasis and development, defects in autophagy and its regulation have been associated with numerous diseases such as cancer, neurodegenerative diseases (Parkinson) and diabetes (reviewed in Levine and Kroemer (2008)). Given the plethora pathophysiological processes linked to autophagy deregulation, it is not surprising that intense research has been directed into how normal autophagic processes are regulated.

The mammalian target of rapamycin (mTor) kinase is an important regulator of autophagy induction where active mTOR suppresses autophagy while inhibition of mTOR activity promotes autophagy onset (Rabinowitz and White, 2010), although a growing number of examples of mTOR-independent autophagy induction have been reported providing the notion that autophagy induction is not a single linear process but can rather be induced through the action of multiple interconnected key regulators (Corona Velazquez and Jackson, 2018). While autophagy is largely a cytoplasmic event, recent studies have focussed on the nuclear transcription factors and epigenetic marks that modulate the expression of various autophagic components (reviewed in Fullgrabe et al. (2014)). To date, more than 30 vertebrate transcription factors have been identified as positive transcriptional regulators of autophagy through activation of autophagy-related genes in vertebrates in response to various cellular stresses (reviewed in Pietrocola et al. (2013)). For example, under conditions of nutrient deprivation, phosphorylation of the MiTF transcription family member, TFEB by the mTor kinase in the cytoplasm results in its nuclear translocation and transcription of numerous autophagy-related genes (Settembre et al., 2013), a process that is evolutionary conserved in Drosophila (Bouche et al., 2016).

While numerous transcription factors activating autophagy have been identified and extensively studied, comparatively few are known to be clear inhibitors of autophagy induction. Nonetheless, master transcriptional repressors of autophagy have been identified in vertebrates and Drosophila. In Drosophila, the Hox family of transcription factors have been described as master transcriptional repressors of autophagy induction, where the clearance of all Hox proteins is required for either starvation or developmental autophagy induction (Banreti et al., 2014). Similarly, in vertebrates, the zinc finger with a SCAN and a KRAB domain 3 (ZKSCAN3) has been described as a master transcriptional repressor of autophagy (Chauhan et al., 2013), where its cytoplasmic relocation from the nucleus through JNK2-mediated phosphorylation is required for starvation or lysosomal stress-induced autophagy induction (Li et al., 2016).

If the transcription factors driving autophagy activation are mostly conserved throughout evolution (Chandra et al., 2016), it is less clear whether this is the case for transcription factors responsible for preventing autophagy induction. Given that the SCAN and KRAB domain are only found in vertebrate transcription factors, identifying a Drosophila homologue of ZKSCAN3 through similarity searches is not as straight forward as for other transcription factors. Nonetheless, Drosophila M1BP is a functional cofactor of Drosophila Hox proteins (Zouaz et al., 2017) and the presence of both M1BP and Hox are required for preventing autophagy induction in the Drosophila fat body (Banreti et al., 2014; Zouaz et al., 2017). Thus, while structural similarity between Drosophila M1BP and vertebrate ZKSCAN3 is restricted to their C-terminal C2H2 zinc finger domains, they are both required for autophagy inhibition. Moreover, the zinc finger associated domain (ZAD) of M1BP, while restricted to zinc finger proteins of dipteran and closely related insects, is analogous to the vertebrate KRAB domain, participating in a lineage-specific expansion of zinc finger proteins in insect and vertebrate genomes (Chung et al., 2002; Lespinet et al., 2002; Looman et al., 2002). Together, these functional and structural similarities led us to hypothesise that Drosophila M1BP and vertebrate ZKSCAN3 are functionally homologous proteins.

Here, we show that expression of vertebrate ZKSCAN3 in the Drosophila fat body prevents premature developmental autophagy induction caused by the loss of M1BP expression. Additionally, ZKSCAN3 binds the same genomic loci as M1BP in Drosophila cells and in the Drosophila fat body ZKSCAN3 transcriptionally controls two-thirds of M1BP-controlled genes. Similarly, we show that expression of M1BP in vertebrate cells is sufficient to prevent starvation-induced autophagy due to the cytoplasmic translocation of ZKSCAN3. Taken together, these data provide evidence that vertebrate ZKSCAN3 and Drosophila M1BP are functional homologues in the control of autophagy.

## Results and Discussion

### ZKSCAN3 expression in the Drosophila fat body rescues premature autophagy induced by M1BP loss-of-function

There are 23 vertebrate C2H2 zinc finger transcription factors containing SCAN and KRAB domains. Of these, ZKSCAN4 is the most similar to ZKSCAN3 in terms of sequence identity (Figure 1A,B). It has been proposed that the high sequence identity between ZKSCAN3 and ZKSCAN4 may result in functional redundancy, which could explain the absence of autophagy induction in ZKSCAN3-null mice (Pan et al., 2017). As both ZKSCAN3 and ZKSCAN4 share similar levels of identity to M1BP (Figure 1B), to study functional homology with Drosophila M1BP, we created independent myc-tagged ZKSCAN3 and ZKSCAN4 transgenic Drosophila fly lines under the expression control of the Gal4/UAS system (Duffy, 2002). Ubiquitous expression of either vertebrate gene using the ubiquitous Act5C-Gal4 driver had no apparent deleterious effects on general Drosophila health or longevity (Fig. 1C).

**Figure 1.**
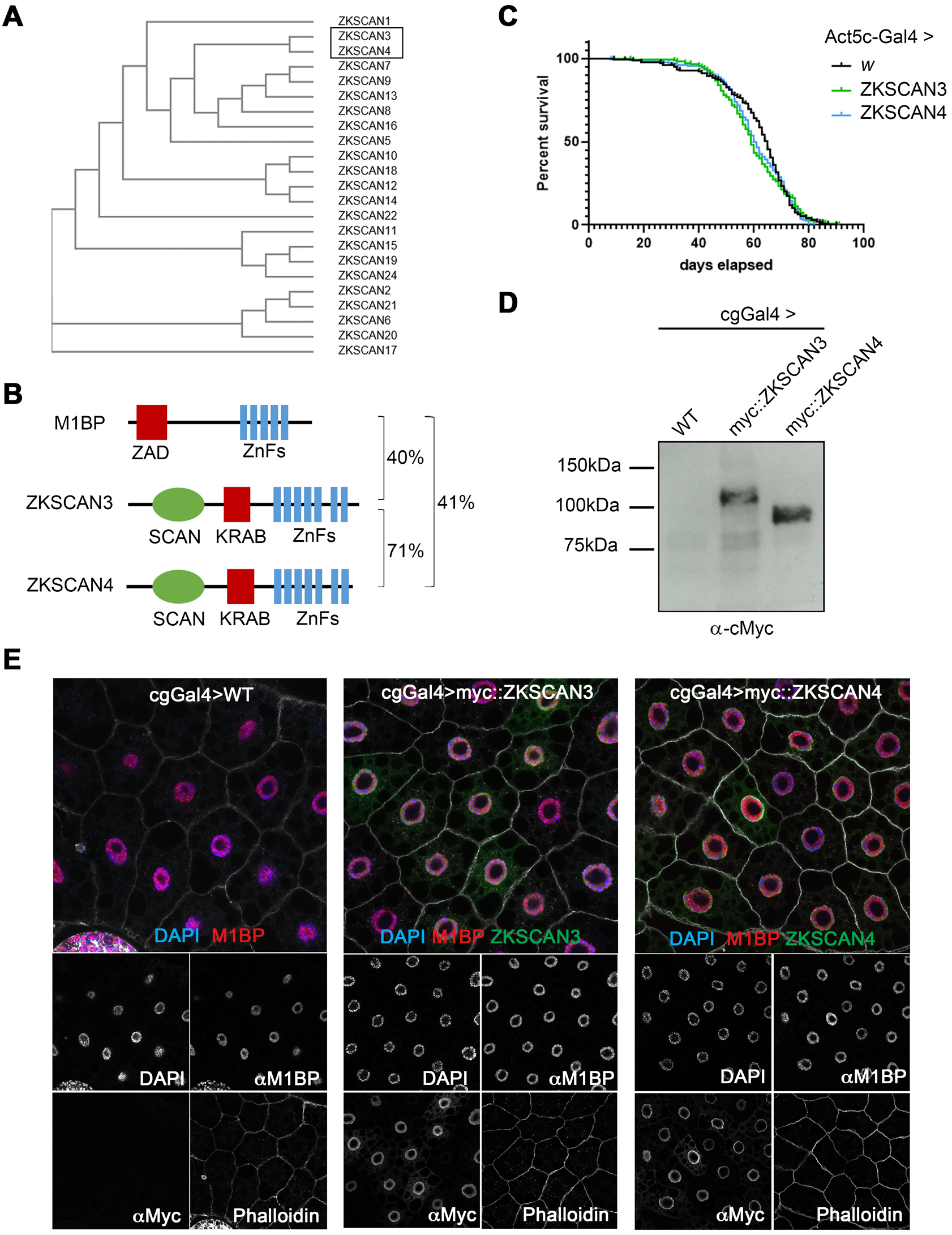
Using Drosophila to study ZKSCAN3 and ZKSCAN4 function in M1BP-controlled processes. **(A)** Phylogenetic tree analysis of primary sequence structure similarity of the vertebrate family of C2H2 zinc finger family transcription factor members containing a SCAN and KRAB domain demonstrates that ZKSCAN3 and ZKSCAN4 are paralogous family members. **(B)** The structural domains of Drosophila M1BP and vertebrate ZSKCAN3 and ZKSCAN4 are shown. C2H2 zinc finger domain clusters are depicted in blue, the SCAN domain, which is not conserved in Drosophila is depicted in green and the evolutionarily analogous ZAD and KRAB domains depicted in red. Percent sequence identity by BLAST conservation searches are shown. **(C)** Drosophila male lifespan was analysed when either ZKSCAN3 or ZKSCAN4 were ubiquitously expressed from the Act5CGal4 driver. No significant change to longevity was observed. **(D)** ZKSCAN3 and ZKSCAN4 expressed in the Drosophila fat body with the cgGal4 driver results in production of full-length protein as determined by western blot analysis. **(E)** Expression of ZKSCAN3 and ZKSCAN4 in the Drosophila fat body with the cgGal4 driver results in nuclear localised exogenous protein localisation (green channels) without changing nuclear M1BP staining (red channel). Nuclei were counterstained with DAPI (blue channel) and cell membranes revealed with Phalloidin (white). Scale bar represents 20 pm and contrasts of individual channels are shown.

The Drosophila fat body, a nutrient storage organ analogous to the vertebrate liver, is commonly used to study autophagy since autophagy in the Drosophila fat body is engaged as both an essential developmental process required for metamorphosis to give rise to the adult fly (Lorincz et al., 2017; McPhee and Baehrecke, 2009) and in response to nutrient deprivation under starvation conditions (Scott et al., 2004). When expressed in the Drosophila fat body using the cgGal4 driver, both ZKSCAN3 and ZKSCAN4 are expressed as full-length proteins (Fig. 1D) and, like in vertebrate cells (Li and al, 2007), both are mainly nuclear (Fig 1E), with weak accumulation observed in the cytoplasm (see Figure S1) with no apparent effects on fat body size or developmental progression (data not shown).

In the third instar larval feeding stage (L3F), autophagy is blocked because of the action of M1BP and Hox proteins (Banreti et al., 2014; Zouaz et al., 2017). Loss of expression of M1BP through RNA interference (RNAi) rapidly induces autophagy in the L3F stage (Zouaz et al, 2017 and see Fig. 2A). To test whether the co-expression of ZKSCAN3 or ZKSCAN4 is capable of preventing autophagy induction due to M1BP knockdown, we co-expressed either ZKSCAN3 or ZKSCAN4 with M1BP RNAi and monitored autophagy induction by cytoplasmic Atg8a accumulation, which is a marker of early autophagy induction, where its nuclear export into the cytoplasm is followed by lipidation recruiting it to early autophagosome membranes (Shpilka et al., 2011). Whether ubiquitously expressed in the fat body using the cgGal4 driver (Fig. 2A) or by clonal analyses (Fig. 2B), we found that the expression of ZKSCAN3, but not ZKSCAN4, rescues M1BP RNAi-induced autophagy as observed by the absence of cytoplasmic Atg8a (Fig. 2A,B). While it has been suggested that ZKSCAN4 may be able to compensate for ZKSCAN3 in ZKSCAN3-mediated autophagy repression (Pan et al., 2017), these data demonstrate that in Drosophila at least, ZKSCAN4 cannot replace ZKSCAN3 in autophagy repression induced by loss of M1BP function. M1BP RNAi induces increased Atg8a expression and lipidation of Atg8a, indicating increased autophagosome-associated Atg8a, and this is largely prevented upon ZKSCAN3 co-expression (Fig. 2C). ZKSCAN3 expression alone had no effect on normal expression (see Figure 2C) and nuclear localisation (Fig. S1A,B) of Atg8a suggesting that the autophagy rescue observed by ZKSCAN3 is not simply due to ectopic repression of Atg8a expression levels in the non-autophagic cellular state. Like Atg8a, Atg8b and Atg7 are also essential for autophagosome formation (Xiong, 2015). Similarly to Atg8a, we observed that co-expression of ZKSCAN3 restores expression of these early autophagosome markers that are prematurely induced in the L3F stage upon M1BP RNAi (Fig. 2D). Finally, by transmission electron microscopy of fat body cells, we observed large numbers of autophagosomes and late-stage autolysosomes due to M1BP RNAi, which were rarely observed in wild type cells or fat bodies co-expressing ZKSCAN3 (Fig. 2E).

**Figure 2.**
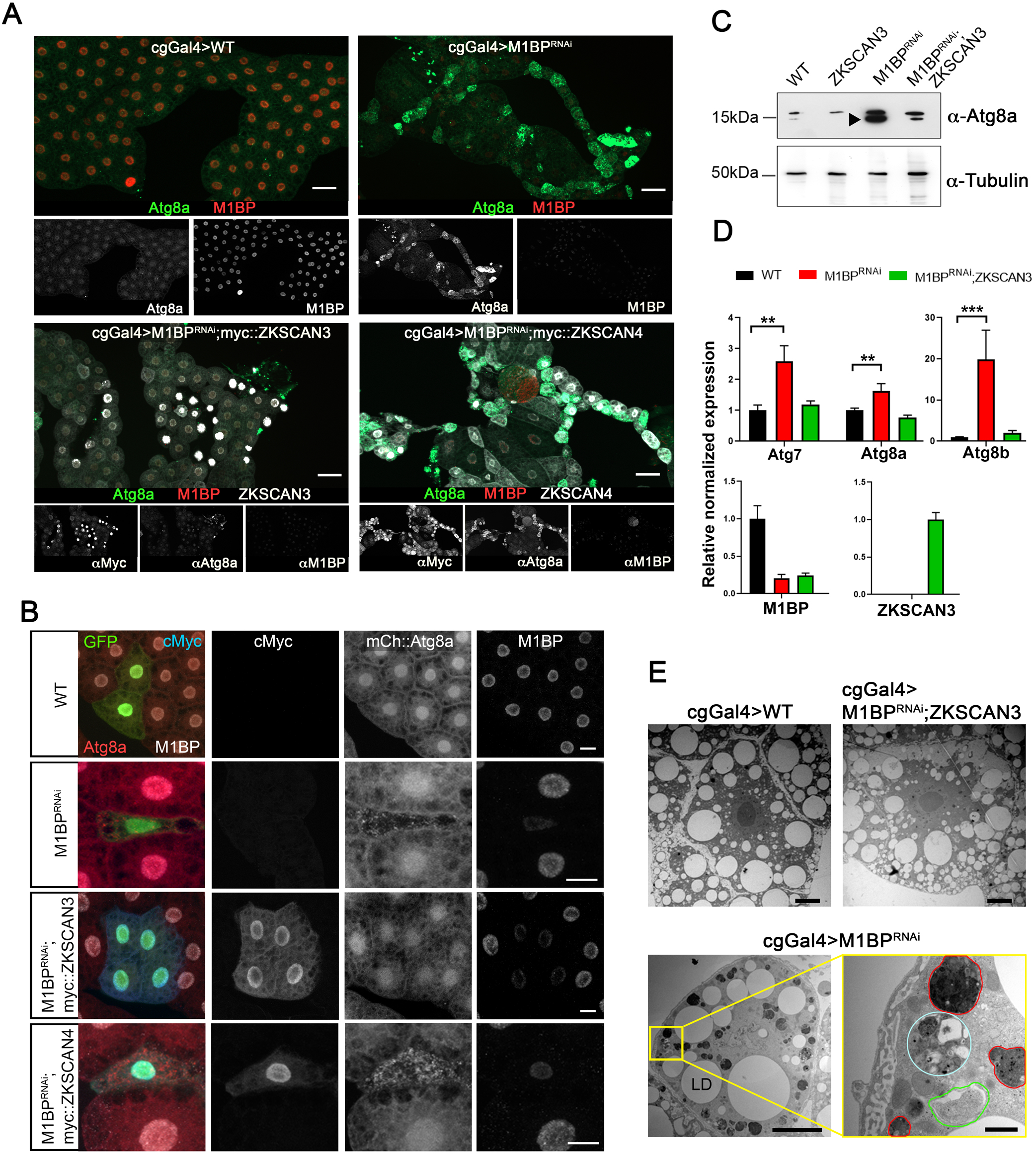
Expression of ZKSCAN3 in the Drosophila fat body prevents induction of autophagy due to loss of M1BP function. **(A)** Upon loss of M1BP expression (red) through expression of M1BP RNAi in L3F fat body cells using the fat body-specific cgGal4 driver, autophagy induction is widespread, as seen by the upregulation and cytoplasmic location of the Atg8a autophagy marker (green). Autophagy induction is largely prevented by the co-expression of myc-tagged vertebrate ZKSCAN3 (bottom left) but not by ZKSCAN4 (bottom right). Contrasts of individual channels are shown and scale bar represents 100 pm. **(B)** Clonal loss, RNAi-expressing cells are GFP-identified and autophagy monitored using mCherry*::Atg8a* confirming that co-expression of myc-tagged ZKSCAN3 can prevent autophagy induction through M1BP knockdown, whereas ZKSCAN4 expression does not. Contrasts of individual channels are shown and scale bars represent 100 pm. **(C)** Western blots of wild type, M1BP RNAi, ZKSCAN3, and M1BP RNAi;ZKSCAN3 expressing whole fat body protein preparations confirm autophagy induction due to M1BP RNAi, through the upregulation of Atg8a expression and the presence of major phosphatidylethanolamine-modified forms of Atg8a (arrowhead), which are largely prevented through ZKSCAN3 co-expression. The blots were reprobed with anti-tubulin antibodies for loading control. **(D)** RT-qPCR analyses confirm significant upregulation of autophagy-related Atg gene expression upon M1BP knockdown through M1BP RNAi in the fat body using the cgGal4 driver. Co-expression of ZKSCAN3 prevents Atg gene overexpression without modifying M1BP knockdown. **(E)** Transmission electron migrographs of Drosophila fat body cells display widespread autophagy induction upon M1BP RNAi through the presence of numerous autophagosome vesicles (lower panels) which are largely not observed in wild type cells or cells co-expressing ZKSCAN3 with M1BP RNAi (upper panels). Autophagosomes from all stages of maturity from early phagophore onset (green outline), autophagsomes containing large quantities of cellular material (cyan outline) and late-stage autophagosomes/autolysosomes (red outline) can be observed in M1BP RNAi expressing cells. LD: lipid droplets. Scale bars 10 pm in main migrographs and 1 pm in enlarged area (bottom right).

Together, these data demonstrate that the expression of vertebrate ZKSCAN3, but not the paralogous ZKSCAN4, in the Drosophila fat body is sufficient to prevent autophagy induction due to loss of M1BP expression, which we sought to study further.

### Loss of M1BP function leads to widespread gene deregulation that is largely restored upon ZKSCAN3 expression

To better understand the transcriptional changes occurring during loss of M1BP function, we performed RNA-seq analyses on fat body mRNA at the L3F stage in the absence and presence of M1BP RNAi. Differential gene expression analyses at a 1:1000 false discovery rate demonstrates that M1BP loss of function leads to the overexpression of 644 genes while down regulating 881 genes (Table S1). Gene set enrichment analyses of M1BP RNAi differentially expressed genes highlighted biological processes associated with responses to cellular nutrient levels, which include autophagy related genes, in genes that are overexpressed upon M1BP RNAi (Fig. S2A, left panel) and metabolic biological processes were enriched gene ontology terms associated with genes downregulated upon M1BP RNAi (Fig. S2A, right panel). Developmental autophagy induction during the L3 wandering stage (L3W) compared to L3F leads to the deregulation of 3738 genes (1819 upregulated; 1919 downregulated) (Table S1). Comparison of genes deregulated during developmental autophagy induction at the L3W stage with genes deregulated by M1BP RNAi at the L3F stage shows 916 commonly deregulated genes (Fig. S2B). These data demonstrate that M1BP RNAi in the L3F stage deregulates genes that are also deregulated during normal third instar larval development at the L3W stage when autophagy occurs, which suggests that M1BP RNAi in the L3F stage induces premature autophagy. Nonetheless, there are differences observed in genes deregulated upon M1BP RNAi compared to developmental autophagy (see (Fig. S2B) suggesting that M1BP performs other functions in the fat body than autophagy repression and that loss of M1BP function is not simply accelerating L3F development to that of an L3W stage.

Expression of ZKSCAN3 alone in the wild type fat body has little effect on gene expression, with very few genes classed as differentially expressed (Table S1). However, of the 1525 genes differentially expressed due to M1BP RNAi, more than two-thirds (1026) are restored to normal wild type L3F expression levels upon simultaneous co-expression of ZKSCAN3 (Table S1 and Fig. 3A), including a number of autophagy-related Atg genes (Fig. 3B). Similar to the gene set enrichment analysis of enriched gene ontology terms affected by M1BP RNAi (Fig. S2A), Reactome Pathway analysis (Fabregat et al., 2017) showed significant enrichment of the Autophagy Reactome for upregulated genes upon M1BP RNAi (30% of Autophagy Reactome entities) and significant enrichment of the Metabolism Reactome for M1BP RNAi-downregulated genes (23% of Metabolism Reactome entities) (Fig. 3C). The co-expression of ZKSCAN3 with M1BP RNAi restores expression of the majority of these Reactome pathway genes to wild type levels (Fig. 3C). Together, these data demonstrate that M1BP controls thousands of genes in the Drosophila fat body, the majority of which are normally deregulated during larval development and autophagy. Moreover, vertebrate ZKSCAN3 can transcriptionally control the majority of these genes, suggesting that ZKSCAN3 can bind the same genomic targets as M1BP.

**Figure 3.**
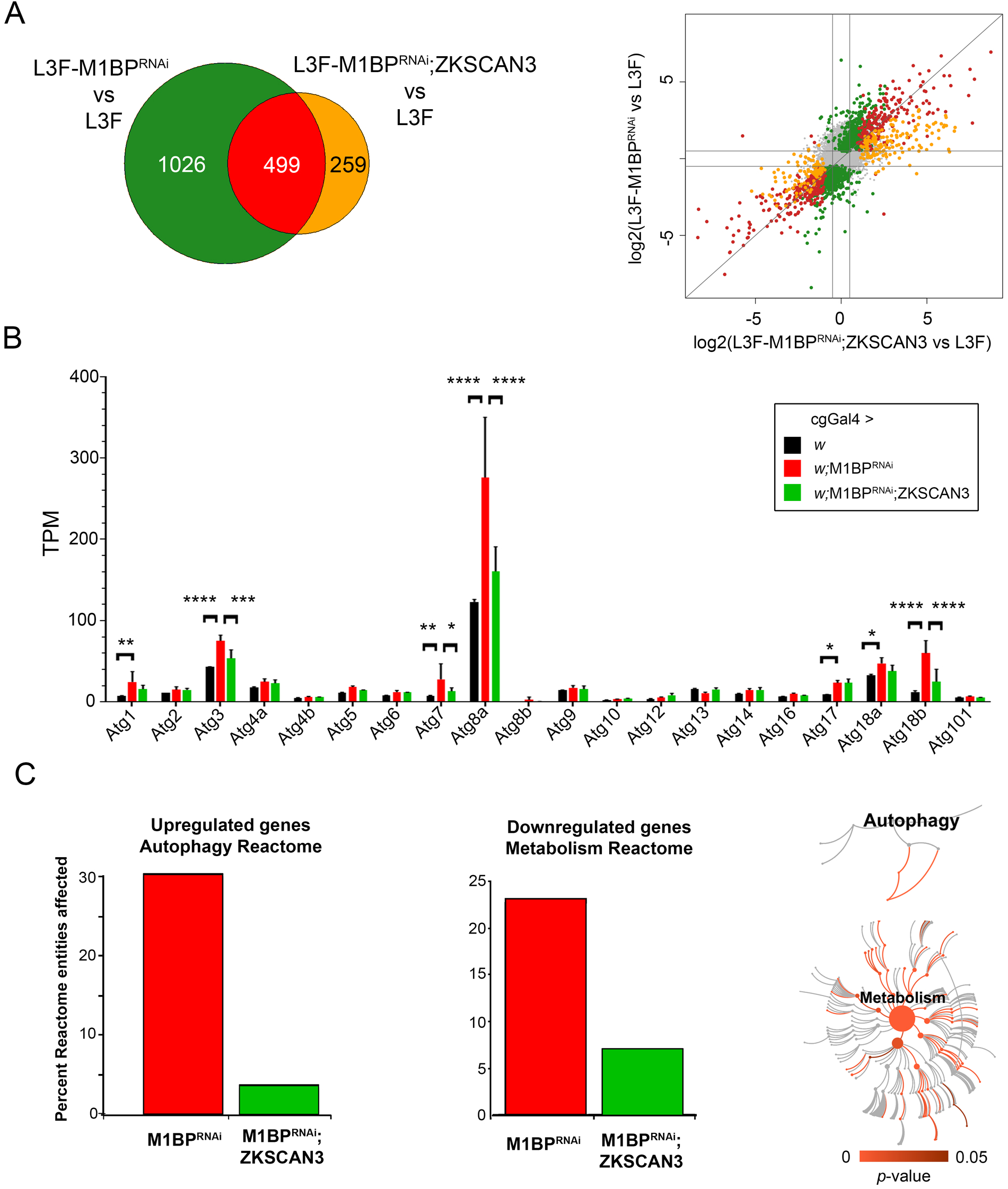
M1BP RNAi leads to widespread gene deregulation that is largely prevented through ZKSCAN3 co-expression. **(A)** Weighted Venn diagram representation of RNA-seq-identified L3F differentially expressed genes (DEGs) upon M1BP RNAi knockdown and DEGs in M1BP RNAi;ZKSCAN3 coexpressing fat body cells highlighting that more than two-thirds (1026) of M1BP RNAi-induced DEGs are prevented by ZKSCAN3 co-expression. Scatter plot of Iog2 fold changes in gene expression upon M1BP RNAi (ordinate) and M1BP RNAi;ZKSCAN3 (abscissa) coexpressing L3F fat body cells. Differential gene expression analyses identifying DEGs in both conditions are marked in red, genes prevented from being differentially expressed upon M1BP RNAi through ZKSCAN3 co-expression are shown in green and DEGs present in only M1BP RNAi;ZKSCAN3 fat bodies are shown in orange. Genes not differentially expressed in any condition are in grey. **(B)** Average transcripts per million (TPM) from biological replicate RNA-seq counts for all autophagy-related genes (Atg) show significant upregulation of numerous Atg genes upon M1BP RNAi that is prevented by ZKSCAN3 co-expression. Error bars represent standard deviation of mean replicate values. **(C)** Reactome pathway enrichment analysis (Fabregat et al., 2017) identified the “Autophagy Reactome” pathway as significantly enriched with 30% of pathway entities being upregulated upon M1BP RNAi and the “Metabolism Reactome” as significantly enriched with 23% of pathway entities being downregulated upon M1BP RNAi. In both cases, the majority of deregulated pathway genes are restored to wild type values upon ZKSCAN3 co-expression leading to the absence of pathway enrichment. The pathways significantly affected within each of the reactomes are shown in red while not significant pathways are shown in grey.

### ZKSCAN3 binds identical genomic sites as M1BP genome-wide

We sought to compare the genomic binding profiles of ZKSCAN3 and M1BP by genome-wide ChIP-seq analyses. However, using a range of regular chromatin preparation and sonication protocols and kits, we were unable to obtain adequate DNA fragment sizes following fixation and sonication for high-resolution ChIP-seq analyses (data not shown). We believe this is due to the heavy polytinised nature of chromosomes from the Drosophila fat body (256+ copies) (Richards, 1980), which may explain the lack of public whole-genome ChIP datasets from this tissue. Nevertheless, the binding profile of M1BP in Drosophila S2 cell culture cells has been extensively studied (Li and Gilmour, 2013; Zouaz et al., 2017). Thus to study whether ZKSCAN3 is capable of binding to similar genomic loci as M1BP, we established stable cell lines conditionally expressing HA-tagged ZKSCAN3 and as comparison, HA-tagged ZKSCAN4 (Fig. S3) and performed anti-HA ChIP-seq analyses. Remapping of previous M1BP S2 ChIP-seq datasets (Zouaz et al., 2017) to the BDGP Release 6 genome assembly (dm6) and peak calling with highly significant parameters (1% irreproducible discovery rate) identifies 6116 high confidence MlBP-bound genomic regions. Using identical stringent parameters, we identified 8991 ZKSCAN3 bound regions and 8747 ZKSCAN4 bound regions in S2 cells. Eliminating so-called high occupancy target (HOT) regions that appear indiscriminately bound by numerous transcription factors (Boyle et al., 2014), we identified 5279 M1BP peaks, 7884 ZKSCAN3 peaks, and 7734 ZKSCAN4 peaks. Since the DNA recognition sites of ZKSCAN3 and ZKSCAN4 *in vivo* have not been experimentally determined, we performed *de novo* motif discovery on ZKSCAN3 and ZKSCAN4 peaks and found that the most significant overrepresented motif at ZKSCAN3 genomic targets highly resembles the M1BP “Motif 1” binding motif, whereas ZKSCAN4 peaks are overrepresented with DNA binding motifs resembling that of GATA transcription factors (Fig. 4A). While ZKSCAN3 and ZKSCAN4 are similar in protein sequence identity (see Figure 1B), the DNA sequence motif overrepresentation in genomic binding sites is clearly distinct. Identifying the preferred binding sequence motif nonetheless affords an explanation as to how ZKSCAN3 can functionally replace M1BP in autophagy repression, through binding similar genomic sequences, and conversely explain the lack of autophagy rescue by ZKSCAN4.

**Figure 4.**
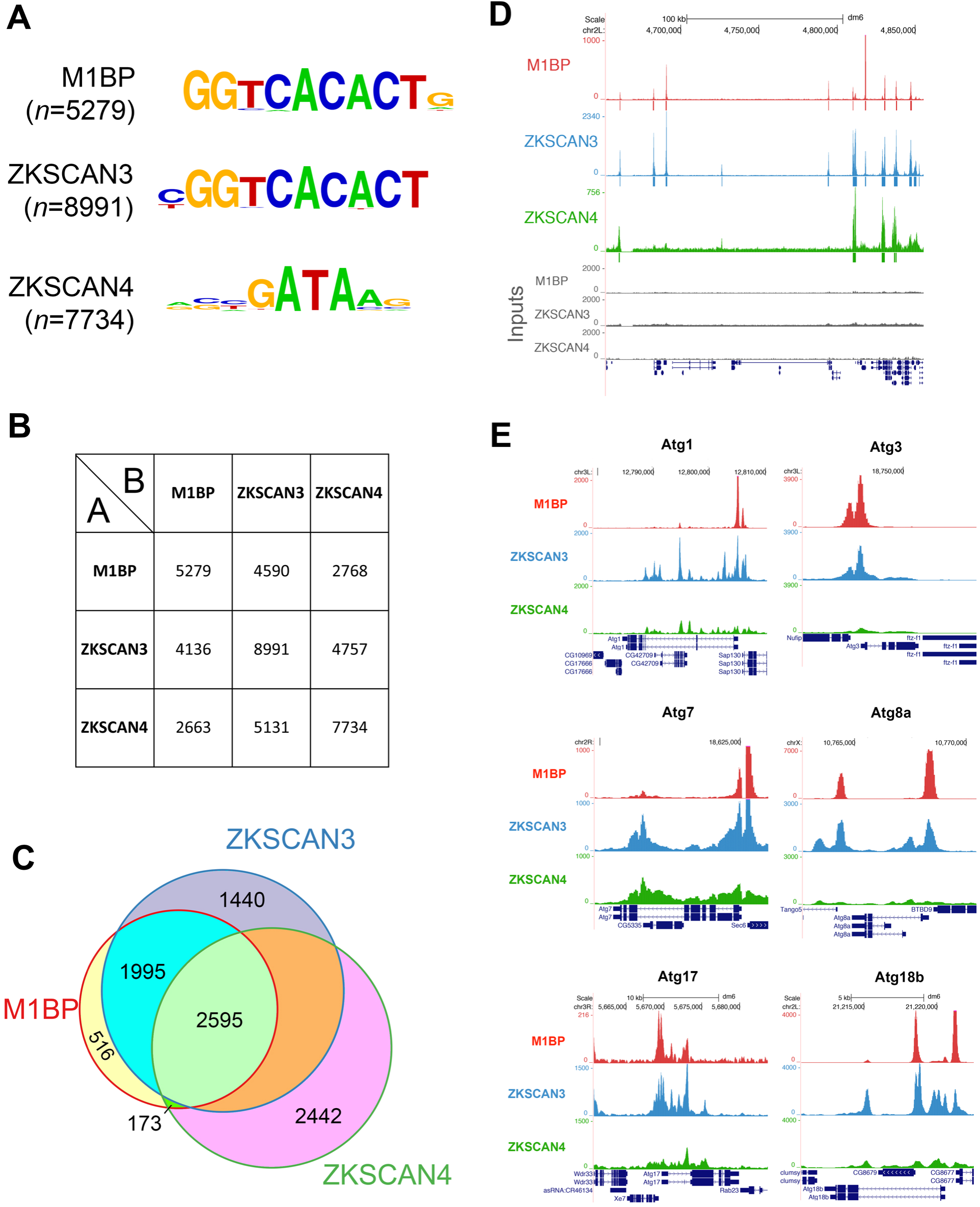
ZKSCAN3 binds identical M1BP genomic targets in Drosophila S2 cells. **(A)** The most significant DNA sequence motif found by *de novo* motif discovery analyses on high confidence called peaks (1% irreproducible discovery rate) of M1BP (*n*=5279 peaks), ZKSCAN3 (*n*=7884 peaks), and ZKSCAN4 (*n*=7734 peaks) ChIP-seq profiles shows that Motif 1 DNA element (Ohler et al., 2002) is significantly enriched at ZKSCAN3 and M1BP peaks. **(B)** Pairwise peak intersection analyses is shown with of the number of peaks in the “A” peak datasets (vertical dataset identity) overlapping with peaks of the “B” peak dataset (horizontal dataset identity) displayed. **(C)** Weighted Venn diagram representation of the number of M1BP peaks overlapping with ZKSCAN3, ZKSCAN4 or both ZKSCAN3 and ZKSCAN4 peaks are shown. Since the numbers of peaks change slightly depending on which dataset is used as the source identity (see “A” versus “B” and vice versa in (B) above), only the numbers of peaks not overlapping with any other dataset can be shown for ZKSCAN3 and ZKSCAN4. **(D)** Representative genome browser view of M1BP, ZKSCAN3, and ZKSCAN4 ChIP-seq profiles when expressed in Drosophila S2 cells. Sequenced input DNA of each condition is shown in grey. **(E)** Genome browser profiles demonstrating that M1BP and ZKSCAN3 target similar autophagy related gene (Atg) promoters in S2 cells.

Comparing the genomic binding profiles of M1BP with ZKSCAN3 and ZKSCAN4, we observed that of the 5279 HOT-excluded genomic regions bound by M1BP (Zouaz et al., 2017), 87% (4590) are identically targeted by ZKSCAN3 and 52% (2768) are targeted by ZKSCAN4 (Fig. 4B). Surprisingly, of the 2768 M1BP target loci that are also targeted by ZKSCAN4, almost all (94%; 2595 peaks) are also targeted by ZKSCAN3 (Fig. 4C). These data demonstrate that while ZKSCAN4 shares some common M1BP genomic loci, the vast majority of M1BP loci are also loci targeted by ZKSCAN3 (for representative example, see Figure 4D). Interestingly, of the numerous Atg autophagy genes deregulated in the Drosophila fat body upon M1BP RNAi and rescued by ZKSCAN3 co-expression (Fig. 3B), many of their promoters are also M1BP targets and these are shared targets of ZKSCAN3 (Fig. 4E). These data suggest that M1BP and ZKSCAN3 are likely direct repressors of autophagy-related gene expression, whereby loss of M1BP/ZKSCAN3 function leads to Atg upregulation, which is essential for both autophagy induction and progression (Yu et al., 2018). Thus, given that ZKSCAN3 can target identical genomic regions as M1BP in Drosophila cells and can restore the expression of the majority of genes deregulated through M1BP RNAi, these data strongly suggest that ZKSCAN3 replaces M1BP function in autophagy repression through similar transcriptional mechanisms employed by M1BP and not simply as an alternative repressor of autophagy.

### M1BP can repress autophagy induction due to cytoplasmic ZKSCAN3 translocation in vertebrates

In vertebrates, autophagy is induced under lysosomal stress and starvation conditions due to the relocalisation of ZKSCAN3 from the nucleus to cytoplasm (Chauhan et al., 2013). To further investigate the interchangeable nature of M1BP and ZKSCAN3 function in the control of autophagy, we asked whether Drosophila M1BP could functionally replace ZKSCAN3 in autophagy repression under starvation conditions in vertebrates. Using LC3A as an autophagy marker, under starvation conditions, autophagy is induced in vertebrate cells through the relocation of ZKSCAN3 from the nucleus to the cytoplasm (Fig. 5A and Chauhan et al. (2013)). We found that transfected Drosophila M1BP is nuclear, even under starvation conditions (Fig 5B). This nuclear localisation of M1BP was sufficient to significantly inhibit autophagy induction, where cytoplasmic LC3A accumulation is inhibited (Fig 5C) and autophagy related gene expression is restored (Fig. 5D). These data suggest starvation-induced autophagy due to the cytoplasmic relocation of ZKSCAN3 can be prevented through the function of Drosophila M1BP in the nucleus.

**Figure 5.**
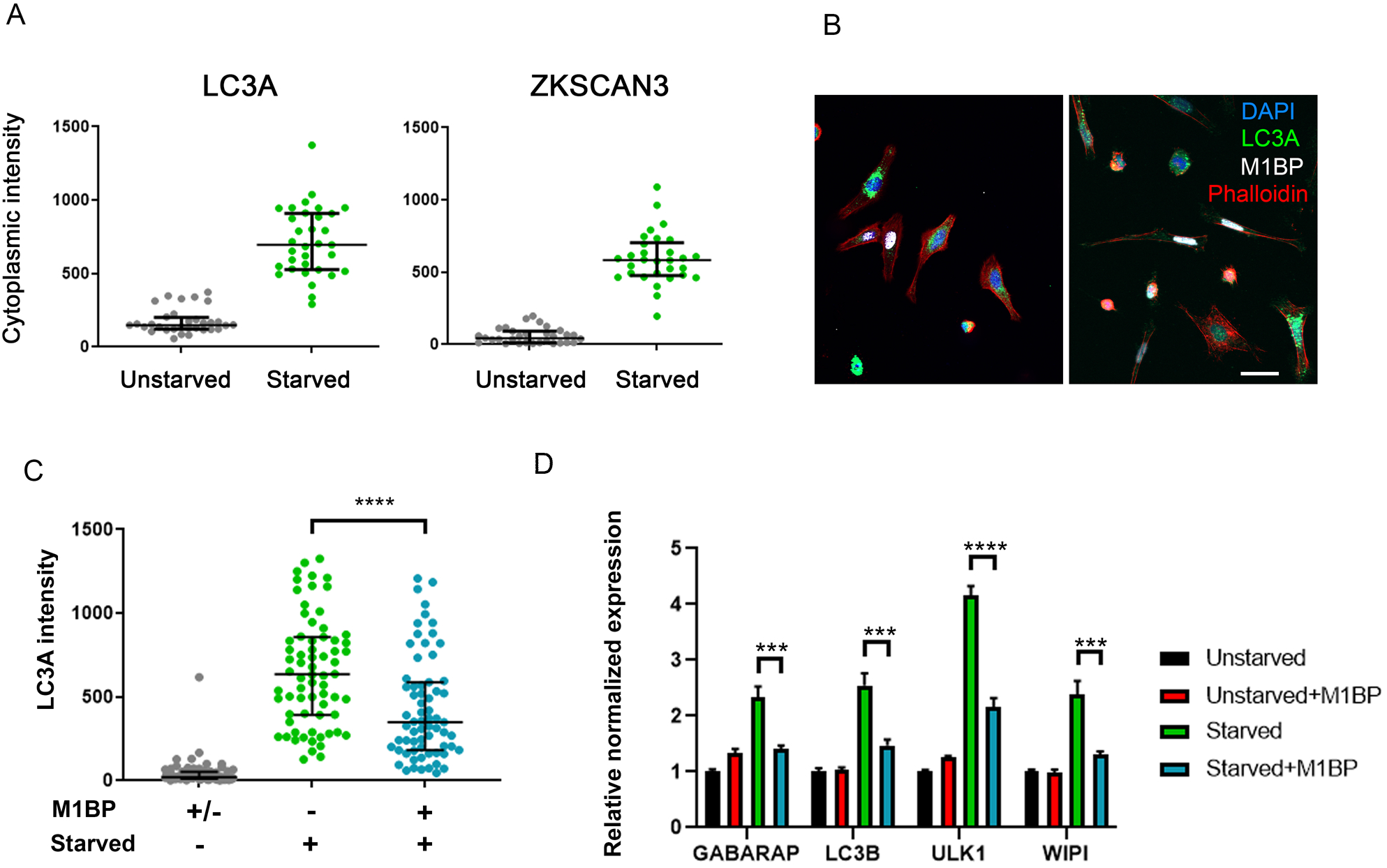
M1BP expression can prevent starvation-induced autophagy when expressed in vertebrate cells. **(A)** Quantification of cytoplasmic immunofluorescence signal of LC3A and ZKSCAN3 in unstarved and starved HeLa cells confirming autophagy induction through the shuttling of ZKSCAN3 into the cytoplasm. **(B)** Representative immunofluorescence staining of LC3A (green) in independent experiments of starved HeLa cells demonstrates that cells containing transiently transfected M1BP (white) display generally lower levels of cytoplasmic LC3A accumulation. Nuclei are counterstained with DAPI (blue) and cell membranes marked with Phalloidin (red). **(C)** Quantification of cytoplasmic LC3A immunofluorescence signal in unstarved and starved HeLa cells demonstrates significant reduction of cytoplasmic LC3A in cells transiently transfected with M1BP. **(D)** RT-qPCR analyses of key vertebrate autophagy genes induced upon HeLa cell starvation demonstrate induction of expression is significantly inhibited in cells transiently transfected with M1BP.

Collectively, these data demonstrate that vertebrate ZKSCAN3 and Drosophila M1BP are homologous transcription factors that bind similar sequences in Drosophila and can functionally replace each other in autophagy repression. In the regulation of autophagy, it is worth noting that in order to induce autophagy, the cellular mechanisms employed by Drosophila and vertebrates would appear to have evolved: In Drosophila, the nuclear function of both M1BP and Hox transcription factors are mutually required to prevent autophagy induction (Zouaz et al., 2017) whereby Drosophila clear all Hox function to induce autophagy during development or under starvation conditions (Banreti et al., 2014); in vertebrates, while the underlying reasons remain unknown, the mechanisms of autophagy induction have evolved to require the nuclear eviction of ZKSCAN3 by JNK2/p38-mediated phosphorylation of Thr153 within a nuclear export signal sequence (Chauhan et al., 2013; Li et al., 2016), a sequence which is not conserved in Drosophila M1BP (data not shown).

### Insights into regulated M1BP and ZKSCAN3 function

M1BP is known to bind promoters of genes with gene ontology terms linked to metabolism (Li and Gilmour, 2013; Zouaz et al., 2017) and our data show that metabolic gene expression is lost upon M1BP knockdown (Fig. S2A and 3C) suggesting that M1BP promoter binding to these genes actively controls their expression. Thus, since loss of M1BP expression leads to loss of metabolic gene expression and that changes in cellular metabolism are well known to induce autophagy (Galluzzi et al., 2014; Rabinowitz and White, 2010), this could be the cellular signal that leads to autophagy induction upon loss of M1BP function. Given that ZKSCAN3 transcriptionally rescues defects in metabolic gene transcription due to M1BP RNAi (Fig. 3C), this also suggests that, like M1BP, ZKSCAN3 regulates metabolic gene expression. From a mechanistic point of view, the sharing of promoter biding and gene regulation by M1BP and ZKSCAN3 also suggest that both molecules control gene expression though similar molecular mechanism. As M1BP controls transcription through RNA Pol II pausing (Li and Gilmour, 2013), ZKSCAN3 may also control this process, a hypothesis that remains to be tested.

Finally, as ZKSCAN3 dysregulation contributes to numerous human cancers (Chi et al., 2018; Kawahara et al., 2016; Kim et al., 2016; Lee et al., 2018; Li et al., 2019; Yang et al., 2008; Yang et al., 2011) and that metabolic reprogramming is a hallmark of the tumour microenvironment (reviewed in (Reina-Campos et al., 2017)), the knowledge that Drosophila M1BP and vertebrate ZKSCAN3 are functionally homologous proteins controlling autophagy and metabolic gene expression will allow use of the powerful Drosophila model system to study ZKSCAN3 function in metabolism, autophagy control and cancerogenesis.

## Acknowledgements

We thank T. Neufeld and the Bloomington and Vienna Stock Centers for providing Drosophila lines, D. Gilmour and G. Juhász for their generous gifts of antibodies, and the IBDM Electron Microscopy and Imaging Services. MP acknowledges support from the Fondation de France. This work was supported by the Fondation pour la Recherche Médicale, Association pour la Recherche sur le Cancer, and la Ligue Nationale Contre le Cancer.

## Materials and Methods

### Fly stocks

The following lines UAS-3xmyc::ZKSCAN3 and UAS-3xmyc::ZKSCAN4 were established by using a pUAST plasmid encoding 3xmyc epitope tag (p131) and cDNA of vertebrate ZKSCAN3 variant 1 (NM_001145778.1) and ZKSCAN4 (IMAGE clone 4336904). For M1BP loss of function the line UAS-M1BP^RNAi^ 30016kk was obtained from the Vienna Drosophila Stock center. The fat body specific cgGal4 (Bloomington 7011) and ubiquitous Act5C-Gal4 (Bloomington 3954) driver lines was obtained from the Bloomington stock centre. For clonal analysis, the yw,hs-Flp;r4-mCherry::Atg8a;Act>CD2>GAL4,UAS-GFPnls fly stock was used (a gift from T.Neufeld).

### Imaging

For immunofluorescences, around 10 larval fat bodies are dissected per condition in 0.01M PBS/0,7% NaCl, fixed for 1h in 1.8% formaldehyde in PBS and washed for 10 min in PBS followed by 15 min in PBTX-DOC (PBS, 0.1% Triton X-100, 0.05% DOC). After 1H of blocking in 1% BSA/PBTX-DOC, fat bodies were incubated overnight with primary antibodies at 4°C. Following 3×30 min washes with PBTX-DOC, secondary antibodies were added for 1H. After washing for 1H in PBTX-DOC followed by 15 min in PBS, fat bodies were mounted in Vectashield and visualised using confocal (Zeiss LSM 880) or apotome (Zeiss AxioImager APO M2) fluorescence microscopy.

Transmission electron microscopy of fat bodies from larvae of cg-Gal4>UAS-M1BP^RNAi^, cg-Gal4>UAS-M1BP^RNAi^;ZKSCAN3 and control cg-Gal4>+ were dissected, fixed and processed as described in (Banreti et al., 2014).

### Antibodies

The following primary antibodies were used: rat anti-atg8a (a gift from G. Jusász), rabbit anti-M1BP (Zouaz et al., 2017), mouse anti-c-Myc (9E10; Santa Cruz Biotech; SC-40), mouse anti-His (Abcam; ab9136) and rabbit anti-LC3A (Cell Signaling Techology).

### Quantitative RT-PCR

Total RNA was isolated from 10 fat bodies of males (gonads removed) in L3F with the RNeasy Mini Kit (Qiagen) and either sent for RNA-seq, after verification of the RNA quality on Bioanalyser (Agilent), or the RNA was reversed transcribed to cDNA by using the SuperScript III according to the manufacturer’s instructions (Life Technologies). qRT-PCR was carried out with SYBR green mix (Life Technologies) and performed in triplicate on CFX96 thermocycler (BioRad). Data and statistical tests were analysed on CFX manager software (BioRad).

The following Drosophila-specific primers were used for qRT-PCR on fat bodies:

~~~
M1BP F: CGCATGGCCTTTGAACTT
M1BP R: GAAGCGCGACTGACAGAGTT
Atg8a F: GGGAGCCTTCTCGACGAT
Atg8a R: TTCATTGCAATCATGAAGTTCC
Atg8b F: CGGTGGGATCACATTGTTTA
Atg8b R: ATCCGCAAGCGTATCAATCT
Atg7 F: GGCTGTCATCGATGTTCATTT
Atg7 R: TTTCTGCTTCAGCAATGTCC
ZKSCAN3 F: AGCAGGATTCATCTCAGGGGA
ZKSCAN3 R: ACTCTTTCCACATTCATGGCAGA
ZKSCAN4 F: GGTGGTGGTGCTATTGGAGT
ZKSCAN4 R: GTCACCAACGGGAACCTG
U6 F: CGATACAGAGAAGATTAGCATGGC
U6 R: GATTTTGCGTGTCATCCTTGC
Rp49 F: CGCTTCAAGGGACAGTATCTG
RP49 R: AAACGCGGTTCTGCATGA
Actin5c F: TCAGTCGGTTTATTCCAGTCATTCC
Actin5c R: CCAGAGCAGCAACTTCTTCGTCA
~~~

### ChIP-seq experiments

S2-DRSC cells (Drosophila Genomics Resource Center, stock #181) were cultured at 25°C in Schneider’s medium (Clinisciences) supplemented with 10% FBS. HA-tagged ZKSCAN3 or ZKSCAN4 were cloned into pMK33/pMtHy (Drosophila Genomics Resource Center), transfected into S2-DRSC cells and stable lines generated through Hygromycin B selection over a 4-week period. ZKSCAN3 or ZKSCAN4 were induced with 100 μm CuSO_4_ addition to the media for 48 h before chromatin preparations and anti-HA ChIP performed as previously described (Zouaz et al., 2017). Anti-HA ChIP-seq on untransfected and CuSO_4_-induced S2 cells has previously been performed yielding no significant anti-HA ChIP peaks (Zouaz et al., 2017).

### High-throughput sequencing and analysis

Sequencing was performed on libraries prepared from duplicates of ChIP or RNA preparations. RNA-seq libraries were prepared using the TruSeq RNA Sample Preparation Kit (Illumina) and ChIP-seq libraries were prepared using theMicroPlex Library Preparation Kit (Diagenode) following the manufacturer’s instructions. All libraries were validated for concentration and fragment size using Agilent DNA1000 chips. Sequencing was performed on a HiSeq 4000 or Next-seq500 sequencer (Illumina) using a 150-nt paired-end protocol. Base calling performed using RTA (Illumina) and quality control performed using FastQC.

M1BP S2 ChIP-seq sequences were previously generated (GSE101557) and remapped to the dm6 UCSC Drosophila genome release using Subread Aligner (Liao et al., 2013) (version 1.6.1) resulting in an average of 14.1 × 10^6^ mapped reads from both replicates. ZKSCAN3 and ZKSCAN4 ChIP-seq sequences were mapped to the dm6 UCSC Drosophila genome release resulting in an average of 42.8 × 10^6^ ZKSCAN3 and 35.6 × 10^6^ ZKSCAN4 mapped reads per replicate. High confidence binding sites were determined through peak calling using MACS2 (version 2.1.1.20160309) using an irreproducible discovery rate (IDR) of 1% for all ChIP-seq data as outlined by the ENCODE and modENCODE consortium using each biological replicate. IDR peaks localising to HOT regions were removed resulting in 5279 M1BP peaks, 7884 ZKSCAN3 peaks and 7734 ZKSCAN4 peaks. Comparison of binding of ZKSCAN3 and ZKSCAN4 to M1BP genomic targets was performed using BEDTools (Quinlan and Hall, 2010) “intersect” using M1BP peaks as the *“A”* dataset and ZKSCAN3 or ZKSCAN4 peaks as the “B” dataset to allow direct comparison of co-targeted regions.

*De novo* motif discovery was performed on all high confidence M1BP, ZKSCAN3, and ZKSCAN4 peak region centers in S2 cells using Homer. Similarity matching of identified motifs with known motifs was performed using STAMP and the most significant overrepresented motif shown.

For RNA-seq analyses, raw counts from RNA-seq were obtained using featureCounts from the Subread package (Liao et al., 2013) (Version 1.6.1). Differentially expressed genes were called using DESeq2 (Love et al., 2014) (version 1.14.1) using a false discovery rate (p-adjusted value in DESeq2) threshold of 0.001.

### Vertebrate Cell culture

HeLa cell line was cultured at 37°C with 5% CO_2_ in Dubelcco’s modified Eagle’s medium (DMEM) supplemented with 1% FBS from ATCC. Transient transfection was performed with X-tremeGENE™ HP DNA Transfection Reagent (Sigma-Aldrich) following the manufacturer’s instructions. For starvation, cells were cultured for 4H in DMEM without amino acid and with a low concentration of glucose as previously described (Banreti et al., 2014).

For qRT-PCR, 0.5×10^6^ cells were seeded and extracted as described above. The following primers were used:

~~~
M1BP F: CGCATGGCCTTTGAACTT
M1BP R: GAAGCGCGACTGACAGAGTT
LC3B F: CGCACCTTCGAACAAAGAG
LC3B R: CTCACCCTTGTATCGTTCTATTATCA
ULK1 F: CAGACGACTTCGTCATGGTC
ULK1 R: AGCTCCCACTGCACATCAG
GABARAP F: CGGGTGCCGGTGATAGTAGA
GABARAP R: TGAGATCAGAAGGCACCAGGTA
WIPI F: TCAAACTCGAGACTGTGAAAGAAA
WIPI R: AGCACTTTCCCGAAGTACCC
ZKSCAN3 F: TTCATCTCAGGGGAATCTCTG
ZKSCAN3 R: GAGGCAAGTCCCTGCTCTTA
GAPDH F: GTCAAGGCTGAGAACGGGAA
GAPDH R: AAATGAGCCCCAGCCTTCTC
B-actin F: AGAGCTACGAGCTGCCTGAC
B-actin R: AGCACTGTGTTGGCGTACAG
RNA18S F: CCGATTGGATGGTTTAGTGAG
RNA18S R: AGTTCGACCGTCTTCTCAGC
~~~

