## Supplementary Figures 1-3 for "Human ZKSCAN3 and *Drosophila* M1BP are functionally homologous transcription factors in autophagy regulation"

### Figure S1

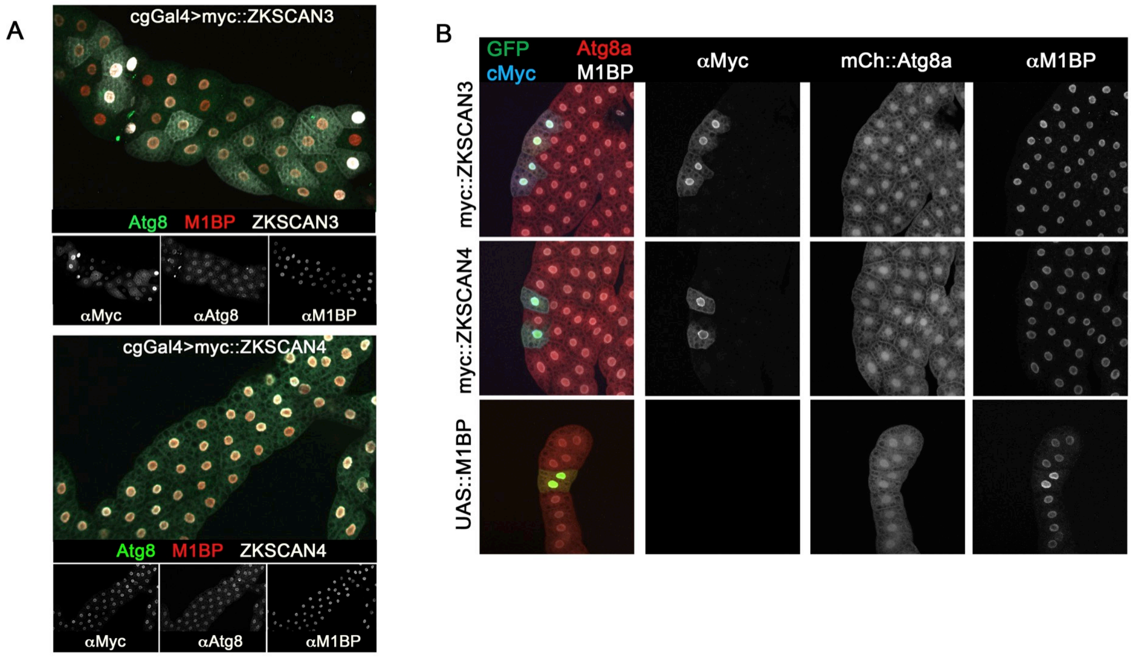


**Figure S1. Expression of ZKSCAN3 and ZKSCAN4 in the Drosophila fat body does not affect normal M1BP or Atg8a expression and localisation.**

**(A)** myc::ZKSCAN3 (upper panels) or myc::ZKSCAN4 (lower panels) transgenes were expressed in the Drosophila fat body using the cgGal4 driver and detected with anti-myc tag antibodies (white staining). Atg8 (green channel) and M1BP (red channel) were detected with respective antibody immunolabelling. Scale bar represents 100 µm and contrasts of individual channels are shown. **(B)** myc::ZKSCAN3 (upper panels), myc::ZKSCAN4 (center panels), or M1BP transgenes were expressed clonally in the Drosophila fat body expressing mCherry-fused Atg8a. Clones are labelled with GFP, Atg8a expression shown with mCherry and clonally expressed transgenes detected by anti-myc tag (ZKSCAN3 and ZKSCAN4) or anti-M1BP immunolabelling. Scale bar represents 50 µm and contrasts of individual channels are shown.

### Figure S2

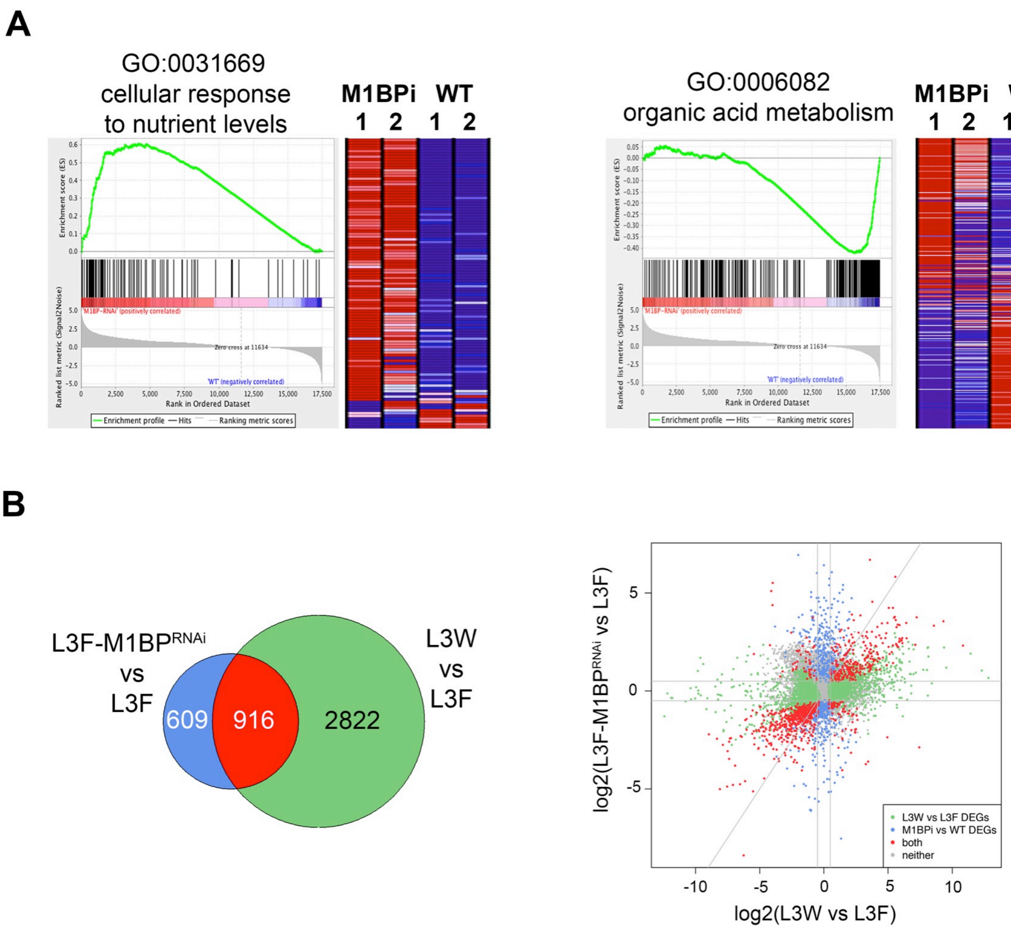


**Figure S2. M1BP RNAi deregulates genes with nutrient and metabolic gene ontologies and genes deregulated during developmental autophagy induction**

**(A)** Gene set enrichment analysis (GSEA) of M1BP RNAi and wild type gene expression profiles identified strong M1BP RNAi positive enrichment (ie upregulated by M1BP RNAi) of genes linked to cellular response to nutrient levels (GO term 0031669) which includes autophagy-related genes, whereas genes downregulated by M1BP RNAi that are enriched in WT samples include genes linked to metabolic gene ontologies (shown GO term 0006082). **(B)** Weighted Venn diagram representation of RNA-seq-identified L3F differentially expressed genes (DEGs) upon M1BP RNAi knockdown and DEGs during developmental autophagy in wandering (L3W) versus L3F fat body cells highlighting that the majority (916) of M1BP RNAi-induced DEGs are also deregulated during L3W-induced developmental autophagy. Scatter plot of log2 fold changes in gene expression upon M1BP RNAi (ordinate) and L3W (abscissa) versus L3F fat body cells. Differential gene expression analyses identifying DEGs in both conditions (*n*=916) are marked in red, genes only differentially expressed in L3W (*n*=2822) are shown in green and DEGs present in only M1BP RNAi fat bodies (*n*=609) are shown in blue. Genes not differentially expressed in any condition are in grey.

Figure S3

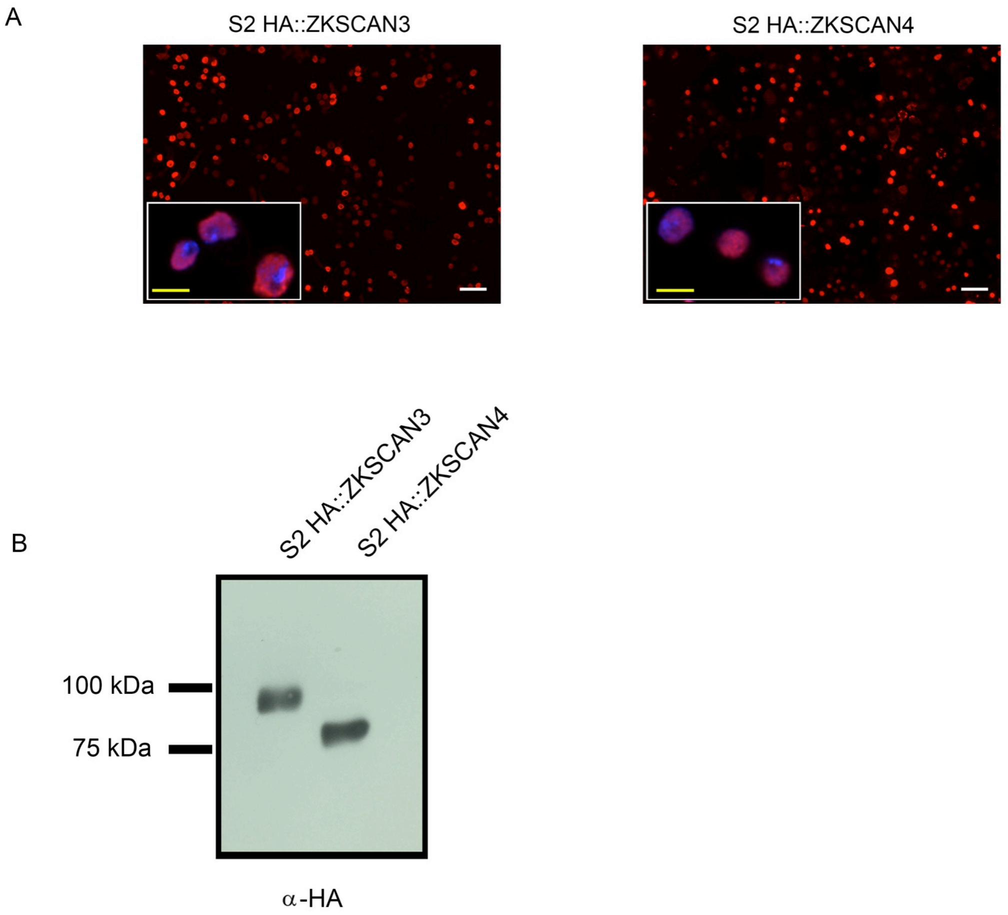


**Figure S3. Establishment of inducible Drosophila cell lines stably expressing HA-tagged vertebrate ZKSCAN3 or ZKSCAN4.**

**(A)** Anti-HA tag immunofluorescent staining (red channel) of HA::ZKSCAN3 (left) or HA::ZKSCAN4 (right) stable expressing S2 cells are shown showing the nuclear localisation of either protein (inset; nuclei counterstained with DAPI). Scale bars represent 25μm (white bars) or 5μm (yellow bars in inset). **(B)** Anti-HA immunoblot of whole cell extracts from induced HA::ZKSCAN3 or HA::ZKSCAN4 stable expressing S2 cells shows expected molecular weights for expression of full length exogenous proteins.
