## Supplementary Table 1 for "Human ZKSCAN3 and *Drosophila* M1BP are functionally homologous transcription factors in autophagy regulation"

**Table S1. Differentially expressed genes in the L3F stage with M1BP RNAi and ZKSCAN3 genotypes and L3W stages.**

L3F cgGal4>M1BP RNAi vs L3F

| Flybase ID | log2(Fold Change) | p value |
|---|---|---|
| FBgn0013278 | 6.94 | 6.28E-60 |
| FBgn0033927 | 6.69 | 9.02E-55 |
| FBgn0267798 | 6.41 | 8.08E-19 |
| FBgn0004034 | 6.06 | 1.79E-16 |
| FBgn0037807 | 6.01 | 8.36E-43 |
| FBgn0283461 | 5.82 | 1.62E-22 |
| FBgn0033926 | 5.72 | 2.84E-84 |
| FBgn0051288 | 5.51 | 1.20E-123 |
| FBgn0263321 | 5.40 | 8.89E-44 |
| FBgn0040251 | 5.39 | 2.02E-25 |
| FBgn0040736 | 5.17 | 3.36E-35 |
| FBgn0039316 | 5.09 | 8.62E-161 |
| FBgn0266873 | 5.05 | 1.77E-10 |
| FBgn0037386 | 5.00 | 1.38E-37 |
| FBgn0051104 | 4.79 | 4.02E-16 |
| FBgn0265652 | 4.75 | 1.21E-16 |
| FBgn0051464 | 4.75 | 2.12E-13 |
| FBgn0259710 | 4.74 | 1.17E-26 |
| FBgn0004635 | 4.72 | 2.04E-12 |
| FBgn0266702 | 4.70 | 5.03E-09 |
| FBgn0260795 | 4.60 | 9.29E-45 |
| FBgn0039620 | 4.57 | 8.36E-35 |
| FBgn0050025 | 4.56 | 5.70E-17 |
| FBgn0001225 | 4.55 | 1.69E-72 |
| FBgn0031888 | 4.52 | 8.55E-15 |
| FBgn0050052 | 4.40 | 1.80E-17 |
| FBgn0039319 | 4.37 | 2.13E-72 |
| FBgn0053503 | 4.27 | 1.81E-41 |
| FBgn0004865 | 4.23 | 3.81E-09 |
| FBgn0004919 | 4.23 | 2.27E-34 |
| FBgn0053468 | 4.23 | 4.55E-08 |
| FBgn0041225 | 4.23 | 2.45E-19 |
| FBgn0259708 | 4.12 | 3.14E-147 |
| FBgn0259709 | 4.12 | 3.14E-147 |
| FBgn0025583 | 4.11 | 9.04E-09 |
| FBgn0029930 | 4.04 | 2.50E-15 |
| FBgn0034005 | 4.01 | 8.81E-17 |
| FBgn0038455 | 3.99 | 7.10E-82 |
| FBgn0032810 | 3.93 | 5.55E-26 |
| FBgn0053192 | 3.92 | 4.67E-14 |
| FBgn0039915 | 3.90 | 1.11E-50 |
| FBgn0013276 | 3.88 | 6.84E-07 |
| FBgn0033875 | 3.85 | 5.26E-28 |
| FBgn0037385 | 3.85 | 2.08E-06 |
| FBgn0040837 | 3.80 | 1.09E-62 |
| FBgn0034915 | 3.79 | 1.33E-100 |
| FBgn0053126 | 3.78 | 4.86E-14 |
| FBgn0260660 | 3.77 | 3.11E-55 |
| FBgn0003886 | 3.77 | 1.96E-08 |
| FBgn0035476 | 3.76 | 2.15E-12 |
| FBgn0031220 | 3.75 | 9.22E-15 |
| FBgn0037612 | 3.74 | 3.84E-15 |
| FBgn0010278 | 3.73 | 5.28E-170 |
| FBgn0260006 | 3.69 | 1.20E-19 |
| FBgn0264542 | 3.66 | 1.30E-07 |
| FBgn0259711 | 3.64 | 1.18E-40 |
| FBgn0001226 | 3.62 | 2.67E-11 |
| FBgn0030438 | 3.61 | 1.97E-05 |
| FBgn0050026 | 3.61 | 1.57E-14 |
| FBgn0051676 | 3.60 | 2.10E-11 |
| FBgn0265651 | 3.59 | 6.78E-11 |
| FBgn0001224 | 3.55 | 2.95E-05 |
| FBgn0265927 | 3.54 | 3.00E-05 |
| FBgn0267482 | 3.50 | 1.69E-05 |
| FBgn0267153 | 3.47 | 1.34E-07 |
| FBgn0051076 | 3.46 | 1.61E-25 |
| FBgn0034162 | 3.45 | 1.10E-13 |
| FBgn0000053 | 3.44 | 1.51E-18 |
| FBgn0026616 | 3.41 | 1.87E-11 |
| FBgn0000071 | 3.39 | 1.65E-11 |
| FBgn0034605 | 3.37 | 1.34E-07 |
| FBgn0266420 | 3.36 | 5.40E-31 |
| FBgn0003162 | 3.36 | 3.31E-16 |
| FBgn0039593 | 3.33 | 5.87E-12 |
| FBgn0030241 | 3.32 | 5.23E-13 |
| FBgn0051897 | 3.29 | 1.11E-08 |
| FBgn0001217 | 3.28 | 1.06E-05 |
| FBgn0035996 | 3.28 | 1.06E-61 |
| FBgn0034670 | 3.27 | 1.53E-09 |
| FBgn0037345 | 3.26 | 1.04E-54 |
| FBgn0260238 | 3.25 | 6.84E-28 |
| FBgn0052573 | 3.25 | 6.84E-28 |
| FBgn0039788 | 3.23 | 5.17E-05 |
| FBgn0051279 | 3.23 | 9.37E-05 |

L3F cgGal4>M1BP RNAi; ZKSCAN3 vs L3F

| Flybase ID | log2(Fold Change) | p value |
|---|---|---|
| FBgn0013278 | 8.73 | 5.08E-98 |
| FBgn0040736 | 7.78 | 2.53E-104 |
| FBgn0035434 | 7.66 | 3.22E-105 |
| FBgn0283461 | 7.52 | 3.34E-41 |
| FBgn0025583 | 7.43 | 2.35E-48 |
| FBgn0034328 | 6.65 | 3.53E-36 |
| FBgn0036910 | 6.60 | 7.22E-28 |
| FBgn0265652 | 6.40 | 4.28E-63 |
| FBgn0263321 | 6.32 | 1.07E-50 |
| FBgn0040653 | 6.32 | 8.63E-25 |
| FBgn0067905 | 6.27 | 2.65E-23 |
| FBgn0053470 | 6.14 | 9.87E-75 |
| FBgn0040735 | 6.12 | 1.94E-33 |
| FBgn0038523 | 6.10 | 3.72E-34 |
| FBgn0004592 | 6.09 | 3.99E-64 |
| FBgn0283462 | 6.03 | 3.21E-54 |
| FBgn0040251 | 6.00 | 1.40E-32 |
| FBgn0003372 | 5.97 | 2.79E-32 |
| FBgn0265577 | 5.84 | 1.55E-37 |
| FBgn0033927 | 5.84 | 4.74E-45 |
| FBgn0051326 | 5.69 | 4.97E-54 |
| FBgn0037934 | 5.68 | 7.40E-24 |
| FBgn0003374 | 5.64 | 5.39E-48 |
| FBgn0003373 | 5.51 | 1.54E-49 |
| FBgn0260795 | 5.39 | 4.29E-31 |
| FBgn0003375 | 5.39 | 5.35E-27 |
| FBgn0033250 | 5.36 | 3.07E-23 |
| FBgn0262794 | 5.31 | 3.03E-30 |
| FBgn0040734 | 5.27 | 2.33E-15 |
| FBgn0265651 | 5.14 | 2.87E-45 |
| FBgn0039525 | 5.14 | 1.26E-36 |
| FBgn0003377 | 5.13 | 1.70E-41 |
| FBgn0033926 | 5.11 | 1.81E-56 |
| FBgn0003378 | 5.03 | 5.01E-46 |
| FBgn0038083 | 5.02 | 4.57E-27 |
| FBgn0051219 | 5.01 | 6.22E-14 |
| FBgn0028948 | 4.99 | 9.38E-14 |
| FBgn0013710 | 4.98 | 2.31E-38 |
| FBgn0035476 | 4.98 | 2.48E-18 |
| FBgn0039319 | 4.94 | 1.62E-31 |
| FBgn0050052 | 4.85 | 4.05E-30 |
| FBgn0050026 | 4.79 | 2.53E-27 |
| FBgn0037386 | 4.77 | 1.53E-32 |
| FBgn0031307 | 4.67 | 3.02E-23 |
| FBgn0051464 | 4.65 | 1.90E-17 |
| FBgn0266702 | 4.63 | 1.27E-11 |
| FBgn0051288 | 4.59 | 6.58E-49 |
| FBgn0040733 | 4.57 | 4.71E-11 |
| FBgn0034011 | 4.56 | 2.10E-45 |
| FBgn0036950 | 4.55 | 1.20E-13 |
| FBgn0261702 | 4.54 | 4.93E-11 |
| FBgn0013690 | 4.54 | 1.06E-28 |
| FBgn0052224 | 4.53 | 4.84E-12 |
| FBgn0037477 | 4.51 | 7.33E-11 |
| FBgn0266047 | 4.49 | 2.60E-12 |
| FBgn0032472 | 4.40 | 4.20E-16 |
| FBgn0034605 | 4.40 | 1.48E-14 |
| FBgn0030432 | 4.39 | 1.62E-15 |
| FBgn0040565 | 4.37 | 6.64E-27 |
| FBgn0051698 | 4.33 | 2.14E-29 |
| FBgn0030443 | 4.28 | 1.56E-22 |
| FBgn0034480 | 4.22 | 1.11E-14 |
| FBgn0004034 | 4.21 | 2.30E-09 |
| FBgn0040602 | 4.19 | 2.30E-10 |
| FBgn0263083 | 4.18 | 7.80E-17 |
| FBgn0003886 | 4.12 | 9.89E-14 |
| FBgn0266114 | 4.06 | 2.11E-14 |
| FBgn0034005 | 4.05 | 6.76E-21 |
| FBgn0034329 | 4.00 | 4.60E-22 |
| FBgn0051676 | 3.97 | 7.15E-16 |
| FBgn0053329 | 3.97 | 1.77E-16 |
| FBgn0039316 | 3.97 | 1.87E-21 |
| FBgn0038082 | 3.95 | 5.73E-08 |
| FBgn0010241 | 3.92 | 1.36E-14 |
| FBgn0054034 | 3.86 | 2.05E-17 |
| FBgn0034331 | 3.86 | 7.33E-20 |
| FBgn0036467 | 3.84 | 4.12E-26 |
| FBgn0053514 | 3.84 | 2.67E-33 |
| FBgn0031512 | 3.83 | 5.50E-19 |
| FBgn0033821 | 3.68 | 5.94E-24 |
| FBgn0038020 | 3.66 | 9.00E-43 |
| FBgn0266253 | 3.61 | 7.66E-08 |
| FBgn0085308 | 3.59 | 5.61E-11 |
| FBgn0036324 | 3.58 | 3.56E-09 |

L3F cgGal4> ZKSCAN3 vs L3F

| Flybase ID | log2(Fold Change) | p value |
|---|---|---|
| FBgn0001224 | 2.52 | 6.87E-13 |
| FBgn0000639 | 2.43 | 4.15E-24 |
| FBgn0000053 | 2.04 | 3.65E-10 |
| FBgn0001225 | 1.93 | 1.11E-10 |
| FBgn0036023 | 1.80 | 4.23E-07 |
| FBgn0039241 | 1.79 | 3.97E-10 |
| FBgn0051076 | 1.78 | 2.77E-08 |
| FBgn0035041 | 1.78 | 5.03E-09 |
| FBgn0000071 | 1.73 | 2.29E-07 |
| FBgn0003046 | 1.71 | 1.22E-06 |
| FBgn0063667 | 1.69 | 1.10E-06 |
| FBgn0037612 | 1.69 | 2.40E-06 |
| FBgn0032713 | 1.68 | 9.69E-09 |
| FBgn0260238 | 1.67 | 2.83E-07 |
| FBgn0052573 | 1.67 | 2.83E-07 |
| FBgn0001226 | 1.67 | 1.65E-06 |
| FBgn0039915 | 1.62 | 1.03E-06 |
| FBgn0040060 | 1.33 | 1.68E-07 |
| FBgn0022774 | 1.31 | 1.22E-11 |
| FBgn0033439 | 1.16 | 7.00E-07 |
| FBgn0035665 | 1.03 | 6.92E-07 |
| FBgn0051559 | -1.72 | 1.42E-06 |
| FBgn0039486 | -1.77 | 4.55E-08 |
| FBgn0266402 | -2.12 | 3.40E-09 |
| FBgn0045478 | -2.65 | 2.48E-14 |
| FBgn0045477 | -2.65 | 2.48E-14 |
| FBgn0052255 | -3.04 | 9.45E-18 |
| FBgn0045476 | -3.63 | 1.34E-26 |
| FBgn0035486 | -3.65 | 7.93E-27 |

| | | | | | |
|---|---|---|---|---|---|
| FBgn0053542 | 3.21 | 3.44E-05 | FBgn0053503 | 3.58 | 3.99E-26 |
| FBgn0033821 | 3.20 | 8.83E-63 | FBgn0039593 | 3.53 | 3.35E-22 |
| FBgn0003374 | 3.20 | 4.00E-05 | FBgn0036422 | 3.52 | 2.13E-57 |
| FBgn0015351 | 3.16 | 3.00E-30 | FBgn0023094 | 3.46 | 6.23E-13 |
| FBgn0036950 | 3.15 | 5.66E-05 | FBgn0037385 | 3.45 | 1.05E-06 |
| FBgn0054034 | 3.15 | 7.10E-09 | FBgn0264542 | 3.40 | 1.30E-08 |
| FBgn0034756 | 3.12 | 9.09E-180 | FBgn0267281 | 3.39 | 1.07E-06 |
| FBgn0035265 | 3.10 | 1.47E-05 | FBgn0039620 | 3.39 | 6.20E-14 |
| FBgn0039742 | 3.09 | 3.11E-08 | FBgn0028949 | 3.37 | 4.53E-07 |
| FBgn0030189 | 3.09 | 2.50E-64 | FBgn0040837 | 3.33 | 1.91E-21 |
| FBgn0263076 | 3.07 | 4.73E-05 | FBgn0051793 | 3.32 | 4.76E-19 |
| FBgn0004592 | 3.06 | 5.04E-05 | FBgn0053542 | 3.30 | 7.07E-07 |
| FBgn0037811 | 3.04 | 9.28E-05 | FBgn0041087 | 3.27 | 3.44E-08 |
| FBgn0034740 | 3.04 | 1.90E-127 | FBgn0259950 | 3.25 | 1.01E-07 |
| FBgn0032981 | 3.03 | 2.06E-07 | FBgn0040259 | 3.25 | 1.41E-11 |
| FBgn0267028 | 3.03 | 0.00011083 | FBgn0264089 | 3.23 | 5.08E-13 |
| FBgn0031476 | 3.02 | 2.25E-17 | FBgn0038828 | 3.23 | 4.95E-15 |
| FBgn0003377 | 3.02 | 4.37E-05 | FBgn0265927 | 3.19 | 1.61E-05 |
| FBgn0036101 | 3.02 | 4.92E-15 | FBgn0264748 | 3.18 | 1.56E-10 |
| FBgn0000639 | 3.01 | 1.34E-43 | FBgn0010039 | 3.18 | 4.41E-24 |
| FBgn0038470 | 3.01 | 3.30E-12 | FBgn0001225 | 3.16 | 6.75E-24 |
| FBgn0034052 | 3.00 | 8.29E-07 | FBgn0263762 | 3.13 | 2.39E-05 |
| FBgn0261555 | 3.00 | 8.69E-15 | FBgn0051528 | 3.08 | 5.14E-08 |
| FBgn0036849 | 3.00 | 9.87E-05 | FBgn0038455 | 3.06 | 1.75E-20 |
| FBgn0051469 | 2.98 | 4.81E-06 | FBgn0052351 | 3.05 | 3.06E-11 |
| FBgn0034076 | 2.97 | 2.58E-54 | FBgn0034756 | 3.04 | 4.92E-23 |
| FBgn0051793 | 2.97 | 3.54E-35 | FBgn0020508 | 3.04 | 1.87E-07 |
| FBgn0033250 | 2.96 | 4.36E-10 | FBgn0266873 | 3.03 | 4.82E-05 |
| FBgn0036186 | 2.94 | 6.28E-13 | FBgn0264745 | 3.02 | 1.15E-06 |
| FBgn0053514 | 2.94 | 7.80E-11 | FBgn0260463 | 3.01 | 1.32E-33 |
| FBgn0037613 | 2.94 | 1.06E-10 | FBgn0001228 | 3.01 | 2.04E-09 |
| FBgn0039044 | 2.92 | 4.54E-29 | FBgn0001223 | 3.01 | 2.08E-09 |
| FBgn0003378 | 2.90 | 0.00012544 | FBgn0034761 | 3.00 | 6.81E-19 |
| FBgn0030040 | 2.88 | 6.56E-22 | FBgn0053105 | 2.99 | 5.77E-33 |
| FBgn0038953 | 2.88 | 1.86E-54 | FBgn0002571 | 2.97 | 2.23E-05 |
| FBgn0029766 | 2.86 | 3.60E-66 | FBgn0038470 | 2.96 | 1.23E-10 |
| FBgn0003255 | 2.84 | 1.60E-15 | FBgn0034740 | 2.95 | 2.75E-21 |
| FBgn0264006 | 2.83 | 2.05E-05 | FBgn0033875 | 2.94 | 4.96E-06 |
| FBgn0034728 | 2.82 | 9.55E-31 | FBgn0031741 | 2.94 | 5.31E-13 |
| FBgn0262123 | 2.81 | 2.32E-15 | FBgn0267366 | 2.92 | 3.68E-08 |
| FBgn0010222 | 2.81 | 2.39E-07 | FBgn0262945 | 2.89 | 5.52E-17 |
| FBgn0010173 | 2.80 | 1.14E-63 | FBgn0030028 | 2.89 | 1.01E-06 |
| FBgn0031489 | 2.79 | 1.85E-15 | FBgn0053191 | 2.88 | 3.56E-09 |
| FBgn0052195 | 2.78 | 2.41E-24 | FBgn0040104 | 2.85 | 5.45E-08 |
| FBgn0036454 | 2.78 | 1.25E-14 | FBgn0003082 | 2.83 | 2.95E-14 |
| FBgn0038716 | 2.75 | 4.82E-49 | FBgn0262352 | 2.81 | 9.33E-07 |
| FBgn0053191 | 2.74 | 6.27E-06 | FBgn0031489 | 2.80 | 5.76E-08 |
| FBgn0002576 | 2.74 | 7.57E-06 | FBgn0034670 | 2.79 | 1.56E-09 |
| FBgn0031896 | 2.73 | 2.83E-06 | FBgn0037326 | 2.78 | 2.98E-08 |
| FBgn0264089 | 2.72 | 9.53E-14 | FBgn0032390 | 2.74 | 9.99E-10 |
| FBgn0010039 | 2.72 | 3.83E-23 | FBgn0036995 | 2.73 | 6.77E-06 |
| FBgn0033367 | 2.72 | 5.27E-10 | FBgn0031907 | 2.70 | 1.35E-05 |
| FBgn0041337 | 2.71 | 3.01E-06 | FBgn0032668 | 2.70 | 1.47E-06 |
| FBgn0061476 | 2.71 | 2.68E-05 | FBgn0050025 | 2.69 | 2.85E-05 |
| FBgn0052091 | 2.70 | 4.34E-30 | FBgn0031483 | 2.68 | 5.29E-15 |
| FBgn0283741 | 2.70 | 4.27E-07 | FBgn0000053 | 2.68 | 6.36E-07 |
| FBgn0039241 | 2.70 | 1.80E-26 | FBgn0260660 | 2.67 | 2.42E-13 |
| FBgn0033440 | 2.70 | 3.09E-07 | FBgn0260006 | 2.66 | 4.54E-11 |
| FBgn0039328 | 2.69 | 6.62E-23 | FBgn0031039 | 2.65 | 5.06E-13 |
| FBgn0003888 | 2.69 | 2.79E-14 | FBgn0031038 | 2.65 | 5.06E-13 |
| FBgn0267792 | 2.68 | 1.46E-05 | FBgn0039759 | 2.65 | 2.93E-05 |
| FBgn0030864 | 2.67 | 3.00E-10 | FBgn0044812 | 2.64 | 2.36E-11 |
| FBgn0034011 | 2.67 | 1.90E-18 | FBgn0031263 | 2.64 | 5.23E-10 |
| FBgn0031643 | 2.67 | 2.54E-59 | FBgn0041225 | 2.63 | 2.05E-07 |
| FBgn0033439 | 2.66 | 2.43E-14 | FBgn0260933 | 2.63 | 5.16E-10 |
| FBgn0030756 | 2.66 | 1.21E-05 | FBgn0264834 | 2.62 | 9.54E-13 |
| FBgn0030097 | 2.65 | 9.05E-07 | FBgn0264740 | 2.61 | 3.79E-05 |
| FBgn0020270 | 2.65 | 3.21E-15 | FBgn0264741 | 2.61 | 3.79E-05 |
| FBgn0038353 | 2.63 | 4.92E-05 | FBgn0283741 | 2.60 | 1.76E-15 |
| FBgn0283437 | 2.63 | 8.26E-11 | FBgn0010278 | 2.59 | 2.55E-25 |
| FBgn0058006 | 2.63 | 3.80E-07 | FBgn0053126 | 2.57 | 2.67E-07 |
| FBgn0262146 | 2.61 | 0.00010001 | FBgn0032810 | 2.55 | 2.78E-11 |
| FBgn0032390 | 2.61 | 3.47E-07 | FBgn0037531 | 2.55 | 8.35E-10 |
| FBgn0034914 | 2.61 | 4.23E-06 | FBgn0030040 | 2.54 | 4.55E-14 |
| FBgn0037531 | 2.60 | 2.42E-10 | FBgn0031649 | 2.54 | 1.38E-05 |
| FBgn0030837 | 2.60 | 1.02E-07 | FBgn0031432 | 2.51 | 6.81E-19 |
| FBgn0024321 | 2.59 | 2.73E-11 | FBgn0037126 | 2.49 | 6.69E-10 |
| FBgn0031760 | 2.59 | 1.48E-14 | FBgn0031068 | 2.48 | 4.90E-08 |
| FBgn0051633 | 2.58 | 3.18E-32 | FBgn0053192 | 2.48 | 7.18E-08 |
| FBgn0013765 | 2.57 | 7.77E-14 | FBgn0013765 | 2.48 | 2.46E-13 |
| FBgn0039099 | 2.57 | 4.53E-08 | FBgn0061476 | 2.48 | 1.01E-05 |
| FBgn0029831 | 2.55 | 2.42E-18 | FBgn0033775 | 2.46 | 5.84E-11 |
| FBgn0038893 | 2.52 | 2.57E-22 | FBgn0000490 | 2.46 | 2.10E-07 |
| FBgn0003082 | 2.52 | 2.71E-09 | FBgn0034275 | 2.46 | 3.56E-19 |
| FBgn0063667 | 2.50 | 9.01E-19 | FBgn0026751 | 2.45 | 1.62E-09 |
| FBgn0003046 | 2.50 | 7.55E-05 | FBgn0053462 | 2.45 | 6.86E-09 |
| FBgn0051121 | 2.49 | 1.62E-07 | FBgn0010786 | 2.45 | 9.99E-09 |
| FBgn0031515 | 2.48 | 1.45E-25 | FBgn0034709 | 2.45 | 3.29E-11 |
| FBgn0027932 | 2.48 | 8.79E-15 | FBgn0063494 | 2.40 | 7.88E-11 |
| FBgn0034480 | 2.48 | 2.03E-05 | FBgn0032422 | 2.38 | 9.52E-07 |

| | | | | | |
|---|---|---|---|---|---|
| FBgn0028939 | 2.44 | 2.41E-11 | FBgn0031760 | 2.38 | 1.06E-10 |
| FBgn0028411 | 2.44 | 1.56E-21 | FBgn0036764 | 2.36 | 2.32E-08 |
| FBgn0264834 | 2.43 | 1.91E-06 | FBgn0030241 | 2.36 | 1.51E-07 |
| FBgn0260474 | 2.43 | 8.85E-10 | FBgn0261363 | 2.36 | 1.36E-10 |
| FBgn0040364 | 2.43 | 2.75E-09 | FBgn0034052 | 2.34 | 2.15E-05 |
| FBgn0051363 | 2.43 | 5.98E-10 | FBgn0266420 | 2.34 | 1.86E-10 |
| FBgn0050015 | 2.42 | 6.56E-14 | FBgn0035623 | 2.33 | 2.72E-14 |
| FBgn0039003 | 2.42 | 5.24E-31 | FBgn0260955 | 2.32 | 1.89E-10 |
| FBgn0036187 | 2.41 | 5.50E-26 | FBgn0037612 | 2.31 | 6.19E-05 |
| FBgn0004698 | 2.40 | 1.98E-35 | FBgn0259710 | 2.29 | 1.60E-05 |
| FBgn0026751 | 2.39 | 6.46E-15 | FBgn0262902 | 2.29 | 3.76E-09 |
| FBgn0020445 | 2.39 | 6.20E-07 | FBgn0023023 | 2.27 | 8.64E-10 |
| FBgn0040060 | 2.39 | 8.06E-12 | FBgn0013709 | 2.26 | 2.24E-07 |
| FBgn0011589 | 2.36 | 8.99E-05 | FBgn0262123 | 2.26 | 3.80E-10 |
| FBgn0020304 | 2.36 | 6.64E-09 | FBgn0041183 | 2.24 | 1.76E-13 |
| FBgn0262036 | 2.35 | 1.15E-27 | FBgn0033304 | 2.23 | 2.83E-07 |
| FBgn0046876 | 2.35 | 2.52E-05 | FBgn0011592 | 2.22 | 1.84E-10 |
| FBgn0013972 | 2.34 | 5.75E-16 | FBgn0034728 | 2.22 | 4.22E-12 |
| FBgn0030836 | 2.34 | 5.19E-09 | FBgn0028738 | 2.22 | 4.51E-08 |
| FBgn0038082 | 2.33 | 2.08E-11 | FBgn0051436 | 2.21 | 2.52E-12 |
| FBgn0033649 | 2.33 | 1.06E-05 | FBgn0033372 | 2.21 | 2.07E-06 |
| FBgn0032713 | 2.33 | 1.60E-07 | FBgn0046776 | 2.20 | 1.09E-07 |
| FBgn0261530 | 2.32 | 1.42E-28 | FBgn0034915 | 2.20 | 8.16E-16 |
| FBgn0029507 | 2.32 | 2.06E-63 | FBgn0037672 | 2.19 | 1.49E-06 |
| FBgn0035434 | 2.31 | 9.87E-07 | FBgn0027544 | 2.18 | 3.57E-05 |
| FBgn0051344 | 2.31 | 1.50E-11 | FBgn0261704 | 2.18 | 3.57E-05 |
| FBgn0026084 | 2.30 | 1.61E-12 | FBgn0003888 | 2.18 | 3.19E-10 |
| FBgn0051694 | 2.30 | 9.53E-30 | FBgn0034253 | 2.17 | 1.04E-11 |
| FBgn0264869 | 2.30 | 1.23E-06 | FBgn0028514 | 2.17 | 1.05E-13 |
| FBgn0267727 | 2.30 | 6.44E-05 | FBgn0034512 | 2.16 | 8.53E-09 |
| FBgn0003475 | 2.29 | 3.60E-17 | FBgn0038716 | 2.16 | 4.28E-06 |
| FBgn0053158 | 2.29 | 8.45E-45 | FBgn0015801 | 2.15 | 2.21E-09 |
| FBgn0250869 | 2.27 | 3.43E-12 | FBgn0266005 | 2.14 | 1.18E-09 |
| FBgn0261244 | 2.27 | 8.70E-10 | FBgn0020270 | 2.12 | 6.82E-10 |
| FBgn0267795 | 2.26 | 4.30E-19 | FBgn0038302 | 2.12 | 4.16E-14 |
| FBgn0030777 | 2.25 | 5.28E-06 | FBgn0042106 | 2.12 | 2.62E-05 |
| FBgn0014342 | 2.24 | 6.49E-05 | FBgn0038107 | 2.11 | 9.55E-07 |
| FBgn0032549 | 2.24 | 1.16E-07 | FBgn0265083 | 2.11 | 9.55E-07 |
| FBgn0032253 | 2.24 | 3.68E-05 | FBgn0260985 | 2.10 | 8.51E-09 |
| FBgn0033304 | 2.23 | 7.91E-13 | FBgn0033205 | 2.08 | 1.23E-09 |
| FBgn0034709 | 2.23 | 1.33E-21 | FBgn0263600 | 2.08 | 1.35E-07 |
| FBgn0267736 | 2.22 | 3.50E-05 | FBgn0032638 | 2.06 | 8.29E-14 |
| FBgn0038302 | 2.22 | 3.86E-26 | FBgn0029507 | 2.05 | 4.97E-19 |
| FBgn0030673 | 2.21 | 5.86E-06 | FBgn0015351 | 2.05 | 6.09E-08 |
| FBgn0020388 | 2.21 | 5.21E-33 | FBgn0250869 | 2.05 | 1.48E-05 |
| FBgn0002525 | 2.18 | 6.20E-21 | FBgn0038893 | 2.04 | 5.35E-10 |
| FBgn0019952 | 2.17 | 7.98E-17 | FBgn0010042 | 2.04 | 5.87E-06 |
| FBgn0259173 | 2.17 | 8.20E-12 | FBgn0015714 | 2.02 | 4.30E-05 |
| FBgn0004858 | 2.17 | 6.20E-07 | FBgn0041710 | 2.02 | 8.86E-12 |
| FBgn0051789 | 2.15 | 1.30E-09 | FBgn0011769 | 2.02 | 2.36E-08 |
| FBgn0039529 | 2.13 | 5.12E-05 | FBgn0035996 | 2.01 | 2.12E-13 |
| FBgn0260463 | 2.12 | 2.95E-16 | FBgn0033978 | 2.01 | 3.33E-05 |
| FBgn0262902 | 2.11 | 4.54E-06 | FBgn0030628 | 2.01 | 5.23E-07 |
| FBgn0032934 | 2.10 | 2.79E-08 | FBgn0085452 | 2.01 | 4.64E-14 |
| FBgn0036144 | 2.10 | 6.58E-09 | FBgn0036403 | 2.00 | 4.13E-05 |
| FBgn0266084 | 2.10 | 1.27E-19 | FBgn0038953 | 1.99 | 2.02E-11 |
| FBgn0053105 | 2.10 | 1.05E-15 | FBgn0261004 | 1.99 | 3.61E-08 |
| FBgn0262524 | 2.09 | 1.35E-23 | FBgn0001145 | 1.98 | 9.98E-11 |
| FBgn0045761 | 2.08 | 9.67E-36 | FBgn0034362 | 1.98 | 5.27E-06 |
| FBgn0004512 | 2.08 | 2.20E-06 | FBgn0044872 | 1.97 | 7.51E-06 |
| FBgn0035665 | 2.07 | 1.34E-10 | FBgn0033153 | 1.96 | 4.61E-06 |
| FBgn0028978 | 2.07 | 4.08E-52 | FBgn0034259 | 1.95 | 3.14E-06 |
| FBgn0036806 | 2.06 | 5.47E-41 | FBgn0265053 | 1.94 | 2.45E-11 |
| FBgn0035624 | 2.05 | 1.83E-13 | FBgn0023407 | 1.94 | 3.67E-08 |
| FBgn0085249 | 2.04 | 9.49E-06 | FBgn0030189 | 1.93 | 2.83E-13 |
| FBgn0038647 | 2.04 | 1.59E-11 | FBgn0004698 | 1.93 | 2.17E-11 |
| FBgn0003117 | 2.04 | 2.71E-07 | FBgn0028396 | 1.90 | 8.84E-09 |
| FBgn0011638 | 2.04 | 9.71E-31 | FBgn0034931 | 1.90 | 7.49E-08 |
| FBgn0033945 | 2.04 | 1.12E-18 | FBgn0041630 | 1.90 | 1.81E-11 |
| FBgn0266421 | 2.03 | 3.65E-20 | FBgn0039044 | 1.89 | 2.94E-11 |
| FBgn0039488 | 2.02 | 1.02E-16 | FBgn0029664 | 1.89 | 2.49E-06 |
| FBgn0063494 | 2.02 | 4.79E-17 | FBgn0052512 | 1.89 | 1.41E-10 |
| FBgn0036487 | 2.02 | 4.83E-34 | FBgn0052843 | 1.88 | 4.09E-07 |
| FBgn0001229 | 2.02 | 6.08E-05 | FBgn0031643 | 1.88 | 2.78E-11 |
| FBgn0030855 | 2.01 | 2.72E-16 | FBgn0026702 | 1.86 | 1.80E-05 |
| FBgn0031654 | 2.00 | 2.68E-06 | FBgn0026576 | 1.86 | 1.27E-16 |
| FBgn0039006 | 2.00 | 6.52E-20 | FBgn0046878 | 1.86 | 3.77E-05 |
| FBgn0010358 | 1.99 | 2.52E-06 | FBgn0051344 | 1.86 | 1.23E-05 |
| FBgn0036952 | 1.99 | 6.46E-07 | FBgn0266570 | 1.86 | 1.75E-07 |
| FBgn0020647 | 1.99 | 3.12E-08 | FBgn0024947 | 1.86 | 4.17E-05 |
| FBgn0034221 | 1.98 | 4.11E-05 | FBgn0011638 | 1.81 | 2.08E-08 |
| FBgn0030183 | 1.98 | 2.26E-06 | FBgn0030052 | 1.81 | 8.01E-07 |
| FBgn0023129 | 1.98 | 4.60E-16 | FBgn0033438 | 1.80 | 3.71E-07 |
| FBgn0260985 | 1.97 | 6.01E-10 | FBgn0010173 | 1.80 | 2.20E-10 |
| FBgn0029993 | 1.97 | 2.34E-05 | FBgn0036702 | 1.79 | 1.24E-06 |
| FBgn0086370 | 1.97 | 6.64E-13 | FBgn0027657 | 1.78 | 9.57E-06 |
| FBgn0034741 | 1.96 | 6.37E-28 | FBgn0040299 | 1.78 | 9.29E-07 |
| FBgn0033623 | 1.96 | 0.00011537 | FBgn0003062 | 1.77 | 2.13E-06 |
| FBgn0263600 | 1.96 | 6.03E-11 | FBgn0015037 | 1.77 | 3.52E-07 |
| FBgn0011774 | 1.96 | 6.33E-11 | FBgn0039528 | 1.75 | 2.41E-09 |

| | | | | | |
|---|---|---|---|---|---|
| FBgn0028430 | 1.95 | 2.57E-19 | FBgn0052756 | 1.75 | 2.85E-07 |
| FBgn0023407 | 1.94 | 5.72E-07 | FBgn0015393 | 1.74 | 5.59E-06 |
| FBgn0033153 | 1.94 | 3.82E-14 | FBgn0003278 | 1.73 | 4.74E-05 |
| FBgn0052756 | 1.93 | 1.21E-07 | FBgn0038072 | 1.73 | 1.56E-06 |
| FBgn0044823 | 1.93 | 1.77E-11 | FBgn0263241 | 1.72 | 6.93E-06 |
| FBgn0243514 | 1.92 | 3.87E-08 | FBgn0030263 | 1.70 | 5.63E-08 |
| FBgn0028738 | 1.92 | 9.81E-05 | FBgn0036173 | 1.70 | 7.68E-07 |
| FBgn0028494 | 1.91 | 6.23E-11 | FBgn0052091 | 1.70 | 1.17E-06 |
| FBgn0011592 | 1.91 | 5.31E-10 | FBgn0262473 | 1.70 | 1.68E-10 |
| FBgn0050089 | 1.91 | 6.46E-06 | FBgn0261530 | 1.69 | 7.70E-08 |
| FBgn0035889 | 1.91 | 4.77E-28 | FBgn0011774 | 1.69 | 8.46E-06 |
| FBgn0053057 | 1.91 | 4.77E-28 | FBgn0262524 | 1.69 | 4.91E-07 |
| FBgn0020906 | 1.91 | 1.42E-12 | FBgn0013688 | 1.68 | 1.79E-10 |
| FBgn0266005 | 1.89 | 4.50E-06 | FBgn0038306 | 1.68 | 1.09E-05 |
| FBgn0262737 | 1.88 | 3.52E-05 | FBgn0031114 | 1.65 | 1.30E-05 |
| FBgn0001230 | 1.88 | 5.79E-08 | FBgn0261555 | 1.65 | 1.46E-05 |
| FBgn0027657 | 1.88 | 3.05E-20 | FBgn0028411 | 1.64 | 3.43E-07 |
| FBgn0041183 | 1.88 | 1.49E-11 | FBgn0040294 | 1.64 | 6.99E-05 |
| FBgn0035983 | 1.86 | 1.40E-08 | FBgn0010222 | 1.62 | 1.70E-05 |
| FBgn0033205 | 1.86 | 6.28E-40 | FBgn0010803 | 1.61 | 3.40E-05 |
| FBgn0032026 | 1.85 | 2.59E-31 | FBgn0266084 | 1.61 | 1.77E-06 |
| FBgn0040299 | 1.85 | 1.20E-06 | FBgn0037338 | 1.61 | 2.91E-07 |
| FBgn0263235 | 1.85 | 1.36E-05 | FBgn0027499 | 1.60 | 7.26E-11 |
| FBgn0037719 | 1.85 | 1.73E-24 | FBgn0030000 | 1.59 | 2.00E-05 |
| FBgn0033775 | 1.84 | 2.13E-06 | FBgn0052207 | 1.59 | 2.02E-05 |
| FBgn0032587 | 1.82 | 7.01E-08 | FBgn0267795 | 1.58 | 6.41E-06 |
| FBgn0052475 | 1.82 | 1.18E-11 | FBgn0262560 | 1.57 | 1.64E-05 |
| FBgn0030766 | 1.82 | 4.52E-07 | FBgn0051710 | 1.57 | 1.16E-05 |
| FBgn0020762 | 1.82 | 1.09E-05 | FBgn0029672 | 1.56 | 2.03E-05 |
| FBgn0284231 | 1.81 | 1.16E-09 | FBgn0045761 | 1.56 | 6.42E-09 |
| FBgn0051431 | 1.81 | 9.00E-10 | FBgn0051549 | 1.56 | 2.68E-08 |
| FBgn0035623 | 1.81 | 6.12E-08 | FBgn0266421 | 1.55 | 7.97E-07 |
| FBgn0024913 | 1.81 | 7.51E-06 | FBgn0030855 | 1.55 | 2.03E-06 |
| FBgn0002887 | 1.80 | 1.30E-10 | FBgn0266268 | 1.55 | 1.40E-05 |
| FBgn0250815 | 1.77 | 2.20E-11 | FBgn0039328 | 1.55 | 2.50E-05 |
| FBgn0032935 | 1.77 | 4.07E-15 | FBgn0037801 | 1.55 | 1.09E-05 |
| FBgn0041710 | 1.77 | 1.54E-15 | FBgn0051777 | 1.54 | 2.58E-07 |
| FBgn0032262 | 1.77 | 9.12E-08 | FBgn0041094 | 1.54 | 3.85E-05 |
| FBgn0000246 | 1.76 | 1.08E-05 | FBgn0266581 | 1.53 | 3.54E-06 |
| FBgn0001319 | 1.75 | 3.92E-05 | FBgn0034793 | 1.53 | 4.36E-05 |
| FBgn0026319 | 1.75 | 0.00013295 | FBgn0029866 | 1.53 | 3.17E-11 |
| FBgn0038832 | 1.75 | 5.25E-06 | FBgn0029825 | 1.52 | 7.52E-07 |
| FBgn0030876 | 1.72 | 6.70E-11 | FBgn0003996 | 1.52 | 5.45E-05 |
| FBgn0026737 | 1.72 | 3.58E-12 | FBgn0051324 | 1.52 | 9.01E-07 |
| FBgn0039644 | 1.72 | 8.21E-29 | FBgn0045980 | 1.51 | 5.18E-05 |
| FBgn0003328 | 1.71 | 9.75E-13 | FBgn0033741 | 1.51 | 3.91E-05 |
| FBgn0028988 | 1.70 | 8.26E-08 | FBgn0026737 | 1.50 | 1.89E-07 |
| FBgn0039149 | 1.70 | 4.87E-10 | FBgn0037810 | 1.50 | 1.13E-06 |
| FBgn0039207 | 1.68 | 6.08E-11 | FBgn0032032 | 1.49 | 1.59E-05 |
| FBgn0264713 | 1.68 | 2.85E-10 | FBgn0034527 | 1.48 | 1.08E-06 |
| FBgn0039125 | 1.67 | 1.26E-12 | FBgn0029932 | 1.47 | 6.07E-08 |
| FBgn0028426 | 1.67 | 2.43E-14 | FBgn0029133 | 1.47 | 2.43E-05 |
| FBgn0034908 | 1.66 | 2.24E-12 | FBgn0050022 | 1.45 | 6.09E-05 |
| FBgn0037555 | 1.66 | 7.17E-27 | FBgn0260755 | 1.42 | 7.13E-06 |
| FBgn0260817 | 1.66 | 3.88E-10 | FBgn0263706 | 1.42 | 2.21E-06 |
| FBgn0004913 | 1.66 | 4.68E-12 | FBgn0033809 | 1.41 | 5.83E-09 |
| FBgn0052549 | 1.65 | 3.66E-06 | FBgn0050091 | 1.41 | 3.62E-09 |
| FBgn0032140 | 1.65 | 5.53E-06 | FBgn0037345 | 1.41 | 1.31E-06 |
| FBgn0039528 | 1.65 | 6.80E-08 | FBgn0029999 | 1.40 | 1.58E-05 |
| FBgn0051323 | 1.64 | 2.12E-08 | FBgn0028494 | 1.40 | 9.72E-06 |
| FBgn0005672 | 1.64 | 4.91E-06 | FBgn0003464 | 1.39 | 1.16E-05 |
| FBgn0031939 | 1.64 | 3.32E-05 | FBgn0029915 | 1.39 | 6.17E-05 |
| FBgn0046706 | 1.64 | 1.45E-08 | FBgn0260639 | 1.39 | 1.34E-05 |
| FBgn0260755 | 1.63 | 1.00E-23 | FBgn0020388 | 1.37 | 8.87E-06 |
| FBgn0003447 | 1.62 | 3.18E-05 | FBgn0039003 | 1.37 | 4.16E-07 |
| FBgn0037810 | 1.62 | 1.36E-05 | FBgn0002887 | 1.36 | 2.78E-05 |
| FBgn0014380 | 1.61 | 2.26E-08 | FBgn0085407 | 1.36 | 2.34E-05 |
| FBgn0039642 | 1.61 | 9.51E-10 | FBgn0020389 | 1.35 | 2.07E-07 |
| FBgn0003041 | 1.60 | 1.50E-05 | FBgn0002525 | 1.35 | 7.71E-07 |
| FBgn0037338 | 1.59 | 9.37E-06 | FBgn0013269 | 1.35 | 1.32E-05 |
| FBgn0053995 | 1.59 | 1.29E-37 | FBgn0003475 | 1.35 | 8.79E-07 |
| FBgn0264742 | 1.59 | 1.49E-37 | FBgn0052549 | 1.34 | 3.74E-05 |
| FBgn0264743 | 1.59 | 1.49E-37 | FBgn0036815 | 1.33 | 6.90E-05 |
| FBgn0259212 | 1.58 | 6.88E-11 | FBgn0039644 | 1.32 | 1.11E-07 |
| FBgn0051324 | 1.58 | 1.53E-08 | FBgn0046706 | 1.32 | 2.21E-05 |
| FBgn0041160 | 1.57 | 2.07E-05 | FBgn0284231 | 1.32 | 3.20E-05 |
| FBgn0040850 | 1.57 | 4.17E-06 | FBgn0036290 | 1.31 | 1.59E-06 |
| FBgn0029710 | 1.57 | 1.04E-07 | FBgn0266700 | 1.31 | 2.12E-05 |
| FBgn0028381 | 1.57 | 6.25E-12 | FBgn0038842 | 1.30 | 1.72E-05 |
| FBgn0028514 | 1.56 | 1.74E-17 | FBgn0039854 | 1.29 | 3.17E-05 |
| FBgn0029825 | 1.56 | 6.17E-07 | FBgn0259708 | 1.28 | 3.73E-06 |
| FBgn0266277 | 1.56 | 2.76E-10 | FBgn0259709 | 1.28 | 3.73E-06 |
| FBgn0038072 | 1.55 | 7.00E-05 | FBgn0039125 | 1.28 | 3.75E-06 |
| FBgn0259168 | 1.55 | 3.51E-07 | FBgn0034399 | 1.28 | 4.05E-07 |
| FBgn0031673 | 1.55 | 1.25E-08 | FBgn0039908 | 1.28 | 5.34E-06 |
| FBgn0028533 | 1.55 | 3.79E-05 | FBgn0264460 | 1.26 | 2.27E-05 |
| FBgn0030334 | 1.54 | 1.55E-12 | FBgn0051633 | 1.25 | 3.51E-06 |
| FBgn0035461 | 1.53 | 9.96E-06 | FBgn0260817 | 1.24 | 2.89E-05 |
| FBgn0036663 | 1.52 | 1.92E-13 | FBgn0034858 | 1.24 | 7.65E-06 |
| FBgn0085407 | 1.52 | 3.48E-06 | FBgn0036487 | 1.22 | 1.66E-05 |

| | | | | | |
|---|---|---|---|---|---|
| FBgn0015801 | 1.51 | 6.61E-08 | FBgn0010246 | 1.17 | 4.04E-06 |
| FBgn0262617 | 1.51 | 1.73E-10 | FBgn0030749 | 1.16 | 7.97E-06 |
| FBgn0030114 | 1.50 | 2.02E-07 | FBgn0032669 | 1.16 | 1.72E-06 |
| FBgn0034399 | 1.50 | 2.92E-16 | FBgn0031689 | 1.15 | 2.36E-05 |
| FBgn0041094 | 1.50 | 5.80E-32 | FBgn0032322 | 1.14 | 4.35E-07 |
| FBgn0000640 | 1.50 | 4.82E-18 | FBgn0261574 | 1.09 | 3.42E-05 |
| FBgn0014033 | 1.49 | 5.99E-06 | FBgn0265048 | 1.06 | 1.55E-05 |
| FBgn0033820 | 1.49 | 2.39E-08 | FBgn0030334 | 1.05 | 5.53E-05 |
| FBgn0263241 | 1.49 | 6.72E-08 | FBgn0034741 | 1.04 | 3.31E-05 |
| FBgn0037391 | 1.49 | 8.68E-33 | FBgn0032884 | 1.02 | 3.35E-05 |
| FBgn0037975 | 1.48 | 8.35E-06 | FBgn0263996 | 1.02 | 1.23E-05 |
| FBgn0039886 | 1.48 | 3.36E-05 | FBgn0032901 | 0.95 | 5.05E-05 |
| FBgn0041184 | 1.48 | 5.47E-05 | FBgn0030245 | -0.92 | 5.90E-05 |
| FBgn0036030 | 1.48 | 2.12E-13 | FBgn0250785 | -0.95 | 3.33E-05 |
| FBgn0037315 | 1.47 | 7.72E-09 | FBgn0267505 | -0.95 | 3.73E-05 |
| FBgn0037936 | 1.47 | 9.52E-25 | FBgn0085802 | -0.97 | 2.45E-05 |
| FBgn0025549 | 1.46 | 1.29E-07 | FBgn0027596 | -0.99 | 3.09E-05 |
| FBgn0038083 | 1.45 | 7.34E-08 | FBgn0027579 | -1.02 | 3.44E-06 |
| FBgn0051436 | 1.45 | 0.000123 | FBgn0039016 | -1.02 | 1.54E-05 |
| FBgn0030878 | 1.45 | 4.73E-15 | FBgn0010926 | -1.04 | 9.19E-06 |
| FBgn0267808 | 1.44 | 2.89E-16 | FBgn0035252 | -1.04 | 3.96E-06 |
| FBgn0033644 | 1.44 | 6.24E-06 | FBgn0001128 | -1.06 | 4.62E-05 |
| FBgn0267454 | 1.43 | 4.77E-10 | FBgn0039674 | -1.06 | 7.70E-06 |
| FBgn0015393 | 1.43 | 4.25E-06 | FBgn0034585 | -1.06 | 1.60E-05 |
| FBgn0051100 | 1.42 | 3.60E-09 | FBgn0264389 | -1.06 | 1.37E-05 |
| FBgn0266570 | 1.42 | 5.93E-08 | FBgn0024957 | -1.07 | 1.25E-05 |
| FBgn0035710 | 1.41 | 6.08E-11 | FBgn0085446 | -1.07 | 3.52E-06 |
| FBgn0052115 | 1.41 | 1.08E-08 | FBgn0039151 | -1.08 | 8.22E-06 |
| FBgn0035714 | 1.41 | 1.74E-12 | FBgn0033883 | -1.08 | 2.69E-06 |
| FBgn0010246 | 1.41 | 1.42E-15 | FBgn0000317 | -1.10 | 2.77E-05 |
| FBgn0036291 | 1.40 | 3.79E-05 | FBgn0065097 | -1.11 | 3.59E-06 |
| FBgn0032793 | 1.40 | 1.60E-09 | FBgn0261015 | -1.11 | 6.01E-06 |
| FBgn0052243 | 1.40 | 1.45E-08 | FBgn0038865 | -1.11 | 3.55E-05 |
| FBgn0041629 | 1.40 | 6.28E-19 | FBgn0035233 | -1.11 | 7.78E-06 |
| FBgn0052177 | 1.39 | 5.20E-14 | FBgn0039543 | -1.11 | 5.10E-05 |
| FBgn0032781 | 1.38 | 7.02E-19 | FBgn0034885 | -1.11 | 5.74E-06 |
| FBgn0039908 | 1.38 | 1.19E-07 | FBgn0010548 | -1.11 | 1.76E-06 |
| FBgn0010241 | 1.37 | 1.74E-06 | FBgn0032881 | -1.12 | 1.26E-06 |
| FBgn0263617 | 1.37 | 1.62E-05 | FBgn0033453 | -1.12 | 3.16E-05 |
| FBgn0027598 | 1.37 | 8.44E-09 | FBgn0046874 | -1.12 | 4.68E-06 |
| FBgn0003204 | 1.37 | 3.94E-05 | FBgn0039114 | -1.13 | 4.87E-05 |
| FBgn0033136 | 1.37 | 5.80E-07 | FBgn0023516 | -1.13 | 5.72E-06 |
| FBgn0052194 | 1.35 | 0.00010172 | FBgn0034432 | -1.13 | 4.76E-07 |
| FBgn0010300 | 1.35 | 1.31E-09 | FBgn0010516 | -1.14 | 1.82E-07 |
| FBgn0039452 | 1.35 | 6.70E-05 | FBgn0030051 | -1.15 | 4.35E-06 |
| FBgn0033137 | 1.34 | 2.50E-12 | FBgn0033458 | -1.15 | 7.16E-06 |
| FBgn0036126 | 1.34 | 2.68E-11 | FBgn0051144 | -1.16 | 2.40E-07 |
| FBgn0031384 | 1.34 | 4.11E-05 | FBgn0266758 | -1.16 | 5.17E-05 |
| FBgn0037443 | 1.33 | 2.74E-13 | FBgn0033783 | -1.16 | 2.82E-07 |
| FBgn0029002 | 1.32 | 7.32E-05 | FBgn0002719 | -1.16 | 3.48E-07 |
| FBgn0260743 | 1.32 | 0.00011297 | FBgn0031836 | -1.16 | 7.36E-06 |
| FBgn0025456 | 1.32 | 3.63E-05 | FBgn0086371 | -1.17 | 7.63E-07 |
| FBgn0030052 | 1.31 | 3.77E-05 | FBgn0038830 | -1.17 | 2.02E-06 |
| FBgn0034312 | 1.31 | 4.98E-21 | FBgn0004654 | -1.17 | 9.85E-06 |
| FBgn0037350 | 1.31 | 5.79E-05 | FBgn0029648 | -1.17 | 4.19E-06 |
| FBgn0040493 | 1.30 | 1.45E-18 | FBgn0033047 | -1.17 | 2.12E-05 |
| FBgn0052218 | 1.29 | 6.20E-05 | FBgn0030528 | -1.17 | 6.97E-05 |
| FBgn0040294 | 1.29 | 8.61E-09 | FBgn0013733 | -1.18 | 5.13E-07 |
| FBgn0002183 | 1.28 | 3.30E-15 | FBgn0262870 | -1.18 | 1.28E-05 |
| FBgn0033427 | 1.28 | 1.05E-05 | FBgn0278599 | -1.18 | 3.05E-05 |
| FBgn0000043 | 1.28 | 4.26E-20 | FBgn0037440 | -1.18 | 4.73E-06 |
| FBgn0027836 | 1.27 | 1.29E-15 | FBgn0015766 | -1.18 | 7.91E-07 |
| FBgn0034512 | 1.27 | 5.74E-12 | FBgn0036623 | -1.19 | 7.62E-07 |
| FBgn0037801 | 1.27 | 3.80E-09 | FBgn0025373 | -1.19 | 1.51E-07 |
| FBgn0085282 | 1.27 | 8.26E-05 | FBgn0037070 | -1.20 | 5.13E-08 |
| FBgn0032665 | 1.27 | 7.42E-05 | FBgn0267614 | -1.21 | 4.20E-08 |
| FBgn0039492 | 1.27 | 2.93E-07 | FBgn0035356 | -1.21 | 9.33E-06 |
| FBgn0045852 | 1.26 | 7.33E-05 | FBgn0005278 | -1.21 | 2.04E-08 |
| FBgn0026404 | 1.26 | 1.33E-05 | FBgn0040529 | -1.22 | 3.73E-06 |
| FBgn0030662 | 1.25 | 7.05E-06 | FBgn0031282 | -1.23 | 3.28E-05 |
| FBgn0015522 | 1.24 | 2.21E-07 | FBgn0030968 | -1.23 | 2.72E-07 |
| FBgn0044047 | 1.24 | 1.30E-09 | FBgn0030261 | -1.23 | 4.48E-06 |
| FBgn0003062 | 1.23 | 4.54E-06 | FBgn0053225 | -1.23 | 3.33E-07 |
| FBgn0034940 | 1.22 | 3.70E-05 | FBgn0035632 | -1.24 | 3.04E-07 |
| FBgn0034443 | 1.22 | 1.38E-07 | FBgn0034733 | -1.24 | 5.48E-08 |
| FBgn0031571 | 1.22 | 1.56E-05 | FBgn0265991 | -1.24 | 1.11E-06 |
| FBgn0027499 | 1.22 | 5.74E-10 | FBgn0064115 | -1.24 | 3.08E-05 |
| FBgn0259481 | 1.22 | 9.16E-11 | FBgn0004407 | -1.24 | 3.08E-05 |
| FBgn0038053 | 1.21 | 3.03E-06 | FBgn0051915 | -1.24 | 3.54E-08 |
| FBgn0039714 | 1.21 | 3.45E-07 | FBgn0026718 | -1.24 | 5.29E-07 |
| FBgn0267587 | 1.21 | 9.27E-07 | FBgn0052699 | -1.25 | 6.48E-05 |
| FBgn0042175 | 1.21 | 2.09E-10 | FBgn0266417 | -1.25 | 2.09E-05 |
| FBgn0039733 | 1.20 | 3.99E-06 | FBgn0010053 | -1.25 | 2.62E-08 |
| FBgn0034527 | 1.20 | 4.10E-07 | FBgn0034902 | -1.26 | 7.46E-07 |
| FBgn0004606 | 1.20 | 9.30E-07 | FBgn0042138 | -1.27 | 2.42E-05 |
| FBgn0036846 | 1.20 | 2.42E-06 | FBgn0035147 | -1.27 | 5.14E-07 |
| FBgn0037974 | 1.20 | 7.01E-06 | FBgn0032160 | -1.28 | 1.22E-06 |
| FBgn0031675 | 1.20 | 3.54E-12 | FBgn0041205 | -1.28 | 4.25E-06 |
| FBgn0026059 | 1.19 | 1.32E-18 | FBgn0030362 | -1.28 | 4.34E-09 |
| FBgn0267758 | 1.19 | 5.49E-06 | FBgn0038038 | -1.28 | 1.25E-08 |

| | | | | | |
|---|---|---|---|---|---|
| FBgn0045035 | 1.18 | 2.24E-05 | FBgn0035142 | -1.28 | 5.30E-06 |
| FBgn0003209 | 1.18 | 5.27E-14 | FBgn0035187 | -1.29 | 1.85E-05 |
| FBgn0262519 | 1.18 | 9.16E-24 | FBgn0267481 | -1.29 | 2.44E-07 |
| FBgn0024277 | 1.17 | 1.66E-05 | FBgn0037912 | -1.29 | 2.43E-09 |
| FBgn0031717 | 1.15 | 2.88E-14 | FBgn0025469 | -1.29 | 1.24E-06 |
| FBgn0013269 | 1.15 | 9.06E-09 | FBgn0037913 | -1.29 | 4.28E-09 |
| FBgn0040532 | 1.15 | 7.33E-18 | FBgn0031821 | -1.29 | 5.14E-06 |
| FBgn0034997 | 1.14 | 1.60E-06 | FBgn0039697 | -1.29 | 4.63E-09 |
| FBgn0005612 | 1.14 | 1.86E-09 | FBgn0042094 | -1.30 | 9.61E-09 |
| FBgn0042185 | 1.14 | 2.56E-09 | FBgn0033812 | -1.30 | 5.64E-06 |
| FBgn0030122 | 1.14 | 2.44E-06 | FBgn0032652 | -1.30 | 6.64E-07 |
| FBgn0020513 | 1.13 | 1.24E-11 | FBgn0266637 | -1.30 | 2.05E-07 |
| FBgn0043069 | 1.13 | 9.70E-08 | FBgn0038256 | -1.31 | 3.09E-07 |
| FBgn0001228 | 1.13 | 2.15E-08 | FBgn0264489 | -1.31 | 1.80E-05 |
| FBgn0035830 | 1.13 | 1.86E-18 | FBgn0039519 | -1.31 | 3.77E-05 |
| FBgn0001223 | 1.13 | 2.35E-08 | FBgn0028563 | -1.31 | 1.23E-07 |
| FBgn0000565 | 1.11 | 1.34E-06 | FBgn0024293 | -1.32 | 9.60E-06 |
| FBgn0037354 | 1.11 | 2.59E-06 | FBgn0040827 | -1.32 | 6.14E-07 |
| FBgn0029092 | 1.11 | 1.02E-07 | FBgn0025352 | -1.33 | 2.57E-07 |
| FBgn0050115 | 1.11 | 6.55E-05 | FBgn0027580 | -1.33 | 3.90E-07 |
| FBgn0038344 | 1.11 | 1.36E-11 | FBgn0086906 | -1.33 | 6.59E-05 |
| FBgn0024491 | 1.11 | 4.30E-07 | FBgn0030425 | -1.34 | 1.61E-06 |
| FBgn0029657 | 1.11 | 2.23E-06 | FBgn0039537 | -1.34 | 2.27E-09 |
| FBgn0033015 | 1.11 | 9.04E-08 | FBgn0013308 | -1.35 | 1.24E-08 |
| FBgn0039155 | 1.10 | 2.25E-14 | FBgn0031818 | -1.35 | 1.74E-08 |
| FBgn0000052 | 1.10 | 1.84E-08 | FBgn0035026 | -1.36 | 1.20E-08 |
| FBgn0052758 | 1.09 | 1.78E-06 | FBgn0053120 | -1.37 | 1.26E-08 |
| FBgn0031689 | 1.09 | 1.35E-07 | FBgn0026878 | -1.37 | 7.60E-08 |
| FBgn0022213 | 1.09 | 1.07E-10 | FBgn0035734 | -1.37 | 1.24E-05 |
| FBgn0031039 | 1.09 | 2.29E-07 | FBgn0033465 | -1.37 | 5.40E-10 |
| FBgn0031038 | 1.09 | 2.29E-07 | FBgn0027610 | -1.37 | 2.94E-05 |
| FBgn0025454 | 1.09 | 7.22E-05 | FBgn0028360 | -1.37 | 2.06E-05 |
| FBgn0263599 | 1.08 | 5.61E-13 | FBgn0031012 | -1.37 | 8.09E-07 |
| FBgn0085466 | 1.08 | 7.17E-09 | FBgn0010389 | -1.39 | 5.80E-05 |
| FBgn0033809 | 1.08 | 3.01E-12 | FBgn0051548 | -1.39 | 1.80E-07 |
| FBgn0027611 | 1.08 | 0.00011213 | FBgn0034629 | -1.39 | 1.91E-09 |
| FBgn0032923 | 1.08 | 2.03E-19 | FBgn0262782 | -1.39 | 3.92E-10 |
| FBgn0033320 | 1.08 | 5.41E-07 | FBgn0262881 | -1.40 | 5.86E-05 |
| FBgn0034072 | 1.08 | 2.64E-08 | FBgn0030575 | -1.40 | 6.18E-08 |
| FBgn0261574 | 1.07 | 4.00E-06 | FBgn0032913 | -1.40 | 5.56E-07 |
| FBgn0033000 | 1.06 | 1.97E-05 | FBgn0038516 | -1.41 | 8.81E-07 |
| FBgn0031114 | 1.06 | 4.91E-07 | FBgn0029994 | -1.42 | 1.29E-05 |
| FBgn0014133 | 1.06 | 2.81E-05 | FBgn0030932 | -1.42 | 6.53E-07 |
| FBgn0029167 | 1.06 | 8.00E-10 | FBgn0026570 | -1.42 | 1.24E-06 |
| FBgn0263996 | 1.05 | 1.04E-10 | FBgn0035186 | -1.42 | 1.77E-10 |
| FBgn0031432 | 1.04 | 7.92E-12 | FBgn0039051 | -1.43 | 3.22E-09 |
| FBgn0030838 | 1.04 | 3.10E-06 | FBgn0033112 | -1.43 | 1.35E-06 |
| FBgn0262656 | 1.04 | 5.23E-10 | FBgn0026150 | -1.43 | 1.16E-08 |
| FBgn0259152 | 1.04 | 2.21E-13 | FBgn0038928 | -1.45 | 1.22E-05 |
| FBgn0032805 | 1.04 | 1.68E-05 | FBgn0033366 | -1.46 | 1.32E-07 |
| FBgn0014141 | 1.04 | 3.79E-10 | FBgn0039696 | -1.46 | 9.87E-07 |
| FBgn0261811 | 1.04 | 7.72E-06 | FBgn0031069 | -1.46 | 2.73E-10 |
| FBgn0058469 | 1.03 | 3.04E-07 | FBgn0025631 | -1.46 | 2.12E-08 |
| FBgn0032901 | 1.02 | 1.80E-10 | FBgn0086758 | -1.47 | 1.55E-06 |
| FBgn0002645 | 1.01 | 4.05E-10 | FBgn0036273 | -1.47 | 6.89E-06 |
| FBgn0034184 | 1.01 | 0.00012018 | FBgn0020626 | -1.47 | 1.22E-08 |
| FBgn0030749 | 1.01 | 2.76E-11 | FBgn0029935 | -1.48 | 4.50E-07 |
| FBgn0016917 | 1.00 | 7.20E-11 | FBgn0013764 | -1.48 | 8.25E-07 |
| FBgn0022774 | 1.00 | 6.10E-11 | FBgn0033446 | -1.48 | 1.65E-06 |
| FBgn0026239 | 1.00 | 2.47E-06 | FBgn0040732 | -1.48 | 4.77E-05 |
| FBgn0010905 | 1.00 | 1.27E-08 | FBgn0283510 | -1.48 | 2.25E-07 |
| FBgn0050489 | 1.00 | 1.22E-06 | FBgn0034488 | -1.49 | 1.63E-07 |
| FBgn0028341 | 1.00 | 7.51E-06 | FBgn0037537 | -1.49 | 1.85E-08 |
| FBgn0033095 | 1.00 | 4.97E-07 | FBgn0261989 | -1.50 | 6.40E-06 |
| FBgn0266525 | 1.00 | 4.97E-07 | FBgn0034046 | -1.50 | 1.39E-07 |
| FBgn0031471 | 1.00 | 3.57E-05 | FBgn0038312 | -1.50 | 4.70E-10 |
| FBgn0033777 | 0.99 | 1.06E-09 | FBgn0017482 | -1.50 | 2.64E-11 |
| FBgn0034641 | 0.99 | 2.57E-09 | FBgn0039094 | -1.51 | 5.34E-08 |
| FBgn0036514 | 0.99 | 0.00012776 | FBgn0034628 | -1.51 | 3.60E-11 |
| FBgn0038020 | 0.99 | 6.83E-05 | FBgn0016053 | -1.52 | 1.94E-11 |
| FBgn0053494 | 0.99 | 1.67E-12 | FBgn0010387 | -1.52 | 4.39E-05 |
| FBgn0260749 | 0.99 | 0.00012166 | FBgn0039872 | -1.53 | 4.10E-06 |
| FBgn0032884 | 0.99 | 5.94E-10 | FBgn0031695 | -1.54 | 1.56E-07 |
| FBgn0261570 | 0.99 | 3.20E-07 | FBgn0284257 | -1.55 | 1.11E-11 |
| FBgn0010415 | 0.98 | 3.72E-16 | FBgn0051106 | -1.55 | 5.48E-05 |
| FBgn0053969 | 0.97 | 4.46E-09 | FBgn0052446 | -1.55 | 2.93E-09 |
| FBgn0005411 | 0.97 | 5.31E-11 | FBgn0264776 | -1.56 | 1.17E-07 |
| FBgn0286051 | 0.97 | 1.87E-05 | FBgn0029167 | -1.56 | 4.33E-05 |
| FBgn0262160 | 0.96 | 9.09E-06 | FBgn0031936 | -1.56 | 4.15E-08 |
| FBgn0035542 | 0.96 | 5.38E-11 | FBgn0033717 | -1.57 | 2.50E-08 |
| FBgn0087039 | 0.96 | 9.38E-06 | FBgn0034487 | -1.57 | 1.11E-06 |
| FBgn0037339 | 0.95 | 1.69E-10 | FBgn0026602 | -1.57 | 1.04E-07 |
| FBgn0040236 | 0.95 | 8.59E-08 | FBgn0036039 | -1.58 | 1.23E-10 |
| FBgn0029093 | 0.95 | 2.92E-13 | FBgn0035539 | -1.59 | 8.33E-07 |
| FBgn0264607 | 0.95 | 2.05E-08 | FBgn0038795 | -1.59 | 1.79E-06 |
| FBgn0029878 | 0.94 | 3.74E-06 | FBgn0010238 | -1.59 | 2.71E-05 |
| FBgn0053229 | 0.93 | 1.81E-12 | FBgn0265375 | -1.59 | 5.19E-08 |
| FBgn0032388 | 0.93 | 2.27E-09 | FBgn0015558 | -1.59 | 1.06E-07 |
| FBgn0001977 | 0.93 | 3.07E-08 | FBgn0031169 | -1.59 | 2.12E-07 |
| FBgn0031115 | 0.93 | 3.66E-08 | FBgn0035239 | -1.61 | 3.69E-05 |

| | | | | | |
|---|---|---|---|---|---|
| FBgn0031534 | 0.93 | 8.14E-05 | FBgn0035240 | -1.61 | 3.69E-05 |
| FBgn0038035 | 0.92 | 5.61E-11 | FBgn0031117 | -1.61 | 1.81E-10 |
| FBgn0020389 | 0.91 | 1.47E-07 | FBgn0029969 | -1.61 | 1.25E-09 |
| FBgn0038890 | 0.91 | 9.73E-05 | FBgn0028479 | -1.61 | 1.61E-12 |
| FBgn0036290 | 0.91 | 1.21E-08 | FBgn0033742 | -1.62 | 1.32E-11 |
| FBgn0039302 | 0.91 | 1.75E-09 | FBgn0030160 | -1.62 | 5.22E-12 |
| FBgn0033133 | 0.91 | 6.67E-07 | FBgn0016715 | -1.62 | 4.93E-06 |
| FBgn0031461 | 0.90 | 7.52E-07 | FBgn0036822 | -1.62 | 2.95E-08 |
| FBgn0262109 | 0.90 | 8.80E-16 | FBgn0033782 | -1.63 | 2.72E-11 |
| FBgn0035236 | 0.90 | 2.45E-06 | FBgn0032946 | -1.64 | 5.21E-05 |
| FBgn0010909 | 0.90 | 9.12E-08 | FBgn0003892 | -1.64 | 4.12E-06 |
| FBgn0004449 | 0.90 | 1.65E-05 | FBgn0039152 | -1.67 | 4.74E-11 |
| FBgn0260933 | 0.90 | 4.03E-08 | FBgn0051472 | -1.67 | 9.03E-12 |
| FBgn0030740 | 0.90 | 7.22E-05 | FBgn0039157 | -1.68 | 3.76E-13 |
| FBgn0027329 | 0.89 | 0.00013549 | FBgn0033128 | -1.69 | 9.43E-06 |
| FBgn0031263 | 0.89 | 7.49E-08 | FBgn0030731 | -1.69 | 1.87E-14 |
| FBgn0261793 | 0.88 | 7.04E-08 | FBgn0035132 | -1.70 | 1.93E-06 |
| FBgn0027568 | 0.88 | 2.53E-07 | FBgn0035169 | -1.70 | 4.98E-14 |
| FBgn0037556 | 0.88 | 2.95E-05 | FBgn0042627 | -1.70 | 5.13E-05 |
| FBgn0264691 | 0.88 | 1.04E-05 | FBgn0033232 | -1.72 | 8.84E-13 |
| FBgn0023143 | 0.88 | 3.88E-09 | FBgn0036824 | -1.72 | 6.48E-11 |
| FBgn0035499 | 0.88 | 8.47E-15 | FBgn0066101 | -1.73 | 1.25E-06 |
| FBgn0037363 | 0.87 | 4.82E-12 | FBgn0040813 | -1.74 | 4.68E-05 |
| FBgn0029506 | 0.87 | 3.59E-08 | FBgn0031478 | -1.74 | 1.91E-13 |
| FBgn0022987 | 0.86 | 6.35E-06 | FBgn0037936 | -1.74 | 7.81E-08 |
| FBgn0051974 | 0.86 | 4.16E-05 | FBgn0025885 | -1.74 | 2.50E-08 |
| FBgn0004648 | 0.86 | 2.09E-14 | FBgn0032538 | -1.74 | 9.11E-06 |
| FBgn0033725 | 0.85 | 1.60E-05 | FBgn0032943 | -1.75 | 5.21E-09 |
| FBgn0000568 | 0.85 | 5.92E-05 | FBgn0005666 | -1.76 | 2.18E-11 |
| FBgn0036875 | 0.85 | 2.15E-07 | FBgn0034063 | -1.76 | 2.40E-05 |
| FBgn0036921 | 0.85 | 9.03E-09 | FBgn0012034 | -1.78 | 6.19E-14 |
| FBgn0028399 | 0.85 | 5.80E-06 | FBgn0034638 | -1.79 | 2.54E-06 |
| FBgn0259824 | 0.84 | 1.77E-07 | FBgn0034440 | -1.80 | 4.52E-06 |
| FBgn0031389 | 0.84 | 7.98E-05 | FBgn0028544 | -1.81 | 1.18E-05 |
| FBgn0034398 | 0.84 | 3.06E-12 | FBgn0052391 | -1.81 | 3.42E-05 |
| FBgn0003996 | 0.83 | 1.94E-05 | FBgn0027660 | -1.82 | 5.60E-08 |
| FBgn0036915 | 0.83 | 5.67E-06 | FBgn0034479 | -1.83 | 1.38E-10 |
| FBgn0021768 | 0.83 | 5.18E-06 | FBgn0011754 | -1.83 | 1.93E-14 |
| FBgn0038034 | 0.82 | 2.55E-05 | FBgn0261446 | -1.85 | 5.89E-11 |
| FBgn0027363 | 0.82 | 3.31E-05 | FBgn0033519 | -1.85 | 6.89E-07 |
| FBgn0036980 | 0.82 | 9.29E-09 | FBgn0030607 | -1.86 | 9.13E-14 |
| FBgn0002643 | 0.82 | 3.11E-07 | FBgn0039494 | -1.86 | 1.03E-11 |
| FBgn0036173 | 0.82 | 8.19E-05 | FBgn0029769 | -1.87 | 9.02E-07 |
| FBgn0020622 | 0.81 | 4.43E-10 | FBgn0039431 | -1.88 | 3.48E-10 |
| FBgn0002566 | 0.80 | 1.74E-06 | FBgn0260866 | -1.89 | 7.53E-12 |
| FBgn0010488 | 0.80 | 2.08E-06 | FBgn0026593 | -1.89 | 7.74E-08 |
| FBgn0029996 | 0.80 | 1.65E-05 | FBgn0032264 | -1.90 | 3.26E-06 |
| FBgn0051370 | 0.79 | 6.36E-06 | FBgn0037996 | -1.90 | 3.50E-05 |
| FBgn0264357 | 0.79 | 2.61E-11 | FBgn0034141 | -1.91 | 2.20E-08 |
| FBgn0028572 | 0.79 | 1.09E-07 | FBgn0015609 | -1.91 | 8.66E-14 |
| FBgn0030246 | 0.79 | 1.97E-05 | FBgn0035262 | -1.92 | 4.65E-08 |
| FBgn0260962 | 0.78 | 1.78E-07 | FBgn0037853 | -1.93 | 4.34E-07 |
| FBgn0039691 | 0.78 | 6.63E-06 | FBgn0000018 | -1.93 | 4.42E-07 |
| FBgn0036828 | 0.78 | 7.45E-05 | FBgn0027866 | -1.95 | 3.97E-13 |
| FBgn0039944 | 0.78 | 3.69E-06 | FBgn0030966 | -1.95 | 6.16E-13 |
| FBgn0039945 | 0.77 | 4.30E-06 | FBgn0013772 | -1.95 | 5.91E-10 |
| FBgn0030521 | 0.77 | 0.00011967 | FBgn0037167 | -1.96 | 2.60E-08 |
| FBgn0065081 | 0.76 | 1.57E-06 | FBgn0063924 | -1.96 | 1.34E-07 |
| FBgn0041194 | 0.75 | 1.07E-09 | FBgn0265767 | -1.96 | 7.70E-13 |
| FBgn0031968 | 0.75 | 0.00012919 | FBgn0011020 | -1.97 | 2.80E-05 |
| FBgn0045823 | 0.75 | 5.08E-08 | FBgn0036205 | -1.97 | 4.08E-10 |
| FBgn0039120 | 0.74 | 6.29E-08 | FBgn0050485 | -1.98 | 6.34E-13 |
| FBgn0028424 | 0.73 | 9.44E-05 | FBgn0040507 | -1.99 | 3.52E-08 |
| FBgn0010288 | 0.73 | 1.02E-05 | FBgn0033879 | -1.99 | 3.50E-15 |
| FBgn0019662 | 0.73 | 4.65E-05 | FBgn0259714 | -2.01 | 2.59E-08 |
| FBgn0035229 | 0.72 | 5.19E-05 | FBgn0260858 | -2.02 | 6.84E-17 |
| FBgn0037518 | 0.72 | 6.83E-07 | FBgn0034712 | -2.03 | 2.14E-08 |
| FBgn0004811 | 0.71 | 6.79E-06 | FBgn0260658 | -2.04 | 1.41E-08 |
| FBgn0030748 | 0.71 | 5.92E-06 | FBgn0003067 | -2.06 | 7.35E-08 |
| FBgn0036448 | 0.71 | 6.36E-05 | FBgn0266488 | -2.06 | 7.35E-08 |
| FBgn0032208 | 0.70 | 2.63E-10 | FBgn0039156 | -2.07 | 2.99E-20 |
| FBgn0266019 | 0.69 | 9.54E-09 | FBgn0065032 | -2.07 | 3.64E-13 |
| FBgn0052676 | 0.68 | 2.27E-05 | FBgn0035833 | -2.08 | 6.40E-05 |
| FBgn0028577 | 0.68 | 9.51E-05 | FBgn0030701 | -2.09 | 1.95E-13 |
| FBgn0014029 | 0.66 | 3.52E-05 | FBgn0264950 | -2.09 | 6.02E-11 |
| FBgn0039113 | 0.66 | 5.53E-05 | FBgn0031974 | -2.10 | 2.48E-13 |
| FBgn0036685 | 0.66 | 1.06E-05 | FBgn0053056 | -2.11 | 2.33E-18 |
| FBgn0036556 | 0.65 | 8.44E-05 | FBgn0052311 | -2.11 | 2.64E-06 |
| FBgn0035871 | 0.65 | 2.62E-08 | FBgn0031998 | -2.12 | 4.46E-12 |
| FBgn0052672 | 0.65 | 4.79E-05 | FBgn0037750 | -2.14 | 5.66E-10 |
| FBgn0024285 | 0.65 | 1.53E-05 | FBgn0034053 | -2.14 | 1.92E-08 |
| FBgn0261456 | 0.64 | 0.00013116 | FBgn0039737 | -2.14 | 2.63E-22 |
| FBgn0052369 | 0.63 | 2.93E-05 | FBgn0032715 | -2.17 | 9.72E-20 |
| FBgn0037553 | 0.63 | 4.95E-05 | FBgn0267804 | -2.17 | 2.16E-21 |
| FBgn0032456 | 0.63 | 2.02E-08 | FBgn0038332 | -2.19 | 2.93E-16 |
| FBgn0038826 | 0.62 | 2.46E-06 | FBgn0051373 | -2.19 | 9.10E-14 |
| FBgn0036702 | 0.62 | 1.02E-05 | FBgn0034493 | -2.19 | 1.28E-09 |
| FBgn0050273 | 0.62 | 0.00011828 | FBgn0015570 | -2.24 | 5.99E-23 |
| FBgn0050269 | 0.62 | 0.00011828 | FBgn0033397 | -2.29 | 2.29E-09 |
| FBgn0031969 | 0.62 | 2.56E-05 | FBgn0038682 | -2.31 | 7.80E-06 |

| | | | | | |
|---|---|---|---|---|---|
| FBgn0033669 | 0.61 | 5.69E-05 | FBgn0040823 | -2.31 | 2.60E-05 |
| FBgn0039178 | 0.60 | 1.35E-05 | FBgn0036749 | -2.33 | 1.09E-10 |
| FBgn0284421 | 0.60 | 5.40E-05 | FBgn0025720 | -2.35 | 1.54E-09 |
| FBgn0260794 | 0.60 | 0.00011186 | FBgn0038148 | -2.36 | 1.68E-05 |
| FBgn0050359 | 0.58 | 6.91E-06 | FBgn0039938 | -2.36 | 2.62E-05 |
| FBgn0036534 | 0.58 | 1.41E-06 | FBgn0030013 | -2.37 | 9.85E-26 |
| FBgn0265048 | 0.55 | 2.11E-07 | FBgn0033830 | -2.38 | 1.01E-07 |
| FBgn0033226 | 0.55 | 5.36E-05 | FBgn0031064 | -2.38 | 1.61E-05 |
| FBgn0030263 | 0.55 | 4.71E-05 | FBgn0032010 | -2.39 | 1.13E-06 |
| FBgn0004638 | 0.54 | 4.60E-05 | FBgn0035517 | -2.39 | 3.35E-19 |
| FBgn0267398 | 0.54 | 2.46E-05 | FBgn0038674 | -2.40 | 1.80E-10 |
| FBgn0030809 | 0.53 | 3.51E-05 | FBgn0259243 | -2.41 | 2.60E-21 |
| FBgn0038037 | 0.51 | 2.46E-05 | FBgn0038039 | -2.42 | 6.80E-23 |
| FBgn0082598 | 0.51 | 3.07E-06 | FBgn0034710 | -2.43 | 1.19E-13 |
| FBgn0261671 | 0.51 | 5.40E-05 | FBgn0021765 | -2.45 | 3.27E-23 |
| FBgn0036337 | 0.49 | 2.54E-05 | FBgn0051313 | -2.45 | 1.15E-20 |
| FBgn0266410 | 0.49 | 3.20E-05 | FBgn0001970 | -2.45 | 4.24E-14 |
| FBgn0025741 | 0.49 | 8.78E-06 | FBgn0266428 | -2.46 | 3.48E-14 |
| FBgn0032216 | 0.48 | 6.12E-05 | FBgn0265045 | -2.47 | 2.01E-22 |
| FBgn0020309 | 0.44 | 0.00010128 | FBgn0035880 | -2.49 | 3.03E-08 |
| FBgn0259224 | -0.46 | 9.33E-05 | FBgn0035309 | -2.50 | 1.30E-15 |
| FBgn0002526 | -0.47 | 5.85E-05 | FBgn0261564 | -2.50 | 7.34E-27 |
| FBgn0026428 | -0.49 | 5.93E-07 | FBgn0041712 | -2.51 | 1.74E-16 |
| FBgn0261592 | -0.52 | 0.00010473 | FBgn0039102 | -2.52 | 1.62E-26 |
| FBgn0020385 | -0.52 | 1.57E-05 | FBgn0032178 | -2.52 | 2.86E-07 |
| FBgn0001942 | -0.56 | 1.26E-06 | FBgn0010504 | -2.53 | 1.22E-23 |
| FBgn0040398 | -0.56 | 5.63E-05 | FBgn0050457 | -2.54 | 4.77E-05 |
| FBgn0031092 | -0.56 | 2.82E-05 | FBgn0039098 | -2.55 | 1.08E-17 |
| FBgn0033402 | -0.57 | 8.58E-05 | FBgn0030484 | -2.55 | 1.83E-10 |
| FBgn0037146 | -0.58 | 5.00E-06 | FBgn0032167 | -2.55 | 4.17E-24 |
| FBgn0031126 | -0.59 | 1.93E-05 | FBgn0038242 | -2.55 | 6.84E-12 |
| FBgn0021979 | -0.59 | 8.77E-06 | FBgn0264507 | -2.57 | 7.15E-21 |
| FBgn0263594 | -0.59 | 2.09E-06 | FBgn0033872 | -2.59 | 4.81E-06 |
| FBgn0285991 | -0.59 | 7.11E-06 | FBgn0085399 | -2.59 | 3.82E-06 |
| FBgn0032147 | -0.59 | 1.65E-06 | FBgn0264916 | -2.60 | 8.18E-07 |
| FBgn0263598 | -0.59 | 0.0001102 | FBgn0023513 | -2.60 | 1.19E-24 |
| FBgn0025352 | -0.60 | 2.82E-05 | FBgn0038610 | -2.63 | 2.24E-07 |
| FBgn0039260 | -0.60 | 8.21E-05 | FBgn0029791 | -2.65 | 4.87E-27 |
| FBgn0036258 | -0.60 | 1.54E-07 | FBgn0030769 | -2.65 | 1.08E-09 |
| FBgn0034876 | -0.61 | 4.21E-05 | FBgn0033395 | -2.67 | 1.27E-05 |
| FBgn0026199 | -0.61 | 0.00011896 | FBgn0085433 | -2.68 | 7.86E-15 |
| FBgn0016756 | -0.61 | 4.46E-06 | FBgn0002773 | -2.68 | 3.09E-12 |
| FBgn0052479 | -0.62 | 4.51E-05 | FBgn0038331 | -2.70 | 6.86E-29 |
| FBgn0285910 | -0.62 | 0.00012471 | FBgn0032211 | -2.70 | 1.66E-05 |
| FBgn0011284 | -0.62 | 0.00012325 | FBgn0036352 | -2.70 | 7.06E-06 |
| FBgn0051116 | -0.62 | 3.40E-05 | FBgn0033542 | -2.75 | 6.56E-06 |
| FBgn0030327 | -0.63 | 1.21E-05 | FBgn0032891 | -2.75 | 1.03E-15 |
| FBgn0031598 | -0.63 | 9.58E-05 | FBgn0032629 | -2.76 | 1.51E-08 |
| FBgn0040732 | -0.64 | 1.16E-09 | FBgn0262816 | -2.80 | 2.87E-05 |
| FBgn0016701 | -0.64 | 2.70E-05 | FBgn0035839 | -2.82 | 1.28E-20 |
| FBgn0085208 | -0.66 | 3.62E-05 | FBgn0053138 | -2.95 | 7.53E-21 |
| FBgn0029161 | -0.66 | 3.66E-05 | FBgn0267035 | -2.95 | 2.08E-28 |
| FBgn0267384 | -0.66 | 4.63E-05 | FBgn0017561 | -2.96 | 6.11E-22 |
| FBgn0015808 | -0.66 | 2.09E-05 | FBgn0036351 | -2.97 | 7.37E-06 |
| FBgn0031359 | -0.67 | 1.91E-06 | FBgn0015777 | -2.99 | 4.56E-30 |
| FBgn0011336 | -0.67 | 9.11E-05 | FBgn0264384 | -3.00 | 7.23E-22 |
| FBgn0027594 | -0.67 | 2.93E-08 | FBgn0041245 | -3.01 | 1.45E-05 |
| FBgn0033483 | -0.67 | 3.75E-08 | FBgn0030073 | -3.02 | 6.69E-24 |
| FBgn0053120 | -0.67 | 2.13E-05 | FBgn0036353 | -3.02 | 5.04E-35 |
| FBgn0019960 | -0.67 | 1.29E-05 | FBgn0023541 | -3.03 | 3.62E-29 |
| FBgn0004888 | -0.68 | 4.09E-05 | FBgn0035552 | -3.04 | 4.55E-07 |
| FBgn0011642 | -0.68 | 0.00013625 | FBgn0036348 | -3.07 | 1.13E-07 |
| FBgn0034718 | -0.69 | 5.95E-05 | FBgn0037163 | -3.12 | 4.28E-30 |
| FBgn0036929 | -0.69 | 0.00010027 | FBgn0053474 | -3.13 | 7.66E-08 |
| FBgn0039303 | -0.69 | 3.50E-05 | FBgn0266446 | -3.14 | 8.69E-14 |
| FBgn0034951 | -0.69 | 5.20E-05 | FBgn0267753 | -3.17 | 2.01E-05 |
| FBgn0000100 | -0.69 | 1.91E-05 | FBgn0045479 | -3.19 | 1.77E-05 |
| FBgn0027885 | -0.69 | 7.03E-06 | FBgn0041150 | -3.22 | 9.76E-09 |
| FBgn0035321 | -0.69 | 1.16E-05 | FBgn0051493 | -3.23 | 6.91E-06 |
| FBgn0033453 | -0.69 | 2.58E-05 | FBgn0038149 | -3.26 | 1.46E-26 |
| FBgn0004876 | -0.70 | 3.40E-05 | FBgn0035344 | -3.26 | 1.08E-16 |
| FBgn0262782 | -0.70 | 1.29E-07 | FBgn0041707 | -3.31 | 3.12E-14 |
| FBgn0050431 | -0.70 | 4.17E-05 | FBgn0005636 | -3.32 | 8.23E-15 |
| FBgn0034117 | -0.70 | 0.0001093 | FBgn0035241 | -3.32 | 4.19E-17 |
| FBgn0034753 | -0.71 | 6.07E-06 | FBgn0028482 | -3.37 | 1.20E-12 |
| FBgn0052521 | -0.71 | 8.87E-06 | FBgn0026061 | -3.46 | 1.49E-26 |
| FBgn0032480 | -0.71 | 1.07E-06 | FBgn0264269 | -3.49 | 2.18E-06 |
| FBgn0259993 | -0.72 | 2.20E-06 | FBgn0034279 | -3.53 | 1.12E-18 |
| FBgn0033751 | -0.72 | 8.38E-06 | FBgn0085359 | -3.56 | 1.33E-26 |
| FBgn0039611 | -0.72 | 3.61E-05 | FBgn0010482 | -3.58 | 3.51E-18 |
| FBgn0022710 | -0.72 | 6.12E-05 | FBgn0051559 | -3.61 | 1.04E-07 |
| FBgn0031805 | -0.72 | 4.13E-05 | FBgn0261053 | -3.66 | 2.94E-14 |
| FBgn0015575 | -0.73 | 4.86E-05 | FBgn0038097 | -3.67 | 4.45E-13 |
| FBgn0026313 | -0.73 | 5.44E-08 | FBgn0033179 | -3.67 | 1.03E-46 |
| FBgn0031453 | -0.73 | 2.36E-12 | FBgn0030326 | -3.73 | 2.82E-20 |
| FBgn0010100 | -0.73 | 8.12E-06 | FBgn0050297 | -3.76 | 1.05E-07 |
| FBgn0034087 | -0.73 | 2.81E-07 | FBgn0034724 | -3.77 | 1.29E-18 |
| FBgn0020910 | -0.73 | 2.99E-07 | FBgn0085414 | -3.82 | 5.12E-12 |
| FBgn0267978 | -0.73 | 6.49E-05 | FBgn0029518 | -3.84 | 1.11E-20 |
| FBgn0036752 | -0.74 | 7.92E-06 | FBgn0032945 | -3.84 | 1.60E-11 |

| | | |
|---|---|---|
| FBgn0037186 | -0.74 | 3.72E-08 |
| FBgn0028476 | -0.75 | 3.20E-06 |
| FBgn0051144 | -0.75 | 8.99E-08 |
| FBgn0028737 | -0.75 | 9.92E-05 |
| FBgn0013749 | -0.75 | 1.93E-05 |
| FBgn0265901 | -0.76 | 0.00013294 |
| FBgn0028956 | -0.76 | 1.35E-06 |
| FBgn0015040 | -0.76 | 1.76E-05 |
| FBgn0264328 | -0.76 | 1.76E-05 |
| FBgn0262570 | -0.77 | 7.48E-05 |
| FBgn0030026 | -0.77 | 8.69E-05 |
| FBgn0010591 | -0.77 | 5.12E-08 |
| FBgn0029688 | -0.77 | 3.42E-05 |
| FBgn0030262 | -0.77 | 1.77E-09 |
| FBgn0035383 | -0.78 | 4.55E-06 |
| FBgn0025381 | -0.78 | 7.19E-05 |
| FBgn0014469 | -0.78 | 7.67E-05 |
| FBgn0003071 | -0.79 | 2.29E-05 |
| FBgn0267481 | -0.79 | 6.56E-05 |
| FBgn0031897 | -0.79 | 6.90E-06 |
| FBgn0029737 | -0.79 | 5.84E-06 |
| FBgn0041191 | -0.79 | 6.10E-08 |
| FBgn0250848 | -0.80 | 2.41E-07 |
| FBgn0044324 | -0.80 | 1.10E-06 |
| FBgn0015288 | -0.80 | 8.20E-12 |
| FBgn0259735 | -0.80 | 2.74E-06 |
| FBgn0034277 | -0.80 | 0.00011518 |
| FBgn0001186 | -0.80 | 5.27E-08 |
| FBgn0003517 | -0.80 | 7.19E-06 |
| FBgn0011770 | -0.81 | 3.37E-05 |
| FBgn0035111 | -0.81 | 3.57E-06 |
| FBgn0037875 | -0.81 | 8.29E-05 |
| FBgn0001125 | -0.81 | 2.84E-14 |
| FBgn0005771 | -0.81 | 6.87E-05 |
| FBgn0023441 | -0.81 | 7.70E-08 |
| FBgn0023172 | -0.81 | 5.82E-07 |
| FBgn0038291 | -0.81 | 1.85E-06 |
| FBgn0037817 | -0.81 | 7.59E-08 |
| FBgn0027569 | -0.81 | 6.37E-05 |
| FBgn0034066 | -0.82 | 1.61E-06 |
| FBgn0267160 | -0.82 | 1.67E-06 |
| FBgn0020415 | -0.82 | 5.28E-06 |
| FBgn0264294 | -0.82 | 4.93E-05 |
| FBgn0261836 | -0.82 | 1.88E-08 |
| FBgn0266758 | -0.83 | 3.54E-06 |
| FBgn0013763 | -0.83 | 9.69E-06 |
| FBgn0031950 | -0.83 | 1.08E-05 |
| FBgn0032881 | -0.83 | 8.27E-06 |
| FBgn0011584 | -0.83 | 1.96E-06 |
| FBgn0031174 | -0.83 | 3.03E-05 |
| FBgn0033812 | -0.83 | 2.23E-08 |
| FBgn0032075 | -0.83 | 8.53E-08 |
| FBgn0266729 | -0.84 | 4.56E-05 |
| FBgn0086371 | -0.84 | 6.13E-08 |
| FBgn0038947 | -0.84 | 1.46E-05 |
| FBgn0032455 | -0.84 | 2.78E-07 |
| FBgn0023516 | -0.84 | 5.28E-05 |
| FBgn0038463 | -0.84 | 2.31E-05 |
| FBgn0039466 | -0.85 | 1.16E-05 |
| FBgn0031987 | -0.85 | 5.27E-05 |
| FBgn0030724 | -0.85 | 6.24E-05 |
| FBgn0266369 | -0.85 | 3.64E-09 |
| FBgn0031183 | -0.85 | 3.04E-05 |
| FBgn0284222 | -0.86 | 0.00011408 |
| FBgn0267505 | -0.86 | 1.21E-06 |
| FBgn0034989 | -0.86 | 1.07E-05 |
| FBgn0086679 | -0.86 | 4.71E-05 |
| FBgn0030797 | -0.86 | 9.72E-05 |
| FBgn0032224 | -0.86 | 2.94E-05 |
| FBgn0039358 | -0.86 | 1.45E-06 |
| FBgn0261625 | -0.86 | 5.96E-05 |
| FBgn0026415 | -0.86 | 0.00012444 |
| FBgn0033339 | -0.86 | 4.55E-06 |
| FBgn0085802 | -0.87 | 8.06E-07 |
| FBgn0036450 | -0.87 | 5.05E-06 |
| FBgn0035142 | -0.88 | 1.56E-08 |
| FBgn0087002 | -0.88 | 3.47E-06 |
| FBgn0042094 | -0.88 | 6.60E-05 |
| FBgn0027493 | -0.89 | 2.53E-05 |
| FBgn0039450 | -0.89 | 9.37E-06 |
| FBgn0026630 | -0.89 | 3.05E-06 |
| FBgn0283510 | -0.89 | 1.97E-11 |
| FBgn0037548 | -0.89 | 3.06E-05 |
| FBgn0024973 | -0.90 | 5.41E-07 |
| FBgn0050159 | -0.90 | 9.91E-06 |
| FBgn0000064 | -0.90 | 9.65E-05 |
| FBgn0031836 | -0.90 | 3.19E-07 |
| FBgn0027583 | -0.90 | 0.0001037 |
| FBgn0037635 | -0.90 | 4.88E-06 |
| FBgn0016715 | -0.90 | 4.17E-12 |

| | | |
|---|---|---|
| FBgn0041105 | -3.86 | 1.08E-07 |
| FBgn0031971 | -3.90 | 5.17E-23 |
| FBgn0022709 | -4.00 | 1.40E-30 |
| FBgn0029091 | -4.04 | 9.06E-36 |
| FBgn0037405 | -4.07 | 1.20E-13 |
| FBgn0035770 | -4.07 | 1.27E-42 |
| FBgn0265981 | -4.18 | 5.55E-09 |
| FBgn0005586 | -4.18 | 1.11E-11 |
| FBgn0266753 | -4.22 | 3.55E-09 |
| FBgn0037690 | -4.31 | 6.89E-11 |
| FBgn0023001 | -4.40 | 4.65E-53 |
| FBgn0020372 | -4.52 | 9.33E-47 |
| FBgn0038295 | -4.57 | 4.65E-13 |
| FBgn0011828 | -4.59 | 6.55E-19 |
| FBgn0030816 | -4.65 | 1.72E-48 |
| FBgn0086915 | -4.65 | 1.66E-31 |
| FBgn0031261 | -4.71 | 1.56E-19 |
| FBgn0005660 | -4.71 | 3.30E-21 |
| FBgn0267220 | -4.93 | 6.10E-30 |
| FBgn0266402 | -5.10 | 3.39E-14 |
| FBgn0028400 | -5.22 | 1.05E-14 |
| FBgn0264877 | -5.30 | 1.29E-35 |
| FBgn0033129 | -5.35 | 1.12E-42 |
| FBgn0050374 | -5.37 | 9.45E-30 |
| FBgn0039486 | -5.37 | 2.59E-34 |
| FBgn0266591 | -5.42 | 3.78E-22 |
| FBgn0264866 | -5.47 | 7.76E-17 |
| FBgn0267441 | -5.47 | 7.76E-17 |
| FBgn0045478 | -5.70 | 1.13E-22 |
| FBgn0045477 | -5.70 | 1.13E-22 |
| FBgn0037975 | -5.75 | 2.18E-29 |
| FBgn0019929 | -5.91 | 1.13E-43 |
| FBgn0038095 | -5.96 | 4.79E-50 |
| FBgn0052255 | -5.98 | 1.05E-25 |
| FBgn0027106 | -6.27 | 6.59E-123 |
| FBgn0085428 | -6.31 | 1.74E-80 |
| FBgn0034725 | -6.39 | 1.50E-24 |
| FBgn0045476 | -6.48 | 3.97E-30 |
| FBgn0035486 | -6.50 | 2.19E-30 |
| FBgn0035673 | -6.65 | 1.24E-64 |
| FBgn0262593 | -6.79 | 6.09E-72 |
| FBgn0001090 | -7.84 | 2.02E-66 |
| FBgn0030817 | -8.31 | 2.01E-76 |
| FBgn0266430 | -8.36 | 1.55E-55 |

| | | |
|---|---|---|
| FBgn0030056 | -0.91 | 1.64E-05 |
| FBgn0015039 | -0.91 | 2.78E-06 |
| FBgn0030575 | -0.91 | 3.94E-07 |
| FBgn0261015 | -0.91 | 1.03E-07 |
| FBgn0046685 | -0.91 | 7.30E-05 |
| FBgn0027945 | -0.91 | 1.90E-08 |
| FBgn0046874 | -0.92 | 9.37E-08 |
| FBgn0026090 | -0.92 | 8.39E-05 |
| FBgn0040383 | -0.92 | 2.73E-07 |
| FBgn0028327 | -0.92 | 2.60E-05 |
| FBgn0037239 | -0.92 | 7.53E-10 |
| FBgn0015737 | -0.92 | 6.21E-11 |
| FBgn0031148 | -0.92 | 1.11E-06 |
| FBgn0031771 | -0.92 | 2.43E-07 |
| FBgn0040309 | -0.92 | 1.89E-05 |
| FBgn0002563 | -0.93 | 3.13E-05 |
| FBgn0036537 | -0.93 | 1.13E-07 |
| FBgn0001325 | -0.93 | 9.02E-05 |
| FBgn0028509 | -0.94 | 3.16E-05 |
| FBgn0029818 | -0.94 | 5.84E-07 |
| FBgn0015776 | -0.94 | 1.57E-06 |
| FBgn0033982 | -0.94 | 8.35E-07 |
| FBgn0021944 | -0.95 | 4.31E-05 |
| FBgn0034488 | -0.95 | 4.65E-11 |
| FBgn0000562 | -0.95 | 1.85E-09 |
| FBgn0261532 | -0.95 | 7.20E-05 |
| FBgn0010350 | -0.95 | 1.15E-14 |
| FBgn0040305 | -0.95 | 4.73E-09 |
| FBgn0035945 | -0.96 | 5.30E-05 |
| FBgn0001091 | -0.96 | 8.62E-06 |
| FBgn0028658 | -0.96 | 6.84E-05 |
| FBgn0033744 | -0.97 | 7.35E-09 |
| FBgn0041205 | -0.97 | 2.48E-05 |
| FBgn0039187 | -0.97 | 1.25E-09 |
| FBgn0011016 | -0.98 | 2.72E-05 |
| FBgn0031436 | -0.98 | 8.34E-05 |
| FBgn0030734 | -0.98 | 9.29E-08 |
| FBgn0027538 | -0.98 | 2.82E-05 |
| FBgn0037973 | -0.98 | 4.67E-06 |
| FBgn0034354 | -0.98 | 3.46E-05 |
| FBgn0040234 | -0.98 | 2.47E-06 |
| FBgn0011225 | -0.98 | 1.32E-05 |
| FBgn0004396 | -0.98 | 6.68E-14 |
| FBgn0267743 | -0.99 | 1.27E-05 |
| FBgn0040349 | -0.99 | 6.87E-06 |
| FBgn0027607 | -0.99 | 2.66E-12 |
| FBgn0014028 | -0.99 | 4.07E-08 |
| FBgn0031937 | -0.99 | 1.75E-06 |
| FBgn0035186 | -0.99 | 3.50E-06 |
| FBgn0030087 | -0.99 | 7.21E-10 |
| FBgn0032161 | -1.00 | 8.39E-05 |
| FBgn0012036 | -1.00 | 2.58E-05 |
| FBgn0027108 | -1.00 | 9.09E-07 |
| FBgn0027844 | -1.00 | 3.60E-05 |
| FBgn0028670 | -1.00 | 3.78E-07 |
| FBgn0030718 | -1.00 | 6.76E-07 |
| FBgn0017567 | -1.00 | 5.36E-06 |
| FBgn0036787 | -1.00 | 1.22E-05 |
| FBgn0051769 | -1.01 | 2.25E-05 |
| FBgn0031360 | -1.01 | 1.62E-05 |
| FBgn0035978 | -1.01 | 5.33E-11 |
| FBgn0033978 | -1.01 | 1.22E-08 |
| FBgn0034494 | -1.02 | 1.89E-07 |
| FBgn0039993 | -1.02 | 7.38E-06 |
| FBgn0053129 | -1.02 | 1.11E-07 |
| FBgn0033458 | -1.02 | 1.47E-05 |
| FBgn0032160 | -1.02 | 4.53E-07 |
| FBgn0037955 | -1.03 | 5.52E-08 |
| FBgn0028526 | -1.03 | 3.13E-07 |
| FBgn0021967 | -1.03 | 7.10E-05 |
| FBgn0031021 | -1.03 | 8.57E-06 |
| FBgn0033401 | -1.03 | 4.90E-06 |
| FBgn0037913 | -1.03 | 1.28E-06 |
| FBgn0011361 | -1.03 | 4.39E-05 |
| FBgn0264494 | -1.04 | 6.39E-08 |
| FBgn0062442 | -1.04 | 8.82E-06 |
| FBgn0261108 | -1.04 | 8.19E-09 |
| FBgn0053307 | -1.04 | 3.89E-05 |
| FBgn0039877 | -1.04 | 7.15E-06 |
| FBgn0051673 | -1.04 | 3.69E-06 |
| FBgn0003076 | -1.04 | 9.96E-07 |
| FBgn0033783 | -1.05 | 3.52E-12 |
| FBgn0086676 | -1.05 | 1.15E-05 |
| FBgn0030853 | -1.05 | 7.55E-07 |
| FBgn0032034 | -1.05 | 7.09E-05 |
| FBgn0033543 | -1.05 | 4.55E-08 |
| FBgn0034902 | -1.05 | 1.54E-08 |
| FBgn0016031 | -1.05 | 3.23E-17 |
| FBgn0021795 | -1.06 | 1.51E-05 |
| FBgn0010808 | -1.06 | 6.05E-07 |

| | | |
|---|---|---|
| FBgn0028479 | -1.06 | 1.01E-10 |
| FBgn0038038 | -1.06 | 3.13E-06 |
| FBgn0036623 | -1.06 | 2.20E-10 |
| FBgn0017429 | -1.07 | 5.96E-05 |
| FBgn0034585 | -1.07 | 6.59E-06 |
| FBgn0039118 | -1.07 | 0.00010756 |
| FBgn0037912 | -1.07 | 2.75E-07 |
| FBgn0030362 | -1.07 | 5.88E-06 |
| FBgn0264489 | -1.07 | 1.04E-05 |
| FBgn0026602 | -1.07 | 8.62E-09 |
| FBgn0037749 | -1.07 | 9.53E-06 |
| FBgn0030968 | -1.07 | 6.04E-15 |
| FBgn0263391 | -1.08 | 3.75E-06 |
| FBgn0026403 | -1.08 | 5.96E-10 |
| FBgn0037646 | -1.08 | 5.56E-05 |
| FBgn0030675 | -1.08 | 2.41E-05 |
| FBgn0267823 | -1.08 | 5.78E-05 |
| FBgn0031834 | -1.08 | 3.25E-06 |
| FBgn0039098 | -1.09 | 6.72E-09 |
| FBgn0086357 | -1.09 | 6.01E-06 |
| FBgn0062412 | -1.09 | 7.89E-06 |
| FBgn0052250 | -1.09 | 8.95E-06 |
| FBgn0259238 | -1.09 | 0.00010076 |
| FBgn0263200 | -1.10 | 1.03E-05 |
| FBgn0034113 | -1.10 | 1.76E-05 |
| FBgn0040323 | -1.10 | 4.20E-05 |
| FBgn0037074 | -1.10 | 1.67E-08 |
| FBgn0036726 | -1.10 | 8.22E-06 |
| FBgn0261258 | -1.10 | 0.00011361 |
| FBgn0016684 | -1.11 | 1.93E-05 |
| FBgn0263490 | -1.11 | 1.26E-06 |
| FBgn0000116 | -1.11 | 3.76E-07 |
| FBgn0050104 | -1.11 | 0.00012188 |
| FBgn0036336 | -1.11 | 5.42E-05 |
| FBgn0030792 | -1.11 | 8.97E-06 |
| FBgn0035046 | -1.11 | 1.36E-05 |
| FBgn0010548 | -1.12 | 7.07E-14 |
| FBgn0031478 | -1.12 | 8.68E-08 |
| FBgn0263397 | -1.12 | 0.00013637 |
| FBgn0015372 | -1.12 | 1.21E-06 |
| FBgn0033366 | -1.12 | 1.28E-21 |
| FBgn0030482 | -1.12 | 7.95E-05 |
| FBgn0028970 | -1.12 | 3.61E-05 |
| FBgn0029849 | -1.12 | 1.45E-05 |
| FBgn0005278 | -1.12 | 7.62E-05 |
| FBgn0038795 | -1.12 | 3.64E-07 |
| FBgn0264389 | -1.13 | 3.01E-06 |
| FBgn0019957 | -1.13 | 5.74E-10 |
| FBgn0030292 | -1.13 | 2.59E-05 |
| FBgn0003189 | -1.13 | 9.83E-06 |
| FBgn0030245 | -1.13 | 1.21E-21 |
| FBgn0034432 | -1.13 | 7.61E-09 |
| FBgn0033735 | -1.13 | 1.79E-06 |
| FBgn0035911 | -1.13 | 2.05E-06 |
| FBgn0039741 | -1.14 | 2.14E-06 |
| FBgn0266994 | -1.14 | 9.92E-05 |
| FBgn0027596 | -1.14 | 6.05E-17 |
| FBgn0039151 | -1.14 | 9.86E-14 |
| FBgn0052280 | -1.14 | 0.00010944 |
| FBgn0014002 | -1.14 | 3.17E-07 |
| FBgn0042083 | -1.14 | 1.50E-07 |
| FBgn0032511 | -1.14 | 3.12E-07 |
| FBgn0000244 | -1.14 | 3.26E-06 |
| FBgn0035392 | -1.14 | 5.51E-07 |
| FBgn0003961 | -1.14 | 1.75E-07 |
| FBgn0033465 | -1.15 | 4.14E-11 |
| FBgn0030425 | -1.15 | 3.22E-08 |
| FBgn0261283 | -1.15 | 1.01E-09 |
| FBgn0014340 | -1.15 | 6.12E-06 |
| FBgn0037110 | -1.15 | 5.61E-05 |
| FBgn0015623 | -1.15 | 3.98E-09 |
| FBgn0013733 | -1.15 | 1.16E-12 |
| FBgn0032603 | -1.15 | 4.41E-07 |
| FBgn0263607 | -1.16 | 6.32E-08 |
| FBgn0038570 | -1.16 | 6.37E-07 |
| FBgn0038571 | -1.16 | 6.37E-07 |
| FBgn0001989 | -1.16 | 3.94E-07 |
| FBgn0052040 | -1.16 | 5.08E-06 |
| FBgn0030603 | -1.16 | 8.54E-09 |
| FBgn0033949 | -1.16 | 9.78E-11 |
| FBgn0086906 | -1.16 | 1.31E-11 |
| FBgn0033717 | -1.16 | 1.13E-12 |
| FBgn0039464 | -1.16 | 6.37E-09 |
| FBgn0035169 | -1.16 | 9.02E-06 |
| FBgn0267661 | -1.17 | 9.95E-11 |
| FBgn0037242 | -1.17 | 5.86E-11 |
| FBgn0035806 | -1.17 | 1.20E-06 |
| FBgn0027835 | -1.17 | 3.30E-09 |
| FBgn0040513 | -1.17 | 3.44E-05 |
| FBgn0050503 | -1.17 | 3.44E-05 |

| | | |
|---|---|---|
| FBgn0032876 | -1.17 | 3.34E-09 |
| FBgn0037942 | -1.18 | 4.67E-07 |
| FBgn0027579 | -1.18 | 3.41E-25 |
| FBgn0010352 | -1.18 | 2.23E-08 |
| FBgn0037614 | -1.18 | 4.69E-05 |
| FBgn0036844 | -1.19 | 1.31E-05 |
| FBgn0261989 | -1.19 | 4.52E-11 |
| FBgn0027291 | -1.19 | 2.61E-10 |
| FBgn0266945 | -1.20 | 1.00E-06 |
| FBgn0031320 | -1.20 | 2.58E-05 |
| FBgn0037370 | -1.20 | 7.53E-17 |
| FBgn0027660 | -1.20 | 3.89E-07 |
| FBgn0010516 | -1.20 | 1.78E-08 |
| FBgn0037001 | -1.20 | 9.16E-09 |
| FBgn0040636 | -1.20 | 3.40E-05 |
| FBgn0039827 | -1.21 | 7.24E-05 |
| FBgn0037842 | -1.21 | 6.17E-07 |
| FBgn0037057 | -1.21 | 4.37E-07 |
| FBgn0038224 | -1.21 | 7.22E-06 |
| FBgn0032618 | -1.21 | 6.71E-05 |
| FBgn0085342 | -1.21 | 6.71E-05 |
| FBgn0034877 | -1.21 | 1.72E-05 |
| FBgn0021765 | -1.21 | 4.19E-09 |
| FBgn0004652 | -1.21 | 1.91E-08 |
| FBgn0262559 | -1.22 | 2.47E-08 |
| FBgn0029888 | -1.22 | 4.08E-06 |
| FBgn0039112 | -1.22 | 1.59E-08 |
| FBgn0010213 | -1.22 | 3.18E-13 |
| FBgn0035085 | -1.22 | 6.05E-06 |
| FBgn0035110 | -1.23 | 3.46E-06 |
| FBgn0050373 | -1.23 | 0.00010752 |
| FBgn0263782 | -1.23 | 1.13E-06 |
| FBgn0031260 | -1.23 | 4.01E-09 |
| FBgn0019624 | -1.24 | 1.99E-06 |
| FBgn0033065 | -1.24 | 5.86E-05 |
| FBgn0039697 | -1.24 | 1.13E-07 |
| FBgn0011455 | -1.24 | 8.79E-08 |
| FBgn0041342 | -1.24 | 8.97E-06 |
| FBgn0038400 | -1.24 | 1.21E-07 |
| FBgn0029975 | -1.24 | 1.69E-06 |
| FBgn0086368 | -1.24 | 2.29E-05 |
| FBgn0030485 | -1.25 | 0.0001134 |
| FBgn0039051 | -1.25 | 5.96E-11 |
| FBgn0037440 | -1.25 | 3.37E-07 |
| FBgn0034141 | -1.25 | 3.29E-12 |
| FBgn0038271 | -1.26 | 1.22E-09 |
| FBgn0019644 | -1.26 | 2.28E-07 |
| FBgn0003462 | -1.26 | 4.12E-07 |
| FBgn0026878 | -1.26 | 5.81E-11 |
| FBgn0033247 | -1.26 | 5.64E-11 |
| FBgn0003965 | -1.26 | 3.18E-07 |
| FBgn0039909 | -1.26 | 2.56E-19 |
| FBgn0031069 | -1.26 | 3.49E-08 |
| FBgn0032833 | -1.26 | 7.91E-07 |
| FBgn0032770 | -1.27 | 6.48E-05 |
| FBgn0010612 | -1.27 | 7.89E-06 |
| FBgn0035791 | -1.27 | 2.49E-06 |
| FBgn0030478 | -1.27 | 1.25E-09 |
| FBgn0267605 | -1.27 | 2.60E-05 |
| FBgn0039188 | -1.28 | 1.88E-06 |
| FBgn0019830 | -1.28 | 1.10E-12 |
| FBgn0038407 | -1.28 | 3.20E-09 |
| FBgn0015582 | -1.28 | 1.38E-05 |
| FBgn0050410 | -1.28 | 8.61E-07 |
| FBgn0036762 | -1.28 | 1.67E-10 |
| FBgn0031066 | -1.28 | 1.79E-06 |
| FBgn0035600 | -1.28 | 4.19E-07 |
| FBgn0038830 | -1.28 | 2.36E-11 |
| FBgn0054043 | -1.28 | 2.05E-06 |
| FBgn0085263 | -1.28 | 5.58E-05 |
| FBgn0037621 | -1.28 | 3.34E-09 |
| FBgn0035356 | -1.28 | 1.52E-07 |
| FBgn0025620 | -1.29 | 1.02E-06 |
| FBgn0016120 | -1.29 | 2.82E-06 |
| FBgn0031703 | -1.29 | 5.11E-13 |
| FBgn0017566 | -1.30 | 4.95E-10 |
| FBgn0250906 | -1.30 | 3.24E-07 |
| FBgn0030731 | -1.30 | 8.23E-16 |
| FBgn0036007 | -1.30 | 3.65E-10 |
| FBgn0061359 | -1.30 | 3.51E-13 |
| FBgn0061360 | -1.30 | 3.51E-13 |
| FBgn0004598 | -1.30 | 1.36E-07 |
| FBgn0010053 | -1.30 | 2.07E-05 |
| FBgn0030607 | -1.30 | 4.44E-06 |
| FBgn0019643 | -1.30 | 1.21E-06 |
| FBgn0001092 | -1.30 | 4.93E-09 |
| FBgn0034733 | -1.31 | 3.50E-07 |
| FBgn0263278 | -1.31 | 7.06E-07 |
| FBgn0036551 | -1.31 | 6.25E-06 |
| FBgn0266411 | -1.31 | 8.13E-32 |

| | | |
|---|---|---|
| FBgn0250822 | -1.31 | 0.00011357 |
| FBgn0038312 | -1.31 | 8.98E-10 |
| FBgn0035084 | -1.32 | 2.70E-05 |
| FBgn0028916 | -1.32 | 7.89E-07 |
| FBgn0086355 | -1.32 | 9.33E-07 |
| FBgn0265873 | -1.33 | 8.58E-08 |
| FBgn0034075 | -1.33 | 6.38E-10 |
| FBgn0035252 | -1.33 | 1.38E-08 |
| FBgn0031117 | -1.33 | 2.94E-11 |
| FBgn0051198 | -1.33 | 8.07E-05 |
| FBgn0013764 | -1.33 | 2.46E-07 |
| FBgn0038865 | -1.34 | 3.06E-09 |
| FBgn0250838 | -1.34 | 1.94E-05 |
| FBgn0051915 | -1.34 | 9.60E-34 |
| FBgn0032219 | -1.35 | 1.68E-08 |
| FBgn0031170 | -1.35 | 7.51E-07 |
| FBgn0040336 | -1.35 | 1.27E-07 |
| FBgn0261808 | -1.35 | 5.69E-16 |
| FBgn0064115 | -1.36 | 4.15E-08 |
| FBgn0004407 | -1.36 | 4.15E-08 |
| FBgn0037537 | -1.36 | 2.98E-12 |
| FBgn0016691 | -1.36 | 4.19E-08 |
| FBgn0016122 | -1.36 | 2.78E-19 |
| FBgn0036334 | -1.36 | 2.84E-06 |
| FBgn0031830 | -1.37 | 8.70E-06 |
| FBgn0002921 | -1.37 | 1.00E-28 |
| FBgn0038804 | -1.37 | 5.02E-06 |
| FBgn0027580 | -1.37 | 5.05E-08 |
| FBgn0263911 | -1.38 | 5.58E-05 |
| FBgn0266637 | -1.38 | 1.35E-09 |
| FBgn0039674 | -1.39 | 7.11E-08 |
| FBgn0003890 | -1.39 | 3.89E-07 |
| FBgn0021906 | -1.39 | 2.21E-09 |
| FBgn0001250 | -1.39 | 7.42E-06 |
| FBgn0063923 | -1.39 | 9.10E-07 |
| FBgn0038974 | -1.40 | 1.23E-07 |
| FBgn0267725 | -1.40 | 6.19E-05 |
| FBgn0029648 | -1.41 | 9.45E-11 |
| FBgn0039431 | -1.41 | 1.09E-05 |
| FBgn0037891 | -1.41 | 4.34E-08 |
| FBgn0026259 | -1.41 | 1.70E-26 |
| FBgn0020235 | -1.42 | 2.92E-10 |
| FBgn0036273 | -1.42 | 1.09E-05 |
| FBgn0016687 | -1.42 | 1.31E-12 |
| FBgn0030668 | -1.42 | 4.24E-10 |
| FBgn0000579 | -1.42 | 8.41E-11 |
| FBgn0039152 | -1.43 | 3.82E-06 |
| FBgn0023537 | -1.43 | 2.34E-06 |
| FBgn0030872 | -1.43 | 2.17E-08 |
| FBgn0039635 | -1.43 | 4.74E-12 |
| FBgn0011754 | -1.43 | 9.29E-19 |
| FBgn0031010 | -1.43 | 3.26E-07 |
| FBgn0035674 | -1.43 | 4.82E-06 |
| FBgn0051248 | -1.43 | 1.45E-07 |
| FBgn0034716 | -1.44 | 5.33E-05 |
| FBgn0261439 | -1.44 | 1.15E-20 |
| FBgn0043806 | -1.44 | 3.84E-10 |
| FBgn0039157 | -1.44 | 2.90E-07 |
| FBgn0036211 | -1.44 | 1.76E-08 |
| FBgn0040507 | -1.45 | 7.51E-07 |
| FBgn0028940 | -1.45 | 1.12E-20 |
| FBgn0033112 | -1.46 | 2.47E-09 |
| FBgn0010217 | -1.46 | 4.63E-12 |
| FBgn0000150 | -1.46 | 2.83E-07 |
| FBgn0037134 | -1.46 | 6.24E-06 |
| FBgn0051674 | -1.46 | 4.65E-08 |
| FBgn0033883 | -1.47 | 5.32E-09 |
| FBgn0034628 | -1.47 | 1.64E-18 |
| FBgn0003360 | -1.47 | 1.58E-08 |
| FBgn0014869 | -1.48 | 4.10E-08 |
| FBgn0259714 | -1.48 | 1.97E-12 |
| FBgn0016119 | -1.48 | 1.63E-07 |
| FBgn0036728 | -1.48 | 1.61E-07 |
| FBgn0028342 | -1.48 | 8.62E-08 |
| FBgn0035032 | -1.48 | 4.02E-07 |
| FBgn0053054 | -1.48 | 7.83E-10 |
| FBgn0031662 | -1.48 | 7.34E-12 |
| FBgn0038337 | -1.49 | 2.90E-08 |
| FBgn0085244 | -1.49 | 2.59E-06 |
| FBgn0027348 | -1.49 | 1.31E-16 |
| FBgn0266582 | -1.49 | 2.14E-12 |
| FBgn0265191 | -1.49 | 5.28E-08 |
| FBgn0026634 | -1.50 | 5.82E-14 |
| FBgn0264712 | -1.50 | 4.28E-06 |
| FBgn0061356 | -1.50 | 3.22E-33 |
| FBgn0039564 | -1.50 | 2.79E-05 |
| FBgn0011204 | -1.51 | 2.42E-06 |
| FBgn0038524 | -1.51 | 5.18E-08 |
| FBgn0052699 | -1.51 | 1.85E-12 |
| FBgn0265703 | -1.52 | 8.20E-06 |

| | | |
|---|---|---|
| FBgn0026721 | -1.53 | 3.24E-16 |
| FBgn0032021 | -1.54 | 4.59E-08 |
| FBgn0004797 | -1.55 | 2.47E-06 |
| FBgn0037788 | -1.55 | 7.19E-05 |
| FBgn0039798 | -1.55 | 1.49E-18 |
| FBgn0038928 | -1.55 | 1.95E-05 |
| FBgn0250814 | -1.56 | 7.45E-14 |
| FBgn0029971 | -1.56 | 4.64E-07 |
| FBgn0266490 | -1.56 | 4.64E-07 |
| FBgn0039830 | -1.56 | 3.35E-11 |
| FBgn0037814 | -1.56 | 1.33E-10 |
| FBgn0025469 | -1.56 | 3.24E-11 |
| FBgn0261446 | -1.57 | 1.45E-06 |
| FBgn0033232 | -1.57 | 1.13E-21 |
| FBgn0042138 | -1.57 | 3.10E-17 |
| FBgn0001248 | -1.57 | 3.45E-06 |
| FBgn0015558 | -1.57 | 9.56E-08 |
| FBgn0033782 | -1.58 | 5.37E-17 |
| FBgn0050055 | -1.58 | 4.09E-05 |
| FBgn0042112 | -1.59 | 7.84E-06 |
| FBgn0032889 | -1.60 | 3.92E-08 |
| FBgn0031449 | -1.60 | 9.91E-08 |
| FBgn0053506 | -1.60 | 2.26E-09 |
| FBgn0030051 | -1.60 | 2.80E-12 |
| FBgn0020626 | -1.60 | 2.41E-16 |
| FBgn0000317 | -1.60 | 3.71E-19 |
| FBgn0040529 | -1.60 | 3.71E-11 |
| FBgn0053301 | -1.61 | 3.49E-05 |
| FBgn0053178 | -1.61 | 2.90E-06 |
| FBgn0263659 | -1.61 | 1.05E-10 |
| FBgn0034406 | -1.61 | 3.77E-11 |
| FBgn0259163 | -1.61 | 1.76E-08 |
| FBgn0033570 | -1.62 | 7.79E-13 |
| FBgn0044419 | -1.62 | 1.47E-13 |
| FBgn0260458 | -1.63 | 6.60E-14 |
| FBgn0037127 | -1.63 | 1.62E-09 |
| FBgn0032394 | -1.63 | 7.74E-16 |
| FBgn0035132 | -1.63 | 6.90E-15 |
| FBgn0052251 | -1.63 | 0.00012592 |
| FBgn0017482 | -1.64 | 2.19E-09 |
| FBgn0266275 | -1.64 | 1.36E-06 |
| FBgn0032801 | -1.65 | 5.98E-10 |
| FBgn0001970 | -1.66 | 1.72E-12 |
| FBgn0033742 | -1.66 | 8.46E-09 |
| FBgn0036039 | -1.66 | 8.88E-16 |
| FBgn0032782 | -1.66 | 6.42E-07 |
| FBgn0266428 | -1.66 | 9.81E-13 |
| FBgn0037229 | -1.66 | 7.74E-05 |
| FBgn0284257 | -1.67 | 9.35E-27 |
| FBgn0027521 | -1.67 | 2.25E-10 |
| FBgn0015569 | -1.68 | 1.24E-07 |
| FBgn0013469 | -1.69 | 3.43E-05 |
| FBgn0020236 | -1.69 | 1.32E-26 |
| FBgn0034712 | -1.69 | 4.16E-07 |
| FBgn0016053 | -1.69 | 2.32E-09 |
| FBgn0031418 | -1.69 | 2.17E-06 |
| FBgn0031818 | -1.70 | 9.93E-18 |
| FBgn0016693 | -1.70 | 2.82E-34 |
| FBgn0263120 | -1.70 | 1.57E-13 |
| FBgn0262808 | -1.70 | 1.84E-06 |
| FBgn0011211 | -1.71 | 4.12E-14 |
| FBgn0037356 | -1.71 | 1.21E-13 |
| FBgn0283494 | -1.72 | 2.98E-10 |
| FBgn0266889 | -1.72 | 2.96E-10 |
| FBgn0030737 | -1.72 | 9.32E-13 |
| FBgn0267041 | -1.72 | 7.51E-13 |
| FBgn0016075 | -1.72 | 1.63E-15 |
| FBgn0267814 | -1.72 | 0.00012842 |
| FBgn0026409 | -1.73 | 1.50E-19 |
| FBgn0034493 | -1.73 | 3.77E-05 |
| FBgn0036927 | -1.73 | 1.67E-09 |
| FBgn0029765 | -1.73 | 2.63E-05 |
| FBgn0023477 | -1.74 | 2.78E-09 |
| FBgn0040606 | -1.74 | 8.43E-06 |
| FBgn0001124 | -1.74 | 1.75E-35 |
| FBgn0028563 | -1.75 | 2.33E-10 |
| FBgn0029114 | -1.75 | 8.23E-10 |
| FBgn0013308 | -1.75 | 1.59E-12 |
| FBgn0019982 | -1.75 | 3.08E-12 |
| FBgn0032715 | -1.75 | 1.59E-19 |
| FBgn0085488 | -1.76 | 1.19E-10 |
| FBgn0031500 | -1.77 | 2.22E-14 |
| FBgn0262683 | -1.77 | 1.90E-22 |
| FBgn0063924 | -1.78 | 9.89E-06 |
| FBgn0038256 | -1.78 | 1.89E-26 |
| FBgn0035298 | -1.78 | 6.09E-13 |
| FBgn0260753 | -1.78 | 7.09E-06 |
| FBgn0003515 | -1.79 | 8.45E-05 |
| FBgn0086687 | -1.79 | 2.12E-11 |
| FBgn0039108 | -1.79 | 8.18E-11 |

| | | |
|---|---|---|
| FBgn0030593 | -1.80 | 3.52E-17 |
| FBgn0021995 | -1.80 | 7.58E-20 |
| FBgn0028325 | -1.80 | 5.43E-13 |
| FBgn0031561 | -1.80 | 3.51E-05 |
| FBgn0036298 | -1.81 | 1.06E-32 |
| FBgn0037213 | -1.82 | 8.36E-11 |
| FBgn0038290 | -1.83 | 1.58E-10 |
| FBgn0267385 | -1.84 | 2.63E-14 |
| FBgn0000299 | -1.84 | 1.95E-17 |
| FBgn0038179 | -1.85 | 1.71E-07 |
| FBgn0265872 | -1.85 | 7.55E-06 |
| FBgn0065084 | -1.85 | 0.00011686 |
| FBgn0085377 | -1.85 | 1.35E-12 |
| FBgn0025814 | -1.85 | 1.34E-13 |
| FBgn0039016 | -1.86 | 2.64E-26 |
| FBgn0267804 | -1.86 | 2.15E-12 |
| FBgn0029791 | -1.86 | 9.03E-27 |
| FBgn0266384 | -1.86 | 6.75E-16 |
| FBgn0265613 | -1.87 | 6.53E-13 |
| FBgn0030484 | -1.88 | 2.60E-20 |
| FBgn0031695 | -1.88 | 5.62E-12 |
| FBgn0036822 | -1.89 | 7.29E-06 |
| FBgn0029854 | -1.90 | 8.60E-09 |
| FBgn0260960 | -1.90 | 1.87E-14 |
| FBgn0029769 | -1.90 | 3.94E-05 |
| FBgn0041581 | -1.91 | 1.55E-05 |
| FBgn0029969 | -1.91 | 1.15E-10 |
| FBgn0266064 | -1.92 | 1.55E-12 |
| FBgn0063491 | -1.92 | 1.31E-11 |
| FBgn0050090 | -1.93 | 4.64E-08 |
| FBgn0035839 | -1.93 | 1.05E-17 |
| FBgn0026150 | -1.93 | 2.43E-21 |
| FBgn0037709 | -1.94 | 4.62E-10 |
| FBgn0035026 | -1.94 | 6.04E-31 |
| FBgn0000406 | -1.94 | 1.08E-08 |
| FBgn0034046 | -1.94 | 8.47E-18 |
| FBgn0032913 | -1.94 | 8.46E-17 |
| FBgn0260858 | -1.94 | 2.30E-32 |
| FBgn0053082 | -1.95 | 4.99E-12 |
| FBgn0014903 | -1.96 | 5.25E-12 |
| FBgn0032629 | -1.96 | 4.90E-06 |
| FBgn0038332 | -1.96 | 2.46E-15 |
| FBgn0031012 | -1.96 | 3.66E-09 |
| FBgn0025383 | -1.96 | 6.95E-06 |
| FBgn0015570 | -1.96 | 7.53E-15 |
| FBgn0013772 | -1.96 | 7.66E-07 |
| FBgn0030701 | -1.97 | 2.07E-10 |
| FBgn0037607 | -1.97 | 3.19E-11 |
| FBgn0001075 | -1.97 | 8.28E-05 |
| FBgn0005666 | -1.98 | 1.00E-09 |
| FBgn0017448 | -1.98 | 7.76E-10 |
| FBgn0022709 | -1.98 | 3.98E-15 |
| FBgn0033101 | -1.99 | 6.30E-24 |
| FBgn0035517 | -1.99 | 6.88E-25 |
| FBgn0022359 | -1.99 | 1.49E-18 |
| FBgn0002719 | -2.00 | 1.13E-10 |
| FBgn0039189 | -2.00 | 4.91E-06 |
| FBgn0038858 | -2.01 | 1.14E-09 |
| FBgn0051373 | -2.02 | 2.63E-12 |
| FBgn0043783 | -2.02 | 2.50E-08 |
| FBgn0030966 | -2.02 | 2.56E-17 |
| FBgn0267617 | -2.03 | 4.70E-08 |
| FBgn0263999 | -2.03 | 4.15E-08 |
| FBgn0029994 | -2.03 | 2.84E-21 |
| FBgn0016123 | -2.04 | 6.98E-07 |
| FBgn0270926 | -2.04 | 1.51E-12 |
| FBgn0038149 | -2.04 | 1.51E-14 |
| FBgn0051548 | -2.04 | 1.43E-13 |
| FBgn0040256 | -2.05 | 4.25E-07 |
| FBgn0038326 | -2.05 | 1.37E-19 |
| FBgn0034045 | -2.05 | 1.73E-12 |
| FBgn0039737 | -2.05 | 2.02E-13 |
| FBgn0039519 | -2.05 | 9.90E-10 |
| FBgn0003067 | -2.06 | 4.14E-10 |
| FBgn0266488 | -2.06 | 4.14E-10 |
| FBgn0034736 | -2.06 | 1.23E-26 |
| FBgn0266705 | -2.06 | 1.03E-19 |
| FBgn0001128 | -2.06 | 1.91E-09 |
| FBgn0264507 | -2.07 | 1.41E-09 |
| FBgn0036824 | -2.07 | 1.69E-08 |
| FBgn0034710 | -2.07 | 3.03E-07 |
| FBgn0039494 | -2.08 | 2.56E-27 |
| FBgn0050485 | -2.08 | 4.39E-14 |
| FBgn0041605 | -2.08 | 4.48E-14 |
| FBgn0003074 | -2.09 | 8.03E-13 |
| FBgn0263087 | -2.10 | 2.61E-06 |
| FBgn0051313 | -2.11 | 2.26E-11 |
| FBgn0025631 | -2.11 | 1.69E-21 |
| FBgn0025373 | -2.12 | 9.06E-22 |
| FBgn0024957 | -2.12 | 7.89E-11 |

| | | |
|---|---|---|
| FBgn0052446 | -2.13 | 6.71E-35 |
| FBgn0003313 | -2.13 | 2.64E-06 |
| FBgn0038039 | -2.14 | 4.10E-38 |
| FBgn0027866 | -2.15 | 1.27E-24 |
| FBgn0005636 | -2.16 | 1.87E-09 |
| FBgn0035147 | -2.17 | 1.80E-18 |
| FBgn0026593 | -2.18 | 6.94E-13 |
| FBgn0024293 | -2.18 | 2.39E-28 |
| FBgn0028573 | -2.19 | 2.30E-10 |
| FBgn0038662 | -2.20 | 1.05E-21 |
| FBgn0051287 | -2.20 | 2.86E-07 |
| FBgn0052391 | -2.20 | 2.30E-07 |
| FBgn0038610 | -2.20 | 3.79E-06 |
| FBgn0030160 | -2.21 | 3.65E-23 |
| FBgn0026314 | -2.22 | 8.72E-05 |
| FBgn0034198 | -2.22 | 1.97E-06 |
| FBgn0039223 | -2.23 | 1.89E-17 |
| FBgn0039609 | -2.24 | 1.39E-07 |
| FBgn0004654 | -2.24 | 1.95E-16 |
| FBgn0085433 | -2.24 | 9.14E-11 |
| FBgn0010504 | -2.24 | 5.19E-11 |
| FBgn0033879 | -2.25 | 4.67E-16 |
| FBgn0041712 | -2.25 | 2.25E-11 |
| FBgn0032218 | -2.25 | 9.19E-31 |
| FBgn0016920 | -2.26 | 0.00011708 |
| FBgn0259243 | -2.27 | 7.03E-32 |
| FBgn0001187 | -2.28 | 2.92E-06 |
| FBgn0023507 | -2.29 | 4.91E-17 |
| FBgn0069056 | -2.30 | 2.64E-14 |
| FBgn0031801 | -2.31 | 0.00010565 |
| FBgn0039538 | -2.31 | 3.38E-08 |
| FBgn0039094 | -2.31 | 2.65E-13 |
| FBgn0031974 | -2.31 | 2.31E-13 |
| FBgn0039156 | -2.32 | 7.49E-15 |
| FBgn0085359 | -2.32 | 1.31E-06 |
| FBgn0265705 | -2.32 | 3.85E-10 |
| FBgn0035734 | -2.32 | 2.63E-12 |
| FBgn0016694 | -2.33 | 4.35E-16 |
| FBgn0066101 | -2.34 | 5.60E-13 |
| FBgn0033830 | -2.35 | 8.27E-11 |
| FBgn0261564 | -2.35 | 2.15E-50 |
| FBgn0038682 | -2.35 | 9.38E-05 |
| FBgn0039052 | -2.36 | 1.95E-06 |
| FBgn0030326 | -2.37 | 2.14E-19 |
| FBgn0051472 | -2.38 | 4.91E-32 |
| FBgn0034479 | -2.38 | 7.55E-10 |
| FBgn0265045 | -2.38 | 1.28E-41 |
| FBgn0267452 | -2.39 | 1.45E-05 |
| FBgn0037230 | -2.39 | 4.57E-14 |
| FBgn0023513 | -2.40 | 7.25E-29 |
| FBgn0034638 | -2.41 | 4.99E-18 |
| FBgn0040813 | -2.41 | 3.13E-14 |
| FBgn0000592 | -2.41 | 1.60E-23 |
| FBgn0015777 | -2.42 | 1.14E-29 |
| FBgn0005660 | -2.42 | 2.01E-44 |
| FBgn0030331 | -2.43 | 7.76E-05 |
| FBgn0020764 | -2.44 | 4.79E-21 |
| FBgn0029091 | -2.45 | 1.66E-17 |
| FBgn0010043 | -2.45 | 2.41E-11 |
| FBgn0052311 | -2.45 | 2.74E-80 |
| FBgn0031261 | -2.45 | 8.65E-45 |
| FBgn0042627 | -2.46 | 5.97E-11 |
| FBgn0038105 | -2.47 | 1.51E-12 |
| FBgn0033519 | -2.49 | 0.0001142 |
| FBgn0267220 | -2.50 | 3.29E-19 |
| FBgn0037890 | -2.50 | 2.05E-13 |
| FBgn0038331 | -2.50 | 1.33E-53 |
| FBgn0039307 | -2.53 | 1.12E-06 |
| FBgn0031064 | -2.53 | 0.00011979 |
| FBgn0036353 | -2.54 | 6.80E-36 |
| FBgn0034440 | -2.54 | 2.68E-10 |
| FBgn0011828 | -2.55 | 1.88E-67 |
| FBgn0283427 | -2.55 | 8.36E-17 |
| FBgn0265767 | -2.57 | 1.42E-27 |
| FBgn0027610 | -2.58 | 1.25E-47 |
| FBgn0035582 | -2.59 | 7.62E-05 |
| FBgn0034885 | -2.60 | 5.23E-18 |
| FBgn0004057 | -2.60 | 7.52E-31 |
| FBgn0043575 | -2.61 | 1.33E-13 |
| FBgn0051664 | -2.61 | 1.42E-22 |
| FBgn0038846 | -2.61 | 4.21E-05 |
| FBgn0266446 | -2.62 | 1.37E-07 |
| FBgn0261955 | -2.64 | 1.13E-29 |
| FBgn0036205 | -2.67 | 4.17E-35 |
| FBgn0026315 | -2.67 | 1.32E-69 |
| FBgn0010387 | -2.68 | 1.15E-09 |
| FBgn0032891 | -2.69 | 8.40E-11 |
| FBgn0085399 | -2.70 | 9.38E-05 |
| FBgn0037167 | -2.72 | 2.21E-36 |
| FBgn0038242 | -2.73 | 2.29E-06 |

| | | |
|---|---|---|
| FBgn0035083 | -2.74 | 2.64E-19 |
| FBgn0000473 | -2.75 | 9.13E-09 |
| FBgn0023001 | -2.77 | 1.02E-30 |
| FBgn0012034 | -2.77 | 1.13E-14 |
| FBgn0065097 | -2.81 | 6.22E-20 |
| FBgn0015766 | -2.83 | 4.63E-25 |
| FBgn0022160 | -2.84 | 7.53E-21 |
| FBgn0053056 | -2.88 | 8.90E-39 |
| FBgn0259237 | -2.88 | 2.08E-05 |
| FBgn0019929 | -2.88 | 4.48E-28 |
| FBgn0264776 | -2.90 | 3.13E-12 |
| FBgn0036749 | -2.90 | 4.95E-29 |
| FBgn0039324 | -2.91 | 1.27E-05 |
| FBgn0030816 | -2.92 | 2.85E-22 |
| FBgn0025720 | -2.93 | 4.78E-98 |
| FBgn0030073 | -2.93 | 4.37E-28 |
| FBgn0033179 | -2.95 | 9.58E-49 |
| FBgn0033128 | -2.95 | 4.39E-27 |
| FBgn0029518 | -2.96 | 4.83E-08 |
| FBgn0037714 | -2.97 | 1.42E-12 |
| FBgn0025885 | -2.98 | 3.33E-33 |
| FBgn0041150 | -2.99 | 1.68E-08 |
| FBgn0264384 | -2.99 | 4.21E-25 |
| FBgn0026718 | -3.00 | 9.06E-38 |
| FBgn0012042 | -3.03 | 1.02E-05 |
| FBgn0010549 | -3.07 | 1.93E-06 |
| FBgn0038257 | -3.08 | 1.07E-14 |
| FBgn0037163 | -3.16 | 1.49E-32 |
| FBgn0002773 | -3.16 | 2.29E-13 |
| FBgn0051559 | -3.19 | 0.00013358 |
| FBgn0034279 | -3.25 | 2.42E-12 |
| FBgn0030612 | -3.26 | 9.70E-40 |
| FBgn0033397 | -3.29 | 3.24E-40 |
| FBgn0265981 | -3.29 | 0.00012361 |
| FBgn0030817 | -3.30 | 1.80E-25 |
| FBgn0034490 | -3.36 | 3.25E-09 |
| FBgn0266753 | -3.36 | 8.76E-05 |
| FBgn0020372 | -3.37 | 6.44E-25 |
| FBgn0038730 | -3.47 | 1.17E-07 |
| FBgn0037715 | -3.47 | 8.37E-20 |
| FBgn0035880 | -3.48 | 5.35E-12 |
| FBgn0267035 | -3.52 | 5.03E-33 |
| FBgn0032178 | -3.56 | 5.05E-08 |
| FBgn0053474 | -3.56 | 3.41E-14 |
| FBgn0053138 | -3.59 | 3.78E-31 |
| FBgn0037126 | -3.61 | 1.38E-06 |
| FBgn0023541 | -3.63 | 1.20E-36 |
| FBgn0033395 | -3.66 | 7.25E-06 |
| FBgn0261053 | -3.66 | 3.29E-13 |
| FBgn0038095 | -3.74 | 9.07E-44 |
| FBgn0010482 | -3.78 | 3.75E-12 |
| FBgn0034390 | -3.81 | 1.36E-47 |
| FBgn0264679 | -3.81 | 1.85E-48 |
| FBgn0033307 | -3.83 | 5.05E-06 |
| FBgn0085223 | -3.87 | 2.16E-51 |
| FBgn0004426 | -3.88 | 4.02E-06 |
| FBgn0034725 | -3.92 | 1.81E-06 |
| FBgn0035241 | -3.97 | 2.40E-19 |
| FBgn0034724 | -4.05 | 9.16E-16 |
| FBgn0035770 | -4.06 | 5.41E-148 |
| FBgn0001090 | -4.10 | 5.11E-73 |
| FBgn0050374 | -4.15 | 1.04E-12 |
| FBgn0264877 | -4.29 | 9.99E-15 |
| FBgn0037690 | -4.41 | 2.67E-08 |
| FBgn0017561 | -4.46 | 2.35E-35 |
| FBgn0033129 | -4.50 | 8.35E-49 |
| FBgn0035673 | -4.51 | 1.49E-79 |
| FBgn0033246 | -4.61 | 1.01E-50 |
| FBgn0266402 | -4.66 | 9.45E-09 |
| FBgn0052255 | -4.70 | 1.01E-15 |
| FBgn0004427 | -4.76 | 8.57E-10 |
| FBgn0050457 | -4.77 | 1.96E-10 |
| FBgn0033079 | -4.80 | 3.10E-82 |
| FBgn0026061 | -4.84 | 7.41E-205 |
| FBgn0039486 | -4.85 | 4.75E-17 |
| FBgn0034204 | -4.88 | 5.25E-13 |
| FBgn0027106 | -5.00 | 8.57E-148 |
| FBgn0038731 | -5.02 | 3.01E-17 |
| FBgn0266591 | -5.13 | 1.27E-12 |
| FBgn0266430 | -5.15 | 2.40E-54 |
| FBgn0264866 | -5.17 | 5.44E-11 |
| FBgn0267441 | -5.17 | 5.44E-11 |
| FBgn0085428 | -5.42 | 1.13E-43 |
| FBgn0051427 | -5.51 | 4.13E-24 |
| FBgn0045478 | -5.65 | 1.15E-15 |
| FBgn0045477 | -5.65 | 1.15E-15 |
| FBgn0031860 | -6.07 | 6.24E-72 |
| FBgn0045476 | -6.08 | 5.60E-20 |
| FBgn0035486 | -6.12 | 3.36E-20 |
| FBgn0262593 | -7.54 | 5.45E-71 |

| | | |
|---|---|---|
| FBgn0033271 | -8.40 | 5.60E-55 |
